# Predicting taxon-specific benthic cyanobacterial mat cover and anatoxin concentrations in northern California rivers

**DOI:** 10.64898/2026.09.04.748937

**Authors:** Jordan M. Zabrecky, Taryn A. Elliott, Meaghan Hickey, Keith Bouma-Gregson, Gregory L. Boyer, Rich Fadness, Laurel Genzoli, Ramesh Goel, Grant Johnson, Robert Shriver, Rosalina Stancheva, Michael Thomas, Zac Triumph, Joanna R. Blaszczak

**Affiliations:** Department of Natural Resources and Environmental Science, University of Nevada, Reno, NV, USA; U.S. Geological Survey California Water Science Center, Sacramento, CA, USA; Department of Chemistry, State University of New York College of Environmental Science and Forestry, Syracuse, NY, USA; North Coast Regional Water Quality Control Board, Santa Rosa, CA, USA; Department of Civil and Environmental Engineering, University of Utah, Salt Lake City, UT, USA; Department of Natural Resources, Karuk Tribe, Orleans, CA, USA; Department of Environmental Science and Policy, George Mason University, Fairfax, VA, USA

**Keywords:** *Anabaena*, anatoxins, benthic cyanobacteria, cyanotoxins, ecological forecasting, gross primary productivity, *Microcoleus*, rivers, predictive models

## Abstract

I.

Ecological forecasts often rely on established relationships between the ecological process of interest and predictor variables that are more easily measured. Proliferations of benthic (i.e., bottom-dwelling) cyanobacteria have been increasingly observed in rivers globally and are an emerging ecological forecasting issue as they pose a public health threat due to the production of potent neurotoxins known as anatoxins. Controls on these benthic cyanobacteria are poorly understood, thus predicting or forecasting their extent and anatoxin production is a significant challenge. Here, we measured benthic cyanobacterial cover and anatoxin concentrations for two common taxa associated with anatoxins (*Microcoleus* and *Anabaena*) at biweekly to weekly intervals during June to September in 2022 and 2023 in three northern Californian rivers. We then built predictive models to test how incorporation of a biotic predictor (river reach-scale gross primary productivity [GPP]) affected predictive accuracy in addition to widely measured abiotic predictors (i.e., nutrients, discharge, and temperature). Temporal patterns in taxon-specific benthic cyanobacterial cover and anatoxin concentrations were highly variable among rivers and between taxa. While *Microcoleus* cover peaked in rivers during periods of relatively low GPP, there were no clear relationships between GPP and *Anabaena* cover nor either taxon’s anatoxin concentrations among rivers. Among multiple reaches sampled weekly in the South Fork Eel River, magnitudes of taxon-specific benthic cyanobacterial cover and anatoxin concentrations differed, but the timing of peak taxon-specific cover and anatoxin concentrations were generally consistent. Furthermore, *Anabaena* displayed a “hysteresis” relationship where increases in cover were followed by increases in anatoxin concentrations while *Microcoleus* lacked such relationship. While incorporating GPP as a covariate improved our predictions of *Anabaena* cover, we had more success predicting *Microcoleus* cover than *Anabaena* cover due to its strong negative relationship with discharge. In contrast, our models predicting *Anabaena* anatoxin concentrations outperformed those predicting *Microcoleus* anatoxin concentrations due to the “hysteresis” relationship between *Anabaena* cover and anatoxins. Overall, our predictive modeling results highlight the application of ecological forecasting for benthic cyanobacterial cover and anatoxin concentrations in rivers and demonstrate the importance of incorporating taxon-specific predictors into future forecasts of benthic cyanobacteria.

**Open Research Statement:** Data (Zabrecky et al. 2025) collected for this project are available at the Environmental Data Initiative at https://doi.org/10.6073/pasta/dd858a8903f9d22ec7f4ef03584f5bd5. Additional data were used from the U.S. Geological Survey Water Data for the Nation (U.S. Geological Survey 2025), Global Land Data Assimilation System (Rodell et al. 2024), North American Land Data Assimilation System (Xia et al. 2012), and from the Karuk Tribe with consent. The data from the Karuk Tribe are not currently publicly available. Contact the Tribe for further information. Code is currently publicly available at https://github.com/jzabrecky/ATX-synchrony-norcal and will be deposited on Zenodo upon article acceptance.

## II. Introduction

Forecasts of population dynamics (e.g., wildlife, insects, algae) can inform resource and recreation management decisions (Dietze et al. 2018, Carey et al. 2022, Lofton et al. 2023). Population forecasts that predict the future abundance of an organism with uncertainty can be generated from process models based on our understanding of organism-specific dynamics (e.g., Ricker model of fish abundances, Ricker 1954) or from statistical (also known as “data-driven”) models that use proxies to predict abundances (Rousso et al. 2020). While process-models may be more transferable to novel conditions (Yates et al. 2018), the creation of these models requires a strong understanding of an organism’s dynamics (e.g., temperature dependent growth rates, nutrient uptake rates, etc.). Thus, in situations where the mechanisms governing an organism are poorly understood, statistical models relying on correlative relationships may be necessary to forecast population dynamics.

In freshwater ecosystems, populations of toxin-producing cyanobacteria can pose risks to recreational users and drinking water resources (Huisman et al. 2018). In particular, toxigenic benthic (i.e., bottom-dwelling) cyanobacteria have been increasingly detected in rivers globally (Wood et al. 2020) and are associated with the production of secondary metabolites including anatoxins which are a group of neurotoxins that can harm humans and animals (Christensen and Khan 2020). The ingestion of anatoxins has resulted in dog deaths in recreational waters (Gugger et al. 2005, Puschner et al. 2008, Faassen et al. 2012, Wood et al. 2017b, McCarron et al. 2023, Fredrickson et al. 2023) and anatoxins have been found to be lethal to macroinvertebrates at environmentally relevant concentrations (Toporowska et al. 2014, Anderson et al. 2018). A common anatoxin congener, anatoxin-a, has been shown to degrade rapidly once released to the water column under a neutral or elevated pH (Kaminski et al. 2013). Thus, anatoxins are predominantly considered a concern if mat material is ingested. Therefore, to properly characterize and forecast variation in anatoxin concentrations within a river, the extent and toxicity of benthic cyanobacterial mats is important to understand.

The most well-studied anatoxin-producing benthic cyanobacterium found worldwide is *Microcoleus* (formerly known as *Phormidium*, Strunecký et al. 2013) which forms dark mats and has, at present, primarily been observed in rivers in shallower areas with faster flow (McAllister et al. 2016, Kelly et al. 2026). Other benthic cyanobacteria associated with anatoxins include *Anabaena* (Bouma-Gregson et al. 2018) and *Cylindrospermum* (Méjean et al. 2016) which are both nitrogen-fixing members in the order Nostocales and often form emerald green towers in slower moving areas of river channels. As these taxa can be difficult to distinguish, we have grouped them together in our study. Although there remains debate on whether *Anabaena* actually produce anatoxins or if it is another anatoxin-producing taxa (such as *Microcoleus* or *Geitlerinema*) co-occurring within their mats that produce anatoxins (Kust et al. 2018, Kelly et al. 2019), these mats can have high anatoxin concentrations (Bouma-Gregson et al. 2018) and therefore is important to include in assessments and forecasts of benthic cyanobacterial anatoxins.

Despite the increased awareness of toxigenic benthic cyanobacteria in rivers (Quiblier et al. 2013, McAllister et al. 2016, Wood et al. 2020, Kelly et al. 2026), they can be challenging to observe, quantify, and predict. Compared to their planktonic (i.e., free-floating) counterparts that are typically associated with lakes and reservoirs (Huisman et al. 2018, Schaeffer et al. 2022), benthic cyanobacteria, usually found in rivers or lake edges, cannot be easily detected by satellite due to riparian shading, surface turbulence, variable patch sizes, and their often similar color to that of their substrate surface. Monitoring or quantifying benthic cyanobacterial abundances can also be challenging in the field due to destructive sampling techniques, patchiness of biomass, and general difficulties that arise when working in flowing water. Furthermore, unlike their planktonic counterparts, benthic cyanobacteria proliferations do not necessarily occur due to eutrophication (i.e., excess nutrients; Wurtsbaugh et al. 2019). Benthic cyanobacteria can proliferate in low nutrient rivers (Loza et al. 2013, Genzoli et al. 2024, Sohrab et al. 2025, Diez-Chiappe et al. 2025), limiting the predictive power of nutrient-related variables for modeling benthic cyanobacterial biomass. As such, benthic cyanobacteria are infrequently modelled in comparison to planktonic cyanobacteria (Murphy et al. 2025).

Beyond forecasting benthic cyanobacterial mat cover, the prediction of anatoxins within benthic cyanobacterial mats is further complicated as anatoxin concentrations often do not necessarily scale with cover or biomass (McAllister et al. 2018a, Robichon et al. 2025) due to high spatiotemporal variability (Wood et al. 2010, 2012, Heath et al. 2011) and the coexistence of toxic and nontoxic genotypes (Wood et al. 2012). Ultimately, it has been suggested that a combination of abiotic (i.e., nutrients, flow) and biotic (i.e., grazing, density-dependence) factors and interactions influence the proliferation and toxin production of benthic cyanobacteria but their relative importance is poorly understood (Thomson-Laing et al. 2021). Therefore, finding proxies for benthic cyanobacterial population dynamics, such as ecosystem-level processes that are readily obtained with high-frequency sensor data, may provide a valuable way to predict or forecast these populations.

Gross primary productivity (GPP) is an ecosystem level process in rivers that can serve as an integrative metric of photoautotrophic activity, thereby providing insights into successional changes in benthic communities within rivers. Similar to benthic cyanobacteria, GPP is also controlled by a combination of abiotic (Bernhardt et al. 2022) controls such as light and flow and biotic controls such as density-dependent growth (Borlestean et al. 2015) and grazing (Vadeboncoeur and Power 2017). GPP can be estimated in rivers through diel patterns in dissolved oxygen (Odum 1956) and, with recent advances in sensor technology and modeling, can be relatively quickly and accurately estimated from high-frequency sensor data (Appling et al. 2018, Bernhardt et al. 2018). Thus, GPP can provide insights on the timing of autotrophic biomass production and accumulation in rivers (Blaszczak et al. 2023). Even though benthic cyanobacteria may not be the main contributors to GPP, evaluating relationships between the two may provide insight on the relative controls acting on each and GPP may serve as a more easily obtainable proxy to use when predicting or forecasting the timing of benthic cyanobacteria dynamics (referring to both extent of cover and anatoxin concentrations).

In this study, we evaluated patterns of benthic cyanobacterial dynamics and GPP to determine if GPP could be used to improve predictions of benthic cyanobacterial dynamics. We generated time series of benthic cyanobacterial mat cover through benthic cover surveys, collected composite mat samples of two benthic cyanobacterial taxa associated with anatoxin production (*Microcoleus* and *Anabaena*/*Cylindrospermum*) to analyze for anatoxin concentrations, and estimated GPP for two summers in three rivers in northern California with histories of benthic anatoxin detections (Bouma-Gregson et al. 2018, Conklin et al. 2020, NCRWQCB 2022, Genzoli et al. 2024). We then created a series of predictive models to test if the incorporation of GPP improved predictions of taxon-specific benthic cyanobacterial dynamics beyond abiotic predictors alone. Specifically, our research was guided by the following questions:

**Q1.** How do temporal patterns in benthic cyanobacterial cover, anatoxin concentrations, and gross primary productivity (GPP) covary across rivers?

**Q2.** How do relationships between benthic cyanobacterial cover and anatoxin concentrations covary among reaches within a river?

**Q3.** How do predictions of benthic cyanobacterial cover and anatoxin concentrations differ among taxa and how does the incorporation of GPP change these predictions in addition to using abiotic predictors (e.g., temperature, nutrients, discharge)?

If the abiotic and biotic controls acting on GPP and benthic cyanobacterial dynamics are similar, all processes could be expected to peak synchronously (Figure 1a). However, if the controls on benthic cyanobacteria dynamics differ from those on GPP, the two may peak asynchronously (Figure 1b). Alternatively, if anatoxin concentrations are not a function of benthic cyanobacterial cover, we may see benthic cyanobacterial dynamics peak asynchronously (Figure 1c). Further characterizing these relationships may inform whether GPP can be used as a predictor in forecasting benthic cyanobacteria cover and anatoxin concentrations.

**Figure 1.**
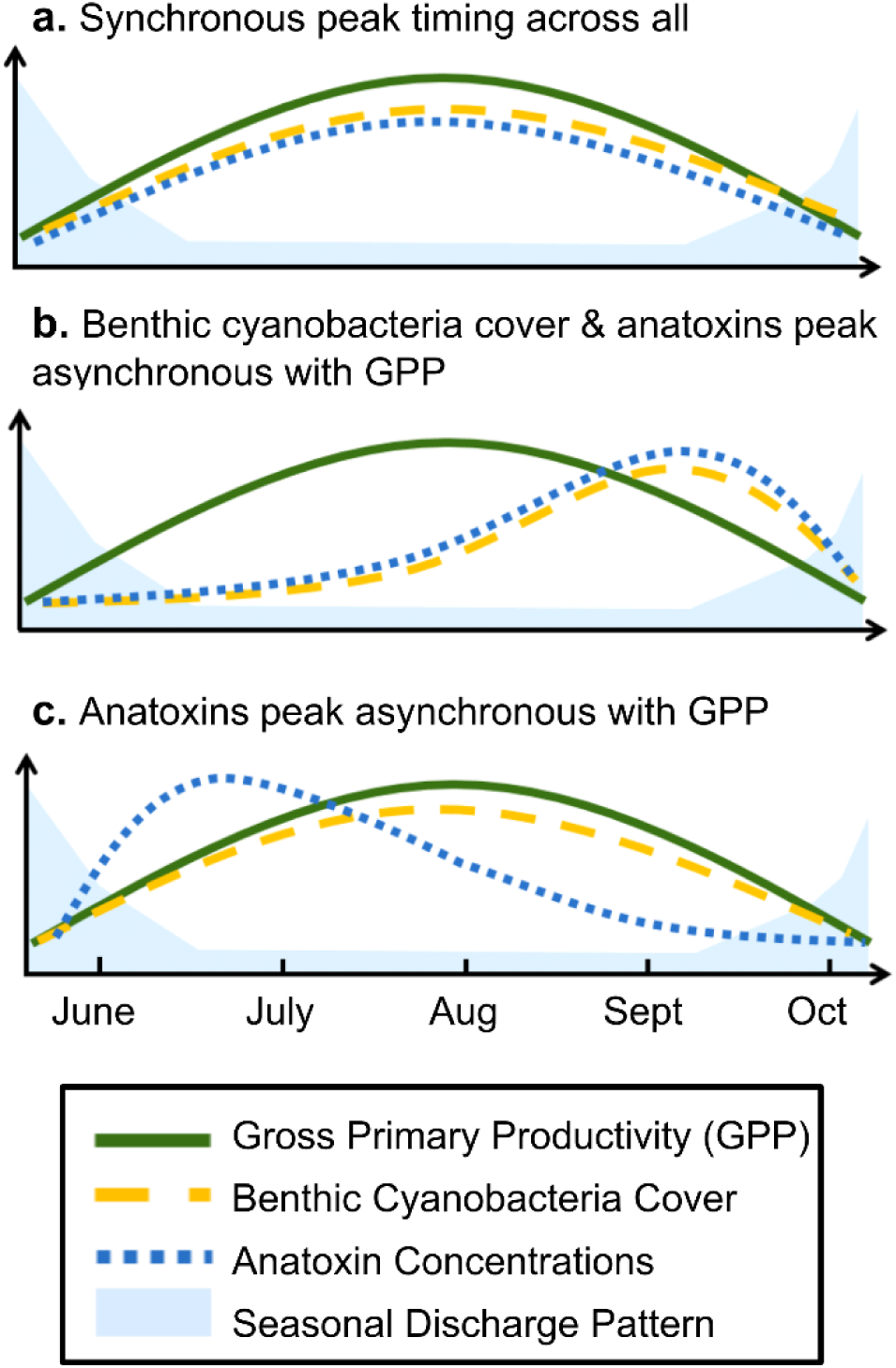
Conceptual diagram of three hypotheses regarding the temporal synchrony of gross primary productivity (GPP), benthic cyanobacterial cover, and anatoxin concentrations. (a) All three processes peak synchronously. (b) Benthic cyanobacterial cover and anatoxin concentrations peak asynchronously from GPP. (c) Anatoxin concentrations peak asynchronously from benthic cyanobacterial cover and GPP.

## Methods

### Site Description

We conducted our study from mid-June to late September in 2022 and 2023 in the South Fork Eel, Russian, and Salmon Rivers of northwestern California (Figure 2a) where prior anatoxin detections have occurred (Bouma-Gregson et al. 2018, Conklin et al. 2020, NCRWQCB 2022, Genzoli et al. 2024). All three rivers have a Mediterranean climate characterized by wet, cool winters and dry, hot summers (Power et al. 2024). As flows subside during the summer and fall, these rivers can host high biological activity (Power et al. 2009). At the location of sampling, each river was a 5^th^ order river by Strahler order classification in National Hydrography Dataset Plus version 2 (NHDPlus V2; McKay et al. 2012) and had roughly similar land cover and nutrient ranges (Appendix 1: Table S1). However, each river had different flow regimes during the study period (U.S. Geological Survey 2025). The South Fork Eel River is a non-regulated river that declined in discharge over the summer (median, range over period of study: 1.2, 0.4-4.2 m^3^ s^-1^ in 2022 and 1.1, 0.3-4.8 m^3^ s^-1^in 2023). In contrast, the Russian River is a regulated river which maintained a more consistent discharge over the summer of 2022 (1.6, 0.8-2.1 m^3^ s^-1^). Lastly, the Salmon River is an unregulated river with a median discharge over six times larger than that of the South Fork Eel River during our study period (6.9, 4.4-19.1 m^3^ s^-1^ in 2022).

**Figure 2.**
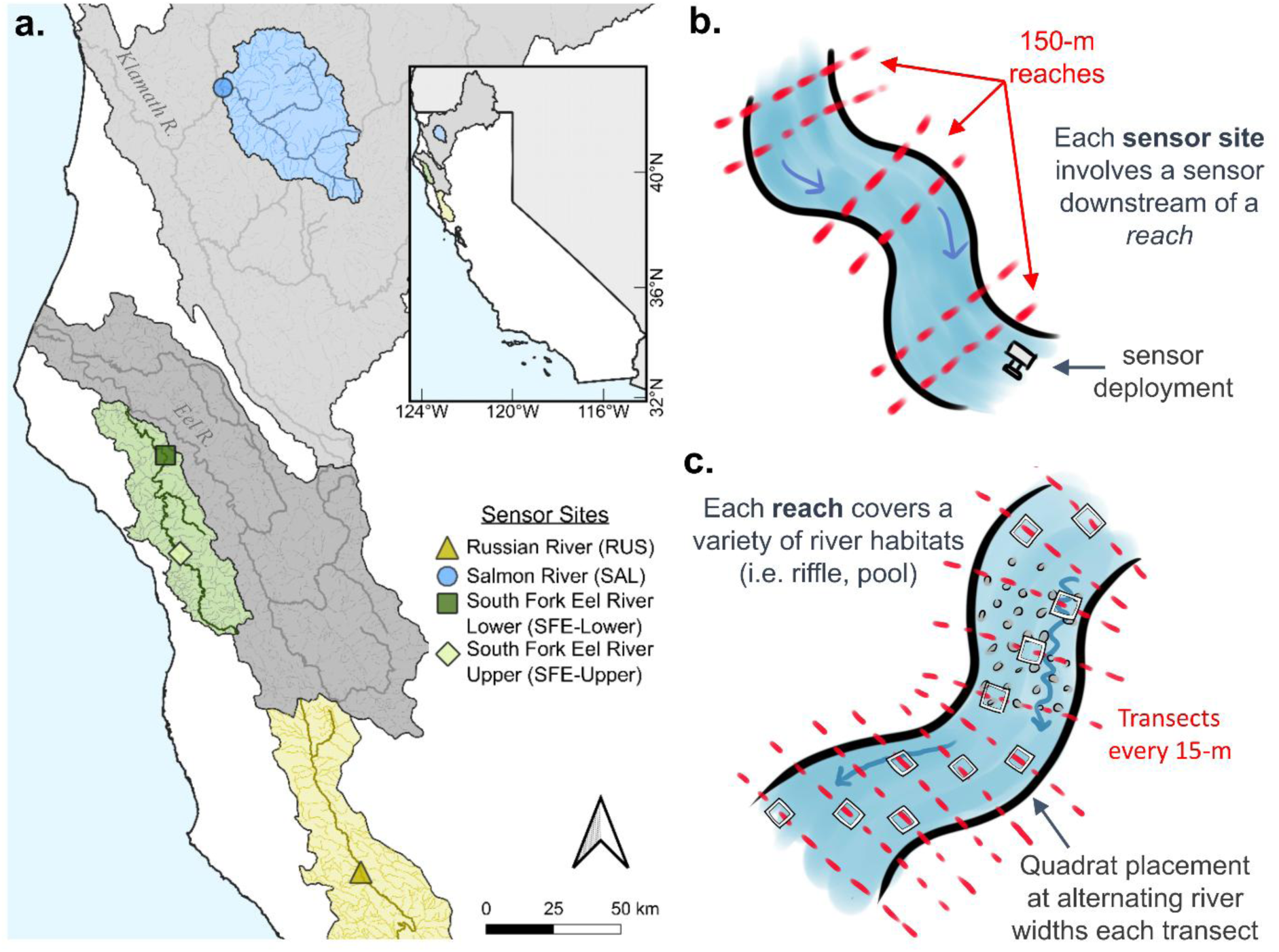
(a) Map of study locations in northern California. Watershed base maps are from the U.S. Watershed Boundary Dataset (U.S. Geological Survey 2021). (b) Overview of sensor deployment with three 150-m reaches upstream. (c) Overview of a single 150-m reach and benthic survey design which involved placing a quadrat at alternating river widths every 15-m transect (red dashed lines).

### Sampling Overview

To compare patterns of benthic cyanobacterial dynamics and GPP across rivers (Q1), we sampled all three rivers biweekly (i.e., every other week) from June to September in 2022 for a total of seven sampling events on each river (Appendix 1: Table S2). Due to wildfires, sampling on the Salmon River was interrupted from mid-August to mid-September, reducing the total number of sampling events to four. In each river, we sampled upstream of a dissolved oxygen sensor (Figure 2b) at three 150 m reaches (Figure 2c) that spanned a variety of river habitats (i.e., riffle, run, and pool) when possible. These 150 m reaches were all located upstream of sensors within the estimated dissolved oxygen residence distance in which 95% of the dissolved oxygen has traveled before equilibrating with the atmosphere and leaving the river (Appendix S1: Table S3; Appendix S1: Section S1 for further details). Within this distance, a river is assumed to be relatively homogenous for metabolism estimation (Hall and Hotchkiss 2017).

To compare patterns of benthic cyanobacterial dynamics and GPP (Q2) and test predictive models (Q3), we sampled weekly at five 150 m reaches on only the South Fork Eel River from June to September in 2023 (Appendix 1: Table S4). These reaches included the three reaches on the South Fork Eel River sampled in 2022 and an additional reach nearby also upstream of the original sensor placement location. The fifth reach was located over 45 kilometers upstream of the other four reaches where we deployed an additional dissolved oxygen sensor. Due to river access limitations, we were only able to sample one reach upstream of this sensor. These two sensors and their associated reaches in the South Fork Eel River in 2023 are distinguished as “SFE-Lower” (original location of 2022 sampling) and “SFE-Upper” (2023 location over 45 kilometers upstream of SFE-Lower; Figure 2a). In 2023, all four SFE-Lower reaches were sampled for a total of fifteen events and the single SFE-Upper reach was sampled for fourteen events. All collected data described are archived in the Environmental Data Initiative (Zabrecky et al. 2025).

### Benthic Surveys and Cyanobacteria Sample Collection

We estimated taxon-specific benthic cyanobacterial cover at each 150 m reach in general accordance with the Reachwide Benthos method from the California State Water Resources Control Board Surface Water Ambient Monitoring Program (SWAMP) Standard Operating Procedure (Ode et al. 2016). Using this method, we threw a 0.5 x 0.5-m quadrat along alternating points along the river cross section (25%, 50%, and 75% of river width) at each 15-m interval of the 150-m reach for a total of eleven quadrat placements. Within each quadrat, we estimated the percent cover of bare sediment or thin biofilm, green algae, *Microcoleus*, *Anabaena*/*Cylindrospermum* (which we grouped together as they were difficult to distinguish macroscopically in the field), and other nitrogen-fixing cyanobacteria (typically *Nostoc*). Where *Microcoleus* or *Anabaena*/*Cylindrospermum* was present but not captured in the quadrat, we recorded the transect interval where we observed it nearby.

We collected composite samples of *Microcoleus* and *Anabaena*/*Cylindrospermum* separately by combining multiple mat subsamples from 5-15 locations across each reach to account for any spatial variability in anatoxin concentrations (Wood et al. 2010; Appendix S1: Figure S1). Composite samples were collected with gloved hands for a final volume of ∼200 mL in amber plastic containers. If no *Microcoleus* or *Anabaena*/*Cylindrospermum* were found, we searched for ten minutes to confirm their absence and did not collect a composite sample for that taxon at that reach. After collection, the composite samples were shaken in their containers for thirty seconds to homogenize the sample and stored on ice during transport. After transport, samples were kept at 4 ℃ for less than 24 hours until they were re-homogenized and subsampled for analyses described in the following subsections.

### Microscopy

For microscopy, we preserved 5-35 mL of each composite sample in a clear falcon tube with 2% glutaraldehyde and stored in a dark box at 4 ℃ until they were analyzed. In 2023, we also saved a portion of sample for live microscopy analyses. Preserved samples were analyzed within two years of collection (Ode et al. 2016) and live samples were analyzed within ten days. We completed microscopy to confirm the presence of *Microcoleus, Anabaena,* and/or *Cylindrospermum* in their respective composite samples and quantify their relative abundance (Appendix S1: Figure S2; Appendix S1: Section S2 for further details). For both *Microcoleus* and *Anabaena*/*Cylindrospermum* composite samples, we were able to confirm their respective presence in all samples. *Microcoleus* accounted for an average of 75.0% of the algal composition in *Microcoleus* composite samples while *Anabaena* and *Cylindrospermum* accounted for an average of 35.9% of the algal composition in *Anabaena*/*Cylindrospermum* composite samples with the remainder consisting of mostly green algae (25.0%) and diatoms (11.0%; Appendix S1: Figure S3). Other potential toxin-producing taxa such as *Microcoleus*, *Geitlerinema*, and *Oscillatoria* were often found in *Anabaena/Cylindrospermum* composite samples.

### Anatoxin Extraction and Measurement

To measure anatoxins in *Microcoleus* and *Anabaena*/*Cylindropsermum* composite samples, we froze ∼40 mL of each composite sample in an amber falcon tube at −20 °C until lyophilization. Frozen composite samples were lyophilized at the University of Nevada, Reno and homogenized to a fine powder with pestle and mortar. When possible, 200 ± 50 mg of lyophilized material was suspended in 10 mL of 50% methanol. For samples with less material available, 100 ± 50 mg of material was suspended in 5 mL of 50% methanol and <50 mg was suspended in 2 mL of 50% methanol. We sonicated samples for three 20 s bursts at ∼15 W using a probe sonicator (Fisherbrand Model 505 Sonic Dismembrator, Fisher Scientific, Hampton, NH) and then centrifuged for 10 min at 2040 *g* (Sorvall Legend RT Refrigerated Benchtop Centrifuge, Sorvall, Newton, CT). We filtered 2 mL of the supernatant through a 0.4 μm nylon filter (Simsii, Issaquah, WA) into a liquid chromatography vial and stored at −20 ℃ until shipping. Samples were shipped overnight on ice to the State University of New York College of Environmental Science and Forestry (SUNY ESF) for analyses where samples were analyzed with liquid chromatography tandem mass spectrometry (LC-MS/MS) using a Waters ACQUITY TQD mass spectrometer (Waters, Milford, MA) coupled with a Waters ACQUITY UPLC solvent delivery system using a 2.0 x 150 mm Ace C-18 column. We analyzed for six anatoxin congeners including anatoxin-a and homoanatoxin-a and their *α* and β dihydro epimers following a modified version of U.S. Environmental Protection Agency (U.S. EPA) method 545 (U.S. EPA 2015) that includes additional congeners and confirmation ions for each analyte (Smith et al. 2020). Dihydroanatoxin-a and dihydrohomoanatoxin-a were both reported as the sum of their *α* and β epimer concentrations. Instrument detection limits were calculated based on standards and method detection limits were calculated for each sample individually based on the instrument detection limit, sample weight, and extractant volume and ranged from 0.008 to 0.23 μg anatoxin per g lyophilized sample.

We normalized anatoxin concentrations to both percent organic matter (OM) and chlorophyll-*a* of the lyophilized sample weight (Appendix S1: Figure S4). Chlorophyll-*a* was measured with the remainder of lyophilized composite samples using the hot ethanol extraction method (Sartory and Grobbelaar 1984) on a Trilogy Laboratory Fluorometer (Turner Designs, San Jose, California; Appendix S1: Section S3 for further details). For OM, we dried ∼3 g of wet samples within 24 hours of collection in a pre-weighed tin at 60 ℃ for at least 48 hours, weighed, combusted at 500 ℃ for two hours, and reweighed. Percent OM was calculated as the percentage of mass lost during combustion. For analyses, we reported anatoxin concentrations as the sum of all anatoxin congener concentrations per g OM.

We also analyzed samples for microcystins and cylindrospermopsins; however, as there were few detections of microcystins and no detection of cylindrospermopsins we do not focus on them here (Appendix S1: Table S5; Appendix S1: Section S4 for further details).

### Water Chemistry

We measured temperature (℃) and pH (Orion Star A121 portable pH meter, Thermo Fisher Scientific, Waltham, MA) and conductivity (μS/cm) and dissolved oxygen (mg/L) (Oakton 3000 series multiparameter smart handheld meter, Environmental Express, Charleston, SC) at each reach every visit. Dissolved oxygen, pH, and conductivity were all calibrated the day before field sampling. At each reach, we also collected a 60-mL surface water sample from a well-mixed portion of the river that was filtered using a 25 mm GF/F filter (Whatman, Clifton, New Jersey) into an acid-washed bottle. Surface water samples were stored on ice during transport and then frozen at −20 ℃ until analyses for orthophosphate (i.e., inorganic phosphate, PO_4_^3-^) and dissolved inorganic nitrogen (DIN; calculated as the sum of ammonium, NH_4_, and nitrate, NO_3_) at the University of Nevada, Reno using a SEAL AQ400 discrete analyzer (SEAL Analytical, Mequon, WI; Appendix S1: Section S5 for further details).

### Sensor Deployment and GPP Modeling

To estimate gross primary productivity (GPP) in each river, we deployed a miniDOT dissolved oxygen and water temperature sensor (Precision Measuring Engineering, Vista, CA) recording at 5-minute intervals downstream of survey reaches throughout the entirety of the sampling period (mid-June to late September) in both 2022 and 2023. Sensors were deployed at or within 10.5 kilometers of a U.S. Geological Survey (USGS) gage which provided continuous discharge. For sensors not deployed directly at the USGS gage, we verified the correspondence between river discharge at the sensor deployment location and USGS gage by measuring discharge at the sensor deployment location with a SonTek FlowTracker2 handheld Acoustic Doppler Velocimeter (SonTek, San Diego, California) biweekly using the velocity-area method (Herschy 1993; Appendix S1: Figure S5). We downloaded sensor data and cleaned sensors before redeployment each field visit. After each summer of sampling, we adjusted the data to account for sensor drift following a dissolved oxygen calibration as instructed by manufacturer protocols (PME 2021). Due to biofouling on the sensors that we placed in the Salmon and Russian Rivers, we instead used dissolved oxygen data from the Karuk Tribe with consent (Salmon River) and from the USGS (gage no: 11463000; Russian River).

We estimated daily river GPP at each sensor location with a single station approach (Hall and Hotchkiss 2017) using the “streamMetabolizer” package in R (R core Team 2024, Appling et al. 2018) which combines dissolved oxygen, temperature, river depth, barometric pressure, and light in a Bayesian state-space model to produce estimates for gross primary productivity (GPP; g O_2_ m^-2^ d^-1^), ecosystem respiration (ER; g O_2_ m^-2^ d^-1^), and an O_2_ gas exchange rate (*K_600_*; d^-1^).

We obtained discharge from nearby USGS gages using the “dataRetrieval” R package (De Cicco et al. 2024), barometric pressure data from the Global Land Data Assimilation System (GLDAS; Rodell et al. 2004), and light data from the North American Land Data Assimilation System (NLDAS; Xia et al. 2012). To obtain a depth-discharge relationship, we used a combination of at least eighty depth measurements we collected via kayak surveys within the dissolved oxygen footprint and USGS channel morphology and USGS discharge data (U.S. Geological Survey 2025) to create a linear model between log-transformed average depth and log-transformed discharge (Leopold and Maddock 1953) (Appendix S1: Figure S6). We calculated informed *K_600_* priors for each site using an equation incorporating velocity, depth, and slope (Raymond et al. 2012) (Appendix S1: Table S6). All Gelman-Rubin statistics (r-hats) of GPP, ER, and *K_600_* for final estimates were below 1.05, indicating model convergence (Brooks and Gelman 1998; Appendix S1: Figure S7). Pelagic contributions to GPP were not measured and assumed to be negligible as these rivers are very clear with relatively low nutrient concentrations (Appendix S1: Table S1).

For more details on sensor calibration, data processing, depth-discharge relationships, and model specifications, see supporting information (Appendix S1: Section S6).

### Data Analyses

For each sampling date and reach, we averaged category-specific percent cover estimates (i.e., bare/biofilm, *Microcoleus*, *Anabaena*/*Cylindrospermum*, green algae, and other nitrogen-fixing cyanobacteria) from all 11 quadrat placements. To compare temporal benthic cyanobacterial dynamics across rivers (Q1), we averaged taxon-specific percent cover and anatoxin concentrations across all reaches within a river on a sampling date. In cases when *Microcoleus* or *Anabaena*/*Cylindrospermum* were present but anatoxins were not detected or, in some cases, not enough material was available for anatoxin extraction (n= 2 in 2022 and n = 17 in 2023), we assumed a value of 0 for all data analyses (Q1-3).

To understand how GPP may influence predictions of taxon-specific cover and anatoxin concentrations in addition to abiotic covariates (Q3), we compared a series of models predicting taxon-specific benthic cyanobacterial cover and anatoxin concentrations using weekly 2023 data from the five reaches in the South Fork Eel River. We compared predictions of seven model types built with different covariate categories (Table 1) including physical variables (water temperature and discharge), water chemistry variables (dissolved inorganic nitrogen, orthophosphate, and conductivity), and biological variables (GPP). To match daily GPP estimates with other covariates that were measured weekly, we used the median GPP across the four days prior to the sampling day (with the exception of the initial time step that used the estimate from that day). For models predicting taxon-specific anatoxins, we also examined how predictive accuracy changed with the inclusion of that taxon’s percent cover as a predictor. As we were primarily interested in predicting the timing rather than differences in magnitude across reaches, we normalized our response variables (i.e., cover, anatoxins) to the maximum observed at each reach bounding the response variables from 0 to 100. Covariates were standardized within each reach using the “scale” function in R (Appendix S1: Figure S8).

**Table 1.**
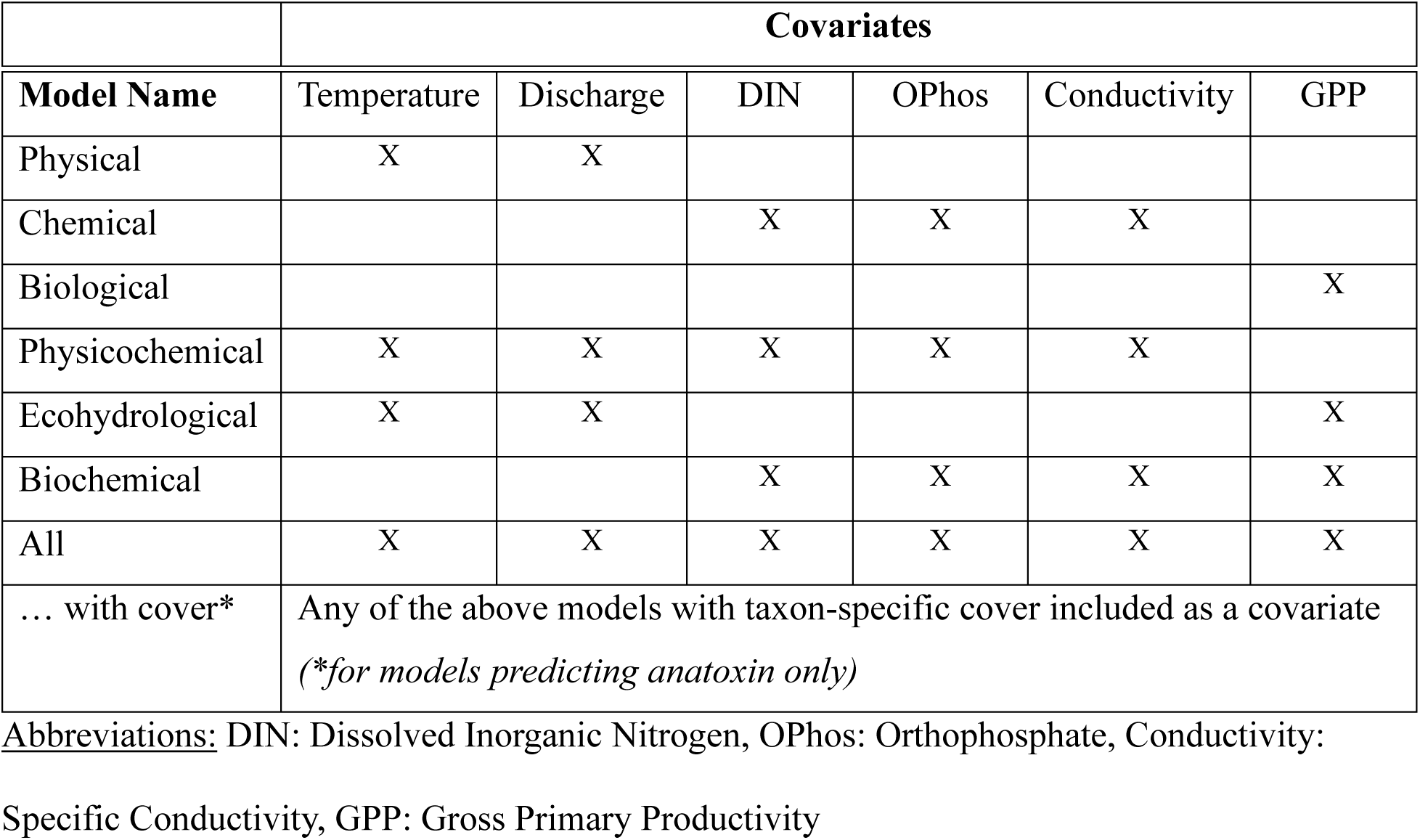
Covariates included in each predictive model as indicated by “X”.

| Model Name | Temperature | Discharge | DIN | OPhos | Conductivity | GPP |
| --- | --- | --- | --- | --- | --- | --- |
| Physical | X | X |  |  |  |  |
| Chemical |  |  | X | X | X |  |
| Biological |  |  |  |  |  | X |
| Physicochemical | X | X | X | X | X |  |
| Ecohydrological | X | X |  |  |  | X |
| Biochemical |  |  | X | X | X | X |
| All | X | X | X | X | X | X |
| ... with cover* | Any of the above models with taxon-specific cover included as a covariate<br>(*for models predicting anatoxin only) |  |  |  |  |  |

Abbreviations: DIN: Dissolved Inorganic Nitrogen, OPhos: Orthophosphate, Conductivity:
Specific Conductivity, GPP: Gross Primary Productivity

We predicted taxon-specific cover and anatoxin concentrations at a near-weekly time step (5-8 days) using measured covariates (C) and the predicted cover (y) from the prior-time step.

We used the following autoregressive process model to predict cover:

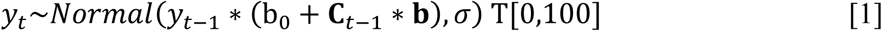

where y is normalized cover at a near-weekly timestep (t), **C** is a matrix of covariates, **b** is a vector of coefficients for the covariates (depending on the model, Table 1), b_0_ is an intercept term, σ is the variance, and T[0,100] indicates that y arises from a normal distribution truncated at 0 and 100. In this cover model, we substituted a small value (0.05) for zero percent cover to allow the model to increase. When making predictions, the initial y_t-1_ value was set to this small value as we observed zero cover on the first sampling date at all reaches. All subsequent y_t-1_ values used to predict y_t_ were values predicted by our model.

For prediction of taxon-specific anatoxin concentrations, we did not include an autoregressive term as anatoxins can be highly variable across time (Heath et al. 2011) and degrade rapidly (Kaminski et al. 2013). We used the following model to predict anatoxins:

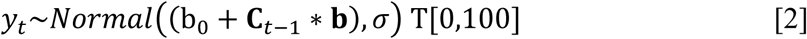

where y is normalized anatoxin concentrations at a near-weekly timestep (t), **C** is a matrix of covariates, **b** is a vector of coefficients for the covariates (depending on the model, Table 1), b_0_ is an intercept term, σ is the variance, and T[0,100] indicates that y arises from a normal distribution truncated at 0 and 100.

In both predictive models [eq. 1 & 2], both the intercept term (b_0_), covariate coefficients (**b**), and sigma (σ) were given priors normally distributed around zero as follows:

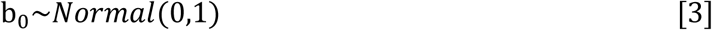

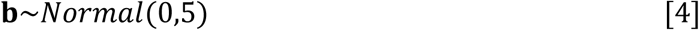

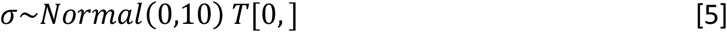

where T[0,] indicates a normal distribution with a lower bound of 0 and no upper bound.

We fit models under a leave-one-out cross validation framework in which we fit a sub-model with data from four out of the five reaches and tested predictive accuracy by predicting the withheld fifth reach for a total of five sub-models for a single model. In other words, a single model (e.g., “physical” model predicting *Microcoleus* cover) was fit five times to five different subsets of data for five sub-models. All models were fit using Bayesian inference via Hamiltonian Monte Carlo in Stan (Carpenter et al. 2017) and the RStan interface (Stan Development Team 2024) in R version 4.4.2 (R Core Team 2024). We ran models for 10,000 iterations, half of which were warmup. All but four models converged with no errors and parameter r-hats all under 1.05 (Appendix S1: Figure S9). The four models that were unable to converge were models predicting *Anabaena*/*Cylindrospermum* cover (“chemical”, “physicochemical”, “biochemical”, and “all” models with reach SFE-Lower-1S withheld) and were excluded from further analysis.

To compare predictive accuracy across models, we calculated normalized root mean squared error (NRMSE) for each predicted time series by dividing the root mean squared error (RMSE) of the predictions by the range of the data. NRMSE is bounded by 0 to 1 and lower values indicate greater predictive accuracy. To calculate a single NRMSE mean and 95% credible interval for each model, we compiled the posterior distributions from each sub-model. We also calculated a single mean and 95% credible interval for each covariate in a model by grouping the posterior distribution from each sub-model. We calculated the coefficient of determination, *r^2^*, for each sub-model using the “lm” function with the predicted values as a function of the observed values and averaged across those values to calculate a single *r^2^* for each model. To evaluate if our models performed better than simply predicting the average, we calculated NRMSEs for “null” models by predicting the average taxon-specific cover or anatoxin concentrations observed from across all five reaches and all sampling dates. We also compared performance among groups (e.g., *Microcoleus* vs. *Anabaena/Cylindrospermum* cover predictions) by comparing vectors of the NRMSE posteriors distributions and calculating the percentage of comparisons where one group had a lower NRMSE than the other.

Lastly, we calculated the relative contributions of parameter, process, and initial condition uncertainty in our predictions. Because of the idiosyncrasies of the truncated normal distribution, the typical approach of starting with parameter uncertainty and sequentially adding additional sources of uncertainty was not possible (Dietze 2017). Instead, we first quantified process error by generating predictions using mean parameter values within the truncated normal distribution. Parameter uncertainty was added with posterior Markov Chain Monte Carlo (MCMC) draws as in our original models above. Lastly, initial condition uncertainty was added to our autoregressive model predicting cover, by making predictions where the first autoregressive value was pulled from a distribution rather than the set value of 0.05. We used the following distribution which assumes that the initial cover in each reach will be low, but allowed for variability:

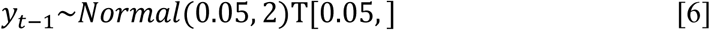

We quantified predictive uncertainty as the standard deviation of the posterior predictions for each sub-model. While we acknowledge our covariates or drivers (i.e., nutrients, temperature, etc.) also contain some degree of uncertainty, we omitted this step as driver uncertainty was assumed to be relatively small compared to other sources of uncertainty at our weekly prediction time step (Lofton et al. 2022). Incorporating covariate uncertainty is a potential next step in forecasting benthic cyanobacterial dynamics but is beyond the scope of this paper.

## Results

### Q1: Variability of benthic cyanobacterial dynamics and GPP across rivers

The presence of macroscopically visible *Microcoleus* and *Anabaena*/*Cylindrospermum* varied across rivers in 2022. Both taxa formed extensive mats in the South Fork Eel River with *Microcoleus* predominantly accruing in riffles and *Anabaena/Cylindrospermum* forming emerald-green towers on *Cladophora* tufts in pools or accumulating on fine sediment or gravel (Appendix S1: Figure S2, Figure S10). However, in the Russian River, we macroscopically observed only *Anabaena*/*Cylindrospermum* (although *Microcoleus* filaments were present in some samples via microscopy) and in the Salmon River, we macroscopically observed only *Microcoleus* until the end of the field season in late September when small amounts of *Anabaena*/*Cylindrospermum* were observed.

The timing and magnitude of cyanobacterial mat cover was specific to each taxon and river (Figure 3a-c; Appendix S1: Table S7, Figure S11). In the South Fork Eel River, *Microcoleus* began as small mats in June that continued to accrue until late September (mean cover across all reaches and sampling dates: 2.9%; mean cover across all reaches on peak date: 12.0%) and eventually comprised >25% cover within riffle habitats. We did not observe *Anabaena*/*Cylindrospermum* in the South Fork Eel River until mid-July when it quickly reached its maximum cover (mean cover across all reaches and sampling dates: 3.4%; mean cover across all reaches on peak date: 25.9%) and then nearly disappeared (<1% mean cover) by late August.

**Figure 3.**
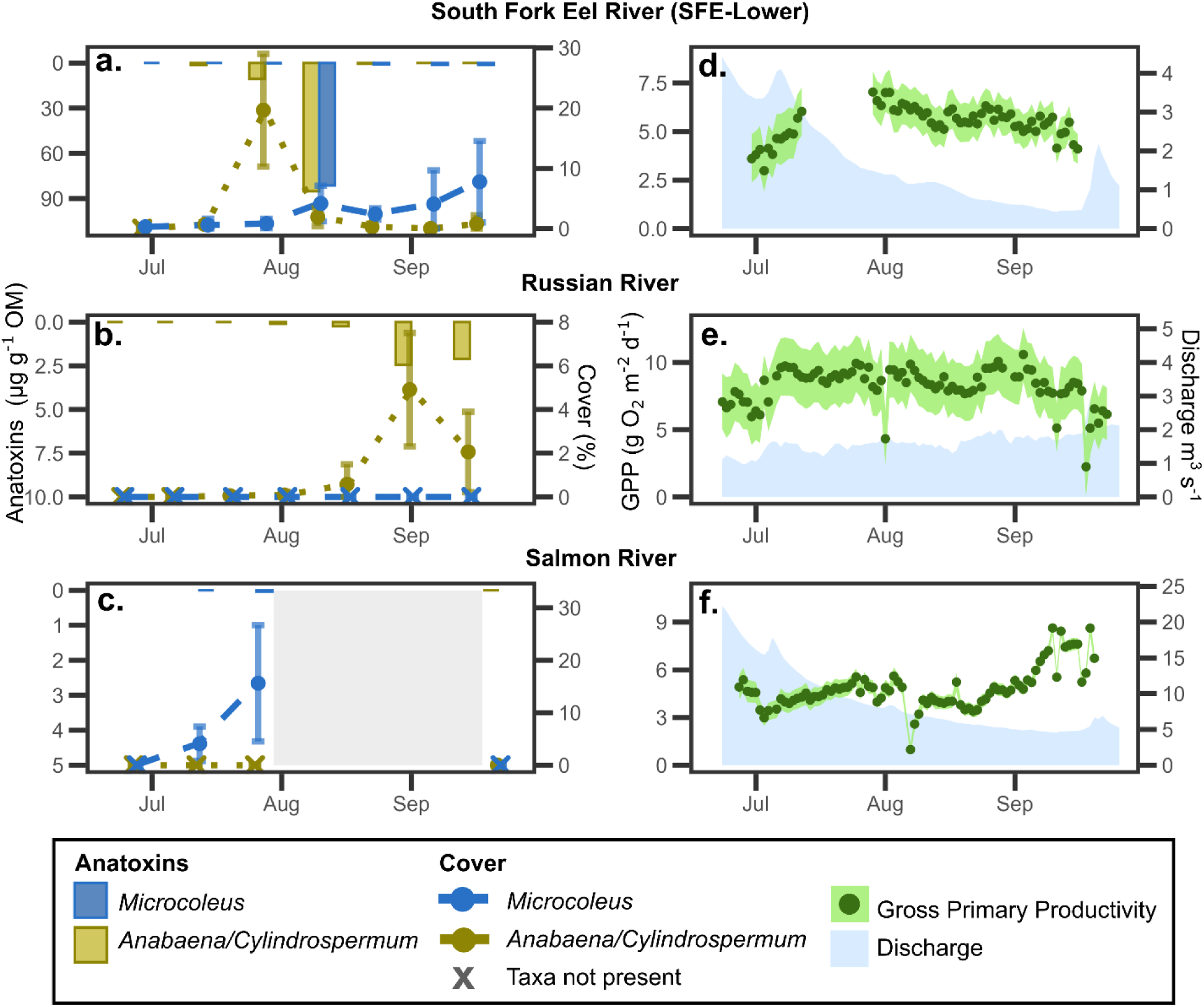
(a-c) Taxon-specific benthic cyanobacterial cover and anatoxin concentrations and (d-f) gross primary productivity (GPP) estimates and discharge for the South Fork Eel River, Russian River, and Salmon River, respectively in 2022. Anatoxins are shown as the mean across all three reaches (bars associated with the left y-axis where absence of a bar indicates no sample was taken or not enough sample was available for analysis on any reach of the river). Cover is shown as the mean across all three reaches (points and lines associated with the right y-axis) with +/- one standard deviation (error bars). GPP is shown as the mean (point) with 95% credible interval (ribbon) associated with the left y-axis and discharge is shown as the blue area associated with the right y-axis.

In the Russian River, we observed a similar peak and decline in *Anabaena*/*Cylindrospermum* cover (mean cover across all reaches and sampling dates: 1.2%; mean cover across all reaches on peak date: 7.5%) but the magnitude of cover was smaller and occurred over a month later than in the South Fork Eel River. In the Salmon River, we observed *Microcoleus* first in early July and saw its rapid accrual until sampling interruption in August from wildfires (mean cover across all reaches and sampling dates, 5.4%; mean cover across all reaches on peak date, 27.7%). Upon return to the Salmon River in late September, we found no visible *Microcoleus*.

Anatoxin presence and concentrations also varied in magnitude and timing across the three rivers (Figure 3a-c; Appendix S1: Table S7, Table S8, Figure S11). Across all sampling dates, composite samples from the South Fork Eel River had the highest measured anatoxin concentrations (mean concentration across all reaches at peak: >80 μg anatoxins g^-1^ OM), while the samples from Russian River had considerably lower concentrations (mean concentration across all reaches at peak: <2.5 μg anatoxins g^-1^ OM), and at the Salmon River, all but a single sample had no detectable anatoxins. Within the South Fork Eel River, anatoxin concentrations were similar across focal taxa (for *Microcoleus* and *Anabaena*/*Cylindrospermum*: mean across all samples was 12.1 and 14.0 μg anatoxins g^-1^ OM, respectively; maximum for a single composite sample was 237.0 and 155.2 μg anatoxins g^-1^ OM, respectively). While both taxa in the South Fork Eel River reached peak anatoxin concentrations in early August, initial *Microcoleus* composite samples from late June and early July did not contain any detectable anatoxin; whereas, for *Anabaena*/*Cylindrospermum* composite samples, only samples collected in September did not contain any detectable anatoxin. In the Russian River, anatoxins were detected in *Anabaena*/*Cylindrospermum* composite samples beginning in August until the end of the field season, peaking in early September, though with overall much lower concentrations (mean across all samples: 0.07 μg anatoxins g^-1^ OM; maximum for a single composite sample: 5.3 μg anatoxins g^-1^ OM) than those from the South Fork Eel River. In contrast, on the Salmon River, anatoxins were only detected in a single *Microcoleus* composite sample taken in late July that had relatively low concentrations (0.11 μg anatoxins g^-1^ OM). The single *Anabaena*/*Cylindrospermum* composite sample from the Salmon River did not contain any detectable anatoxin. The dominant anatoxin congener detected also varied across rivers with anatoxin-a being most dominant in the South Fork Eel River, dihydroanatoxin-a in the Russian River, and homoanatatoxin-a as the only detected congener in the Salmon River (Appendix S1: Figure S12).

The timing and magnitude of GPP also varied across rivers (Figure 3d-f; Appendix S1: Table S7). GPP in the South Fork Eel River increased in July to a peak in early August (though possibly earlier but unknown due to sensor malfunctioning), followed by a steady decrease for the rest of the field season (mean [range]: 5.4 [3.0-7.0] g O_2_ m^-2^ d^-1^). GPP in the Salmon River generally increased throughout the field season, except for a decrease in early August associated with smoke and elevated turbidity from a nearby wildfire (mean [range]: 4.8 g [1.0-8.6] O_2_ m^-2^ d^-1^). In contrast, GPP in the Russian River was fairly consistent at a relatively high magnitude throughout the summer until a small decrease beginning in September (mean [range]: 8.3 [2.2-10.5] g O_2_ m^-2^ d^-1^).

### Q2: Variability of benthic cyanobacterial dynamics among reaches within the South Fork Eel River

In 2023, during weekly sampling at five reaches in the South Fork Eel River, the timing of peak *Microcoleus* and *Anabaena*/*Cylindrospermum* cover was consistent across most reaches, but the magnitude of the cyanobacterial mat cover at peak varied among reaches by up to 15% for *Microcoleus* and 21% for *Anabaena*/*Cylindrospermum* among reaches (Figure 4a; Appendix S1: Table S9). *Microcoleus* appeared in late June to early July in four of five reaches, peaking in late September (reported as range across all reaches, mean cover across all sampling dates: 1.1-8.6%; maximum cover: 8.4-24.1%). The one reach (SFE-Lower-4) that contradicted this pattern contained less riffle habitat than other reaches and had peak *Microcoleus* cover during mid-August. *Anabaena*/*Cylindrospermum* appeared in all reaches in mid-July with cover peaking in mid-August in all but one reach (reported as range across all reaches, mean cover across all sampling dates: 0.03-4.3%; maximum cover: 0.5-22.3%). Subsequently, *Anabaena*/*Cylindrospermum* declined to a sparse and intermittent presence until sampling ended in late September. The one reach (SFE-Lower-2) that contradicted this pattern lacked a pool with abundant *Cladophora* and had *Anabaena*/*Cylindrospermum* cover peak in mid-September.

**Figure 4.**
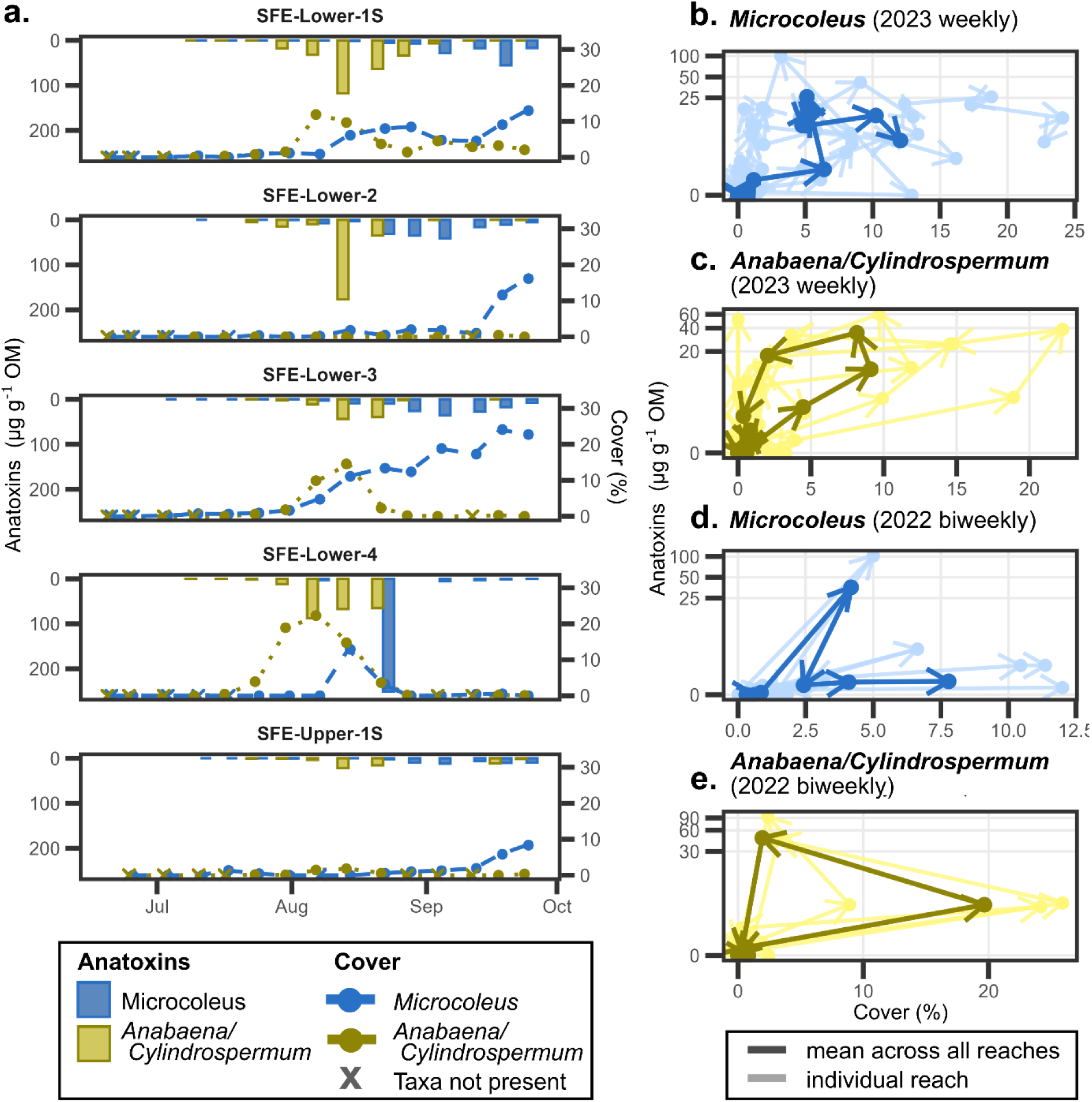
(a) Taxon-specific benthic cyanobacterial cover and anatoxin concentrations for the five study reaches within the South Fork Eel River in 2023. Anatoxins are shown in bars (associated with the left y-axis) where absence of a bar indicates no sample was taken or not enough sample was available for analyses. Cover is shown in points and lines (associated with the right y-axis). (b-e) Taxon-specific benthic cyanobacteria mat anatoxin concentrations versus cover for (b) *Microcoleus* and (c) *Anabaena*/*Cylindrospermum* sampled weekly in 2023 and (d) *Microcoleus* and (e) *Anabaena/Cylindrospermum* sampled biweekly in 2022 both in the South Fork Eel River. Mean behavior across all reaches is shown with the darker bolded line and individual reach behavior are shown with lighter lines. Arrows indicate direction of time across the summer.

In 2023, temporal patterns of taxon-specific anatoxin concentrations were also generally consistent across reaches within the South Fork Eel River, but concentrations varied up to 20-fold (Figure 4a; Appendix S1: Table S9, Table S10, Table S11). As observed in 2022 at the South Fork Eel River, *Microcoleus* composite samples did not contain detectable anatoxin until late July. Anatoxin concentrations in *Microcoleus* composite samples peaked across reaches in late-August to mid-September (reported as range across all reaches, mean across all sampling dates: 3.6-17.6 μg anatoxins g^-1^ OM; maximum: 12.0-250.4 μg anatoxins g^-1^ OM) which occurred a couple weeks later than peak concentrations in 2022. For *Anabaena*/*Cylindrospermum* composite samples, the highest anatoxin concentrations were observed slightly earlier in mid-August (reported as range across all reaches, mean across all sampling dates: 3.9-18.1 μg anatoxins g^-1^ OM; maximum: 21.7-118.1 μg anatoxins g^-1^ OM) closer to when we observed maximum anatoxin concentrations in the prior year. Also, similar to the prior year, non-detects in *Anabaena*/*Cylindrospermum* composite samples were uncommon until September. The dominant detected anatoxin congener across all reaches for both taxa was anatoxin-a as also observed in the South Fork Eel River the prior year (Appendix S1: Figure S12).

Relationships between benthic cyanobacterial cover and anatoxin concentrations exhibited taxon-specific patterns that were synchronous across four of five reaches (Figure 4b; Appendix S1: Figure S13). *Anabaena*/*Cylindrospermum* displayed a hysteresis-like pattern whereby increases in cover preceded increases in *Anabaena*/*Cylindrospermum* anatoxin concentrations and both concurrently declined (Figure 4c). Conversely, increases in *Microcoleus* cover did not lead to increases in *Microcoleus* anatoxin concentrations (Figure 4b). These taxon-specific patterns were also evident in our prior year of sampling on the South Fork Eel River in 2022 (Figure 4d-e).

### Q3: Predictive Modeling within the South Fork Eel River

Models predicting *Microcoleus* cover had a 77 % probability of having a lower NRMSE than models predicting *Anabaena/Cylindrospermum* cover (Figure 5a-b; mean NRMSE across all models: 0.35 fpr *Microcoleus*, 0.52 for *Anabaena*/*Cylindrospermum*). Only two of seven models predicting *Microcoleus* cover (“physical” and “ecohydrological” models) had a mean NRMSE lower than the null model NRMSE (mean NRMSE: 0.32) although their 95% credible intervals always went above the null model NRMSE (mean NRMSE 95% credible interval across all models: 0.16 to 0.62). Conversely, no models predicting *Anabaena*/*Cylindrospermum* cover had a mean NRMSE lower than the null model NRMSE (NRMSE: 0.31), and only five of seven models had 95% credible intervals low enough to incorporate the null model NRMSE (mean NRMSE 95% credible interval across all models: 0.29 to 0.72). Furthermore, models predicting *Microcoleus* cover explained a higher proportion of the variance than models predicting *Anabaena*/*Cylindrospermum* cover (for *Microcoleus* vs. *Anabaena*/*Cylindrospermum* cover models; mean [range] *r^2^*: 0.48 [0.26-0.62] vs. 0.20 [0.11-0.24], respectively). The incorporation of GPP as a covariate was more influential in predicting *Anabaena*/*Cylindrospermum* cover than *Microcoleus* cover. The best performing models for predicting *Microcoleus* cover all included discharge as a covariate (i.e., “physical”, “physicochemical”, “ecohydrological”, and “all” models; Figure 6a; Appendix S1: Figure S14). For these models, discharge had a negative mean posterior estimate indicating that *Microcoleus* cover was higher with lower discharge (Figure 6b). In contrast, GPP and all other covariates except for discharge and orthophosphate had posterior estimates with 95% credible intervals that overlapped zero in some or all models, indicating uncertainty in whether the variable had a positive or negative effect on *Microcoleus* cover (Appendix S1: Figure S15). This was also the case for GPP and all other covariates except specific conductivity in models predicting *Anabaena/Cylindrospermum* cover (Appendix S1: Figure S16). The best performing model predicting *Anabaena/Cylindrospermum* cover was the “biological” model, which included GPP as the only covariate (Figure 7a; Appendix S1: Figure S17). In this model, GPP had a small positive effect size (Figure 7b); however, the model failed to capture peak cover. Moreover, GPP posterior estimate 95% credible intervals in models predicting *Anabaena/Cylindrospermum* cover always overlapped zero (Appendix S1: Figure S16).

**Figure 5.**
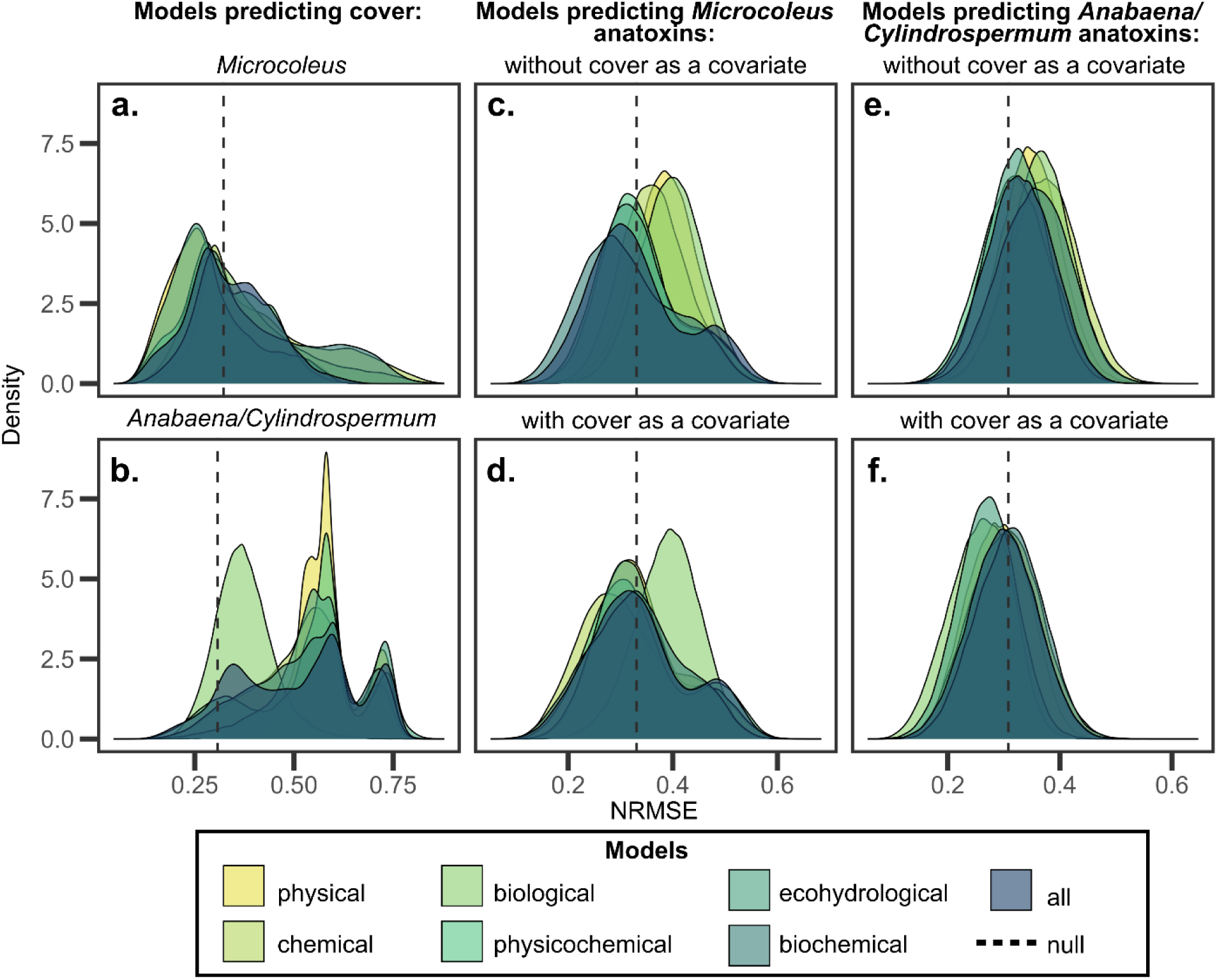
Posterior distributions of normalized root mean square errors (NRMSEs) for every model (a) predicting *Microcoleus* cover, (b) predicting *Anabaena/Cylindrospermum* cover, (c) predicting *Microcoleus* anatoxins without cover as a covariate, (d) predicting *Microcoleus* anatoxins with cover as a covariate, (e) predicting *Anabaena/Cylindrospermum* anatoxins without cover as a covariate, and (f) predicting *Anabaena/Cylindrospermum* anatoxins with cover as a covariate. Black dashed lines show the null model NRMSE. Covariates included in each model are indicated in Table 1.

**Figure 6.**
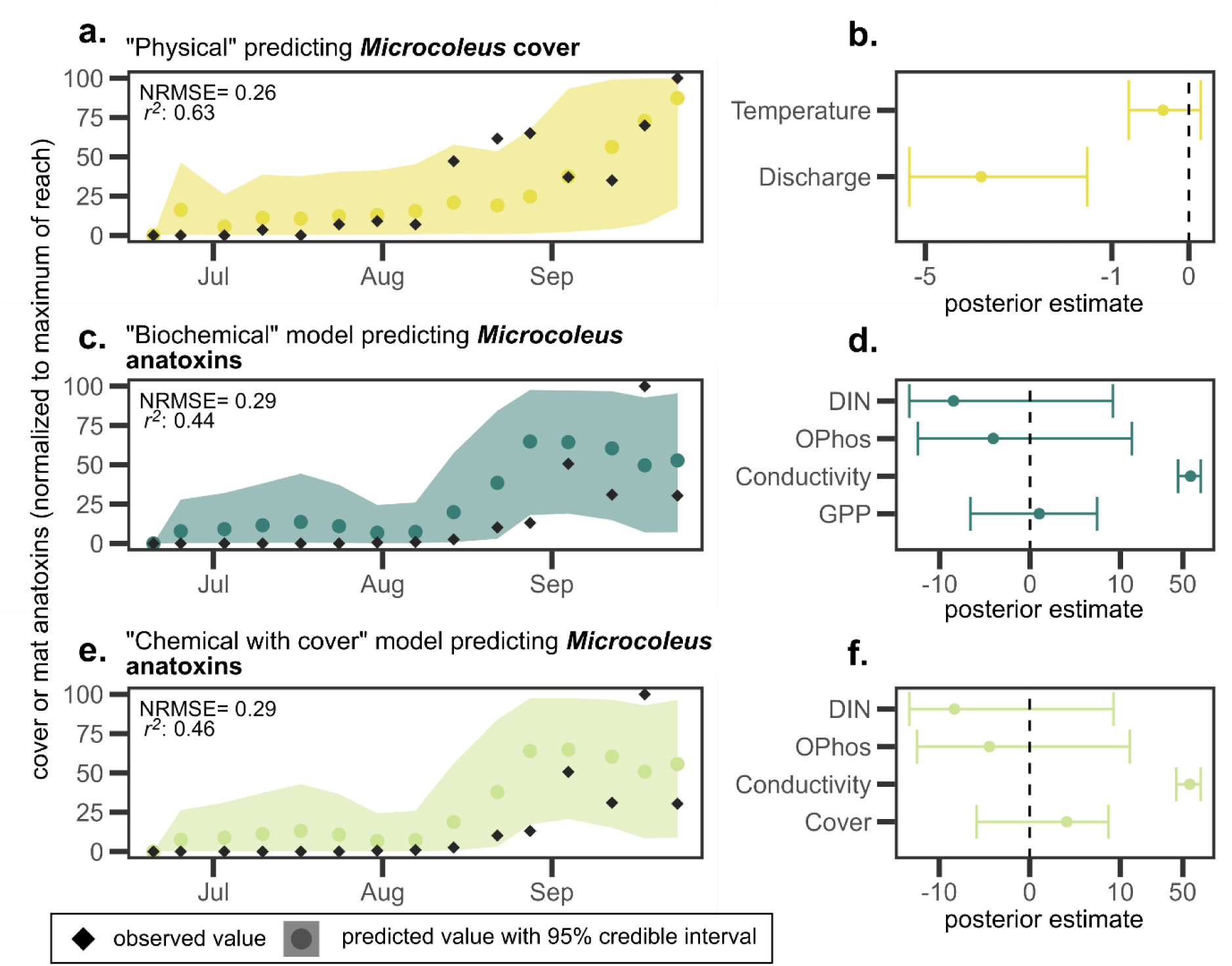
Model predictions and associated covariate posterior estimates for the model with the lowest mean overall normalized root mean square error (NRMSE) predicting *Microcoleus* (a, b) cover, (c-d) anatoxin concentrations (out of models excluding cover as a covariate), and (d-e) anatoxin concentrations (out of models including cover as a covariate). Predictions are shown using the sub-model predicting reach SFE-Lower-1S and covariate posterior estimates are from that of the full model (all sub-models averaged). Predictions and posterior estimates are both shown as mean (point) with 95% credible intervals (ribbon or range respectively).

**Figure 7.**
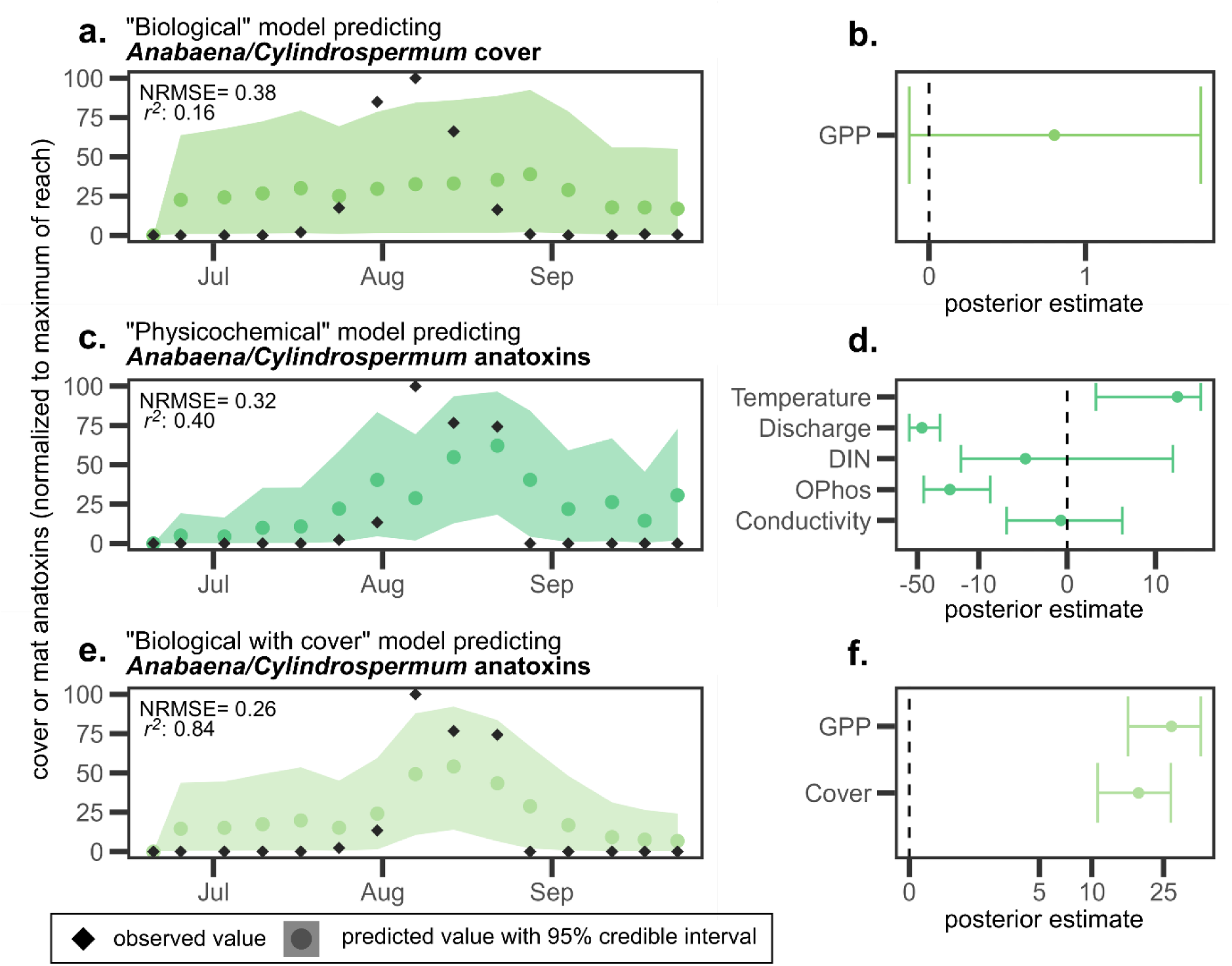
Model predictions and associated covariate posterior estimates for the model with the lowest mean overall normalized root mean square error (NRMSE) predicting *Anabaena/Cylindrospermum* (a, b) cover, (c-d) anatoxin concentrations (out of models excluding cover as a covariate), and (d-e) anatoxin concentrations (out of models including cover as a covariate). Predictions are shown using the sub-model predicting reach SFE-Lower-4 and covariate posterior estimates are from that of the full model (all sub-models averaged). Predictions and posterior estimates are both shown as mean (point) with 95% credible intervals (ribbon or range respectively).

In contrast to our models predicting cover, models predicting *Anabaena/Cylindrospermum* anatoxins had a 62% probability of having a lower NRMSE than models predicting *Microcoleus* anatoxins (Figure 5c-f). Six of fourteen models predicting *Anabaena*/*Cylindrospermum* anatoxins had a mean NRMSE lower than the null model NRMSE (NRMSE: 0.31) while only three of fourteen models predicting *Microcoleus* anatoxins had a mean NRMSE lower than the null model NRMSE (NRMSE: 0.32) though all models had 95% credible intervals going above the null NRMSE (mean NRMSE 95% credible interval across all models predicting *Anabaena/Cylindrospermum* vs. *Microcoleus* anatoxins: 0.20 tco 0.43 vs. 0.21 to 0.52, respectively). Models predicting *Anabaena/Cylindrospermum* anatoxins also explained a similar proportion of the variance (mean [range] *r^2^*: 0.32 [0.08-0.60]) compared to that of models predicting *Microcoleus* anatoxins (mean [range] *r^2^*: 0.36 [0.15-0.46]).

The incorporation of taxon-specific cover as a covariate reduced the NRMSE for models predicting *Anabaena*/*Cylindrospermum* anatoxins but less so for models predicting *Microcoleus* anatoxins (Figure 5e-f). Models predicting *Anabaena/Cylindrospermum* anatoxins with cover as a covariate had a 76% probability of having a lower NRMSE than those not including cover as a covariate while models predicting *Microcoleus* anatoxins with cover as a covariate only had a 56% chance of having a lower NRMSE than those not including cover as a covariate. The incorporation of cover as a covariate also increased the explained variability in *Anabaena/Cylindrospermum* anatoxin models but not *Microcoleus* models (mean *r^2^* for models with cover as a covariate vs. without cover as a covariate: 0.42 vs. 0.22 for *Anabaena/Cylindrospermum* and 0.37 vs. 0.35 for *Microcoleus*). Posterior estimates for cover were all positive for models predicting *Anabaena/Cylindrospermum* anatoxins (Figure 7f; Appendix S1: Figure S18) but were only slightly positive with 95% credible intervals overlapping zero for models predicting *Microcoleus* anatoxins which indicates an inconclusive effect (Figure 6f; Appendix S1: Figure S19).

Other than the inclusion of cover as a covariate, models predicting anatoxins both had relatively similar NRMSEs and a much smaller range across models than that for models predicting cover. The five models predicting *Microcoleus* anatoxins with the lowest mean NRMSEs all included chemical covariates (“chemical with cover”, “biochemical”, “biochemical with cover”, “all”, and “physicochemical with cover” models; Figure 6c, 6e; Appendix S1: Figure S20). In these five models, specific conductivity was the only covariate that consistently had a positive posterior estimate (Figure 6d, 6f; Appendix S1: Figure S20). In contrast, the five models predicting *Anabaena/Cylindrospermum* models with the lowest mean NRMSEs (“biological with cover”, “ecohydrological with cover”, “physical with cover”, “physicochemical with cover”, and “all with cover” model) showed no cohesive theme other than the inclusion of cover as a covariate (Figure 7c, 7e; Appendix S1: Figure S21).

Parameter and initial condition uncertainty contributed negligibly to predictive uncertainty in our predictions which suggests that the process uncertainty is the dominant source of predictive uncertainty (Figure 8). Parameter uncertainty increased predictive uncertainty by 1.0% across all models while initial condition uncertainty only increased predictive uncertainty by 0.1% across all models predicting cover. Furthermore, within each taxon and dynamic (*i.e.,* cover or anatoxin concentrations) being predicted, there were no substantial differences in the amount of predictive uncertainty added by parameter and initial condition uncertainty among different models. Overall, predictive uncertainty was highest for models predicting *Anabaena/Cylindrospermum* cover and lowest for models predicting *Anabaena/Cylindrospermum* anatoxins which echoes the patterns found in our predictive accuracy analyses above.

**Figure 8.**
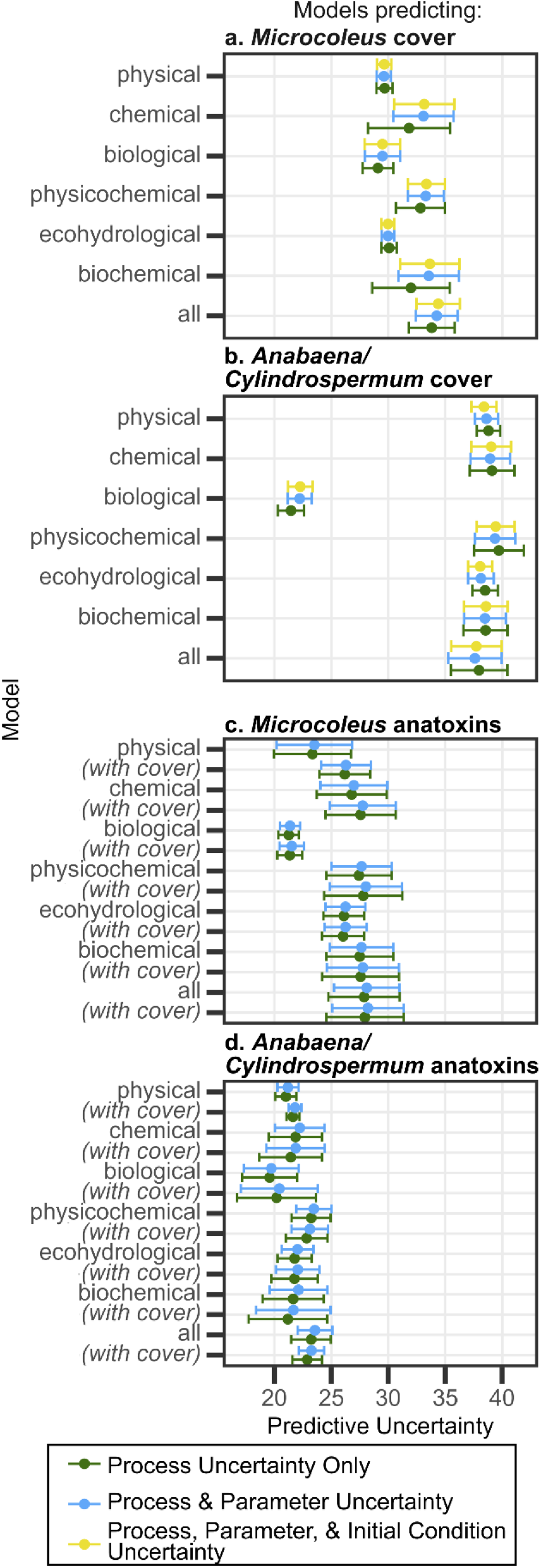
Mean predictive uncertainty (calculated as the standard deviation of the posterior predictions) across all sub-models (point) +/- one standard deviation (error bars) for models predicting (a) *Microcoleus* cover, (b) *Anabaena/Cylindrospermum* cover, (c) *Microcoleus* anatoxins, and (d) *Anabaena/Cylindrospermum*. Models predicting anatoxins labeled *with cover* refer to the model listed above with cover added as a covariate.

## Discussion

In this study evaluating temporal patterns of benthic cyanobacterial dynamics and GPP, we found three key results. First, across rivers within the same region, patterns in taxon-specific benthic cyanobacterial dynamics and GPP were highly variable suggesting that patterns from one river in our study cannot be extrapolated to other rivers, even within a relatively close proximity. Second, by sampling across multiple reaches within a single river, we found that while magnitudes of taxon-specific cover and anatoxin concentrations varied across reaches, the temporal patterns were mostly consistent which suggests that the timing of peak anatoxin exposure risk is more consistent within a river than among rivers. Third, we found our predictive models of *Microcoleus* cover outperformed those for *Anabaena*/*Cylindrospermum* cover, but our predictive models of *Anabaena*/*Cylindrospermum* anatoxins outperformed those for *Microcoleus* anatoxins. This was especially true with the incorporation of taxon-specific cover as a predictor, which suggests that visual assessments of cover within a reach may be an effective predictor of anatoxin concentrations for *Anabaena*/*Cylindrospermum* but not for *Microcoleus*. Taken together, our results demonstrate the importance of understanding the dynamics of each potential anatoxin-producing benthic cyanobacterial taxon within a river rather than treating toxigenic benthic cyanobacteria as a single population when evaluating exposure risk and attempting to make predictions or forecasts of benthic cyanobacterial dynamics.

### Q1: Variability of benthic cyanobacteria dynamics and GPP across rivers

While we found differences in the presence, timing, and magnitude of taxon-specific benthic cyanobacterial cover across rivers and taxa, the magnitude of anatoxin concentrations, the dominant anatoxin congener, and, to a lesser extent, the timing of maximum concentrations, were more consistent between taxa within the same river rather than for the same taxa among rivers. In our study, both *Microcoleus* and *Anabaena*/*Cylindrospermum* composite samples from the South Fork Eel River reached maximum anatoxin concentrations of similar magnitudes, predominantly composed of anatoxin-a, on the same date. In contrast, in the Russian River, maximum anatoxin concentrations in *Anabaena*/*Cylindrospermum* composite samples were more than 10 times lower than South Fork Eel composite sample concentrations and were predominantly composed of dihydroanatoxin-a. Similar to our results, in New Zealand, *Microcoleus* anatoxin concentrations and dominant anatoxin congener were variable among rivers (Wood et al. 2017a, 2018, McAllister et al. 2018a). However, when looking at two taxa within the same watershed (though mostly from one river), Bouma-Gregson et al. (2018) reported similar anatoxin-a concentrations among *Microcoleus*-dominated and *Anabaena*-dominated mats as we also found in our study. Together, these findings suggest that patterns in anatoxin concentrations observed in one river are not generalizable to a region but may be more similar among anatoxin-associated taxa within the same river. Additionally, as some anatoxin congeners may differ in their toxicity (Puddick et al. 2021), knowing the dominant anatoxin congener in a river may better inform risk.

Benthic cyanobacterial dynamics were more dynamic relative to GPP than we originally hypothesized. While GPP estimates in our rivers were less productive than the nearby Klamath River (Genzoli and Hall 2025) but more productive than most rivers in a broader analysis of GPP in U.S. rivers (Savoy et al. 2019), GPP in each of our three rivers was overall more stable during our study period than benthic cyanobacterial dynamics. Focal benthic cyanobacterial taxa cover during our study ranged from nearly absent to >10% cover and anatoxin concentrations ranged from non-detection or low detections to concentrations greater than 80 μg anatoxins g^-1^ OM between two consecutive biweekly sampling dates. The relative consistency in GPP is likely due to the presence of other autotrophs in these rivers than can substantially contribute to GPP (e.g., diatoms; Carter et al. 2025) that were often more dominant and consistent in cover. However, GPP in the rivers we studied were typically more variable than net ecosystem productivity (NEP) as trends in GPP were often mimicked by those in ecosystem respiration (ER) which suggests that comparison of benthic cyanobacterial dynamics and NEP may be less informative (Figure S22).

Despite lack of strong relationships between benthic cyanobacterial dynamics and GPP, *Microcoleus* cover tended to increase during periods of relatively low GPP. In both years of sampling at the South Fork Eel River, we saw *Microcoleus* cover increase as GPP slowly declined from August until the end of the field season. This suggests that *Microcoleus* in the South Fork Eel River proliferates as other biomass senesces (as indicated by the decline in GPP) (Appendix S1: Figure S23). Conversely, from our single year of sampling at the Salmon River, *Microcoleus* cover preceded peak GPP in the Salmon River. Taken together, these results indicate that the controls on *Microcoleus* cover and GPP are dissimilar between these rivers (Figure 1b or 1c). Relationships between GPP and *Anabaena*/*Cylindrospermum* cover were weaker. There was some evidence that peak *Anabaena*/*Cylindrospermum* cover and GPP were relatively synchronous in the South Fork Eel River as the peak dates for each occurred within ten days of each other in 2022 and 2023 (Appendix S1: Figure S23). This suggests that the controls on GPP and *Anabaena*/*Cylindrospermum* cover may be more similar (Figure 1a) possibly due to the epiphytic nature of *Anabaena* which thereby associates it with higher autotrophic biomass; however, we did not observe this relationship in the Russian River. Moreover, anatoxin concentrations within mats of both focal taxa were even more unrelated.

### Q2: Variability of benthic cyanobacterial dynamics among reaches within the South Fork Eel River

Weekly patterns of taxon-specific benthic cyanobacterial cover among five reaches in the South Fork Eel River in 2023 were generally consistent across all reaches with the exception of one reach that had a key distinction from the others. For *Microcoleus*, the reach displaying contradicting cover patterns (SFE-Lower-4) lacked a comparably large riffle feature that was found in other reaches. Some studies have shown greater *Microcoleus* biomass accrual and persistence in regions with faster flow (Hart et al. 2013, McAllister et al. 2018b, 2020); hence, the lack of a large riffle at this reach could have contributed to its relatively low cover. Instead, *Microcoleus* peaked at this reach in mid-August during a brief 2-3-week period in which *Microcoleus* was found in pools growing as an epiphyte on *Cladophora* along with epiphytic *Anabaena* throughout our study reaches (Appendix S1: Figure S24). Although most of our field observations support the importance of higher flows found in riffle habitats for *Microcoleus* proliferation (Hart et al. 2013), this observed epiphytic growth expanded its habitat.

For *Anabaena*/*Cylindrospermum*, the reach with atypical patterns (SFE-Lower-2) had much shallower pools which lacked *Cladophora* growth. As *Anabaena* is often found growing as an epiphyte on *Cladophora* (Bouma-Gregson et al. 2017, Kelly et al. 2019), it is unsurprising that this reach contained minimal *Anabaena* cover. Our single day of recorded *Anabaena*/*Cylindrospermum* cover at this reach was likely *Cylindrospermum* growing on fine sediment in the shallow pool; however, these taxa can be difficult to distinguish macroscopically. Despite the low recorded percent cover, we did observe many *Anabaena/Cylindrospermum* mats accumulating on the river banks at this reach. As we based our benthic surveys on the California SWAMP protocol (Ode et al. 2016) which only assesses cover at 25%, 50%, and 75% of the channel width and does not incorporate mats on river banks, we had a discrepancy between our records of transect presence/absence of our focal taxa and that from our quadrat cover surveys (Appendix S1: Figure S25). Future assessments of benthic cyanobacterial dynamics that incorporate cyanobacterial mats accumulating along river edges would more comprehensively record cyanobacterial abundances, especially as these shoreline mats are more accessible to humans and animals and could create a greater public health risk.

Despite the uncertainty surrounding *Anabaena* as an anatoxin producer (Kust et al. 2018, Kelly et al. 2019), we detected comparable anatoxin concentrations in *Anabaena*/*Cylindrospermum* and *Microcoleus* samples (both up to >150 μg anatoxins g^-1^ OM) from the same river as observed previously in the Eel River watershed (Bouma-Gregson et al. 2018). It is possible that other toxin-producers are responsible for the high levels of anatoxin observed in our *Anabaena*/*Cylindrospermum* composite samples as many samples also contained other taxa associated with anatoxin production including *Microcoleus* and *Geitlerinema* (Appendix S1: Figure S3). Other studies have also observed high anatoxin concentrations in mats dominated by the green alga *Cladophora* (Kelly et al. 2019) or other cyanobacterial taxa (Genzoli et al. 2024) in which *Microcoleus* was subdominant, but molecular analyses of these samples have shown that the anaC genes present within these mats were from Oscillatoriales (likely *Microcoleus*). In addition, our highest anatoxin concentrations were observed in both 2022 and 2023 during a period in which *Microcoleus* grew as an epiphyte on *Cladophora* in tandem with *Anabaena* in mid-August. In a concurrent study completed by the North Coast Regional Water Quality Board that took individual mat samples weekly (NCRWQCB 2024), epiphytic *Microcoleus* mat samples contained from 10x-500x higher anatoxin concentrations than a *Microcoleus* mat sample taken from a cobble in a riffle on the same day (Appendix S1: Figure S24). Our reach-scale composite did not capture variation in anatoxins from mats at smaller spatial scales, but future studies could investigate anatoxin production potential of *Microcoleus* growing on different substrates to further understand public health risk.

While we observed comparable anatoxins in *Microcoleus* and *Anabaena*/*Cylindrospermum* samples, each taxon displayed a different relationship between anatoxin concentrations and cover within the same river. *Anabaena*/*Cylindrospermum* anatoxin concentrations were roughly synchronous with cover, while *Microcoleus* anatoxins peaked prior to maximum cover. Positive relationships between *Microcoleus* cover and anatoxin concentrations have been documented previously (Wood et al. 2017a) but are not common (McAllister et al. 2018a, Robichon et al. 2025). In our study, the hysteresis relationship between *Anabaena/Cylindrospermum* cover and anatoxin concentrations suggest that *Anabaena/Cylindrospermum* cover can equate with anatoxin risk (Figure 1b) in the South Fork Eel River and, to a lesser extent, in the Russian River (Appendix S1: Figure S26). In contrast, high *Microcoleus* cover is not necessarily a good indicator of high anatoxin concentrations (Figure 1c) as we saw both on the South Fork Eel and Salmon Rivers.

### Q3: Predictive Modeling within the South Fork Eel River

We had more success (as indicated by lower NRMSE) predicting *Microcoleus* cover than *Anabaena*/*Cylindrospermum* cover largely due to the negative relationship between *Microcoleus* cover and discharge during our study period. Other studies have found a negative relationship between discharge and benthic cyanobacterial cover (Robichon et al. 2023) and specifically *Microcoleus* (Heath et al. 2011, Wood et al. 2017a). Large increases in discharge can cause “flushing flows” which can remove *Microcoleus* from the river bed. The concept of this high discharge flushing flow has been used as the removal term in other studies and reports predicting *Microcoleus* occurrence and cover(Heath et al. 2011, Atalah et al. 2018, Thomson-Laing 2018) and in models predicting autotrophic biomass (Uehlinger et al. 1996, Blaszczak et al. 2023, Lowman et al. 2024). Otherwise, *Microcoleus* appears to accrue in the absence of flushing flows, as we saw in our study, which occurred during stable summer baseflows before the flushing flow from seasonal autumn and winter rains.

While GPP was a poor predictor of *Microcoleus* cover, the “biological” model incorporating GPP as a covariate was the best predictor of *Anabaena/Cylindrospermum* cover. Due to its epiphytic nature, the presence of *Anabaena* may mostly relate to within river succession of *Cladophora* which senesced prior to the end of the field season (Power 1992, Power et al. 2009). This would explain why the “biological” model incorporating GPP, which reflects total autotrophic production, had better predictive accuracy than models with physical or chemical covariates; though, this model still did not outperform the null model. Based on the poor performance and large predictive uncertainty of our models predicting *Anabaena/Cylindrospermum* cover, we do not have a clear understanding of what triggers the growth of *Anabaena*, as *Cladophora* and other green algae were present initially without *Anabaena*, nor what causes the sudden senescence of both. Overall, the results of our cover predictions suggest that the controls on benthic cyanobacterial cover are taxon-specific and may be more complex than the linear relationships explored by our models.

In contrast, we had more success predicting *Anabaena*/*Cylindrospermum* anatoxins than *Microcoleus* anatoxins, which was mostly attributable to the incorporation of taxon-specific cover as a covariate. This further supports the synchronous relationship we observed in this study between *Anabaena*/*Cylindrospermum* cover and anatoxin concentrations (Figure 4c, 4e), but not for *Microcoleus* (Figure 4b, 4d). Beyond the inclusion of cover as a covariate, models predicting anatoxins performed similarly, suggesting that none of our other abiotic covariates were strongly influential in the production of anatoxins. This was also reflected in the posterior estimates for our anatoxin models where no covariate (excluding cover) had a 95% posterior estimate that was consistently positive or negative and did not overlap zero (Appendix S1: Figure S20, S21).

Previous studies evaluating the relationships between anatoxins and abiotic predictors have also had limited success. As found in our study, temperature has been mostly shown to have a limited or no effect on *Microcoleus* anatoxins (Wood et al. 2017a, McAllister et al. 2018a, Robichon et al. 2025). Some studies have found an association between anatoxin concentrations and low flows (Heath et al. 2011, Robichon et al. 2025) which we saw in some, but not all, of our models as the 95% posterior estimate for discharge overlapped zero in other models. Wood et al. (2017a) noted a negative association between *Microcoleus* anatoxins and specific conductivity; however, we more frequently observed the opposite in our models predicting *Microcoleus* anatoxins where specific conductivity had an often highly positive posterior estimate. The other two covariates in “chemical” models, orthophosphate and DIN had posterior estimates alternating from positive to negative despite some evidence of higher anatoxin concentrations at depressed phosphorus concentrations and elevated nitrogen concentrations in prior studies (Heath et al. 2016, Wood et al. 2017a). As our study and many others do not control for the relative proportion of toxigenic and nontoxigenic strains in a sample (Wood and Puddick 2017), it may be hard to parse out what environmental controls potentially influence anatoxin production in the field. Furthermore, it may be possible that only finer scale processes such as a strain’s growth rate stage (Brown et al. 2025) or nitrogen limitation (Stancheva et al. 2025) explain anatoxin production rather than the broader covariates we measured, or that nonlinear methods may be necessary (Thomson-Laing et al. 2021). These findings were also highlighted in our uncertainty analyses which revealed that process uncertainty was the dominant contributor to predictive uncertainty in all of our models. As is often the case with ecological forecasts (Dietze et al. 2017), including forecasts of planktonic cyanobacterial density (Lofton et al. 2022), the lack of understanding of the process was our biggest limit to forecasting benthic cyanobacterial dynamics.

While some of our predictive modeling findings may not translate to other rivers, as we only focused on one river (Atalah et al. 2018), we found three general takeaways for future forecasting efforts of benthic cyanobacteria: (1) implementing frequent sampling (i.e., weekly), (2) creating models with geomorphologically comparable reaches, and (3) developing a more complete understanding of the processes controlling benthic cyanobacterial dynamics. The initial results of our 2022 field season in which we sampled three rivers biweekly (every other week) indicated that we were not fully capturing the rapidly changing benthic cyanobacterial cover and anatoxin concentrations. Our 2023 weekly sampling within the South Fork Eel River improved our ability to capture the quick changes of benthic cyanobacterial dynamics. This is particularly true for anatoxin concentrations, as the timing of peak concentrations can be easily missed if sampling is too sparse. We also found geomorphologically similar reaches to be important when predicting benthic cyanobacterial cover within a river, as habitats can influence the type of taxa present within a reach. As aforementioned, the lack of a large riffle in one of our reaches likely contributed to different temporal patterns in *Microcoleus* cover in this reach compared to the other four reaches. Consequently, the sub-model predicting this reach performed poorly, as it was built from reaches with habitats that likely allowed for the proliferation of more *Microcoleus* (Appendix S1: Figure S14). Lastly, as our uncertainty analyses demonstrated, process uncertainty was the dominate source of uncertainty in our predictions; therefore, a more complete understanding of the processes governing benthic cyanobacteria dynamics could reduce our predictive uncertainty and likely allow us to make more accurate forecasts.

## Conclusion

The prediction or forecasting of toxigenic benthic cyanobacteria populations, as well as the overall scientific understanding of the processes governing benthic cyanobacteria dynamics, lags behind that of planktonic cyanobacteria despite their significant public health threat (Rousso et al. 2020, Wood et al. 2020, Murphy et al. 2025). Our study evaluated GPP as a potential proxy for benthic cyanobacterial dynamics but was ultimately met with limited success as relationships between GPP and taxon-specific benthic cyanobacterial dynamics were unclear or varied among rivers. Instead, our predictive models built for reaches within the South Fork Eel River demonstrated the importance of taxon-specific patterns as the best models predicting each taxon’s dynamics varied. Overall, our study showcases the applicability of ecological forecasting to the public and ecological health threats posed by benthic cyanobacteria, as well as emphasizes the importance of accounting for specific predictors of each toxigenic benthic cyanobacteria taxon in future predictions or forecasts rather than treating toxigenic benthic cyanobacteria as a single population.

## III. Acknowledgements

This research was supported by the National Science Foundation’s Division of Environmental Biology (DEB: 2042915), Emerging Frontiers Office (URoL:EN: 2222322), and Graduate Research Fellowship Program. Additionally, Keith Bouma-Gregson was partially supported by the USGS-National Parks Service Water Quality Partnership Program. We thank the UC Angelo Coast Reserve, especially Peter Steel, the reserve manager, for hosting us while completing field work and Mary Power for hosting the 2022 Algal Foray right before this work began. We would also like the Karuk Natural Resources Department for permission and guidance for sampling on the Salmon River and access to additional field data. We thank Bob Hall for guidance with river metabolism modeling and initial conversations pertaining to this project. Lastly, we thank Helen Lei, Jasmine Krause, Leon Katona, Andrea Garcia-Jiménez, and Sam Struthers for lab and field assistance. Any use of trade, firm, or product names is for descriptive purposes only and does not imply endorsement by the U.S. Government.

## IV. Author Contributions

Following CRediT authorship guidelines: **Jordan M. Zabrecky:** Conceptualization, Data Curation, Formal Analysis, Investigation, Writing-Original Draft, Visualization, Writing-Reviewing and Editing. **Taryn A. Elliott**: Investigation, Writing-Reviewing and Editing. **Meghan Hickey**: Investigation, Writing-Reviewing and Editing. **Keith Bouma-Gregson:** Conceptualization, Writing-Reviewing and Editing. **Greg Boyer**: Resources, Investigation, Writing-Reviewing and Editing. **Rich Fadness:** Investigation, Writing-Reviewing and Editing. **Laurel Genzoli**: Conceptualization, Formal Analysis, Investigation, Writing-Reviewing and Editing. **Ramesh Goel:** Funding Acquisition, Writing-Reviewing and Editing. **Grant Johnson**: Data Curation, Supervision, Writing-Reviewing and Editing. **Robert Shriver:** Formal Analysis, Funding Acquisition, Writing-Reviewing and Editing. **Rosalina Stancheva**: Funding Acquisition, Supervision, Writing-Reviewing and Editing. **Michael Thomas**: Investigation, Writing-Reviewing and Editing. **Zac Triumph:** Investigation. Writing-Reviewing and Editing. **Joanna R. Blaszczak:** Conceptualization, Formal Analysis, Funding Acquisition, Resources, Supervision, Writing-Original Draft. Writing-Reviewing and Editing.

## References

Anderson, B., J. Voorhees, B. Phillips, R. Fadness, R. Stancheva, J. Nichols, D. Orr, and S. A. Wood. 2018. Extracts from benthic anatoxin-producing *Phormidium* are toxic to 3 macroinvertebrate taxa at environmentally relevant concentrations: Cyanobacteria toxicity to 3 invertebrates. Environmental Toxicology and Chemistry 37:2851–2859.

Appling, A. P., R. O. Hall, C. B. Yackulic, and M. Arroita. 2018. Overcoming Equifinality: Leveraging Long Time Series for Stream Metabolism Estimation. Journal of Geophysical Research: Biogeosciences 123:624–645.

Atalah, J., H. Rabel, G. Thomson-Laing, and S. Wood. 2018. Drivers of Phormidium blooms in Southland rivers and the development of a predictive model. Page 31 p. appendices. Cawthron Report No. 3196, Prepared for Environment Southland.

Bernhardt, E. S., J. B. Heffernan, N. B. Grimm, E. H. Stanley, J. W. Harvey, M. Arroita, A. P. Appling, M. J. Cohen, W. H. McDowell, R. O. Hall, J. S. Read, B. J. Roberts, E. G. Stets, and C. B. Yackulic. 2018. The metabolic regimes of flowing waters. Limnology and Oceanography 63.

Bernhardt, E. S., P. Savoy, M. J. Vlah, A. P. Appling, L. E. Koenig, R. O. Hall, M. Arroita, J. R. Blaszczak, A. M. Carter, M. Cohen, J. W. Harvey, J. B. Heffernan, A. M. Helton, J. D. Hosen, L. Kirk, W. H. McDowell, E. H. Stanley, C. B. Yackulic, and N. B. Grimm. 2022. Light and flow regimes regulate the metabolism of rivers. Proceedings of the National Academy of Sciences 119:e2121976119.

Blaszczak, J. R., C. B. Yackulic, R. K. Shriver, and R. O. Hall Jr. 2023. Models of underlying autotrophic biomass dynamics fit to daily river ecosystem productivity estimates improve understanding of ecosystem disturbance and resilience. Ecology Letters 26:1510–1522.

Borlestean, A., P. C. Frost, and D. L. Murray. 2015. A mechanistic analysis of density dependence in algal population dynamics. Frontiers in Ecology and Evolution 3.

Bouma-Gregson, K., R. M. Kudela, and M. E. Power. 2018. Widespread anatoxin-a detection in benthic cyanobacterial mats throughout a river network. PLOS ONE 13:e0197669.

Bouma-Gregson, K., M. E. Power, and M. Bormans. 2017. Rise and fall of toxic benthic freshwater cyanobacteria (*Anabaena* spp.) in the Eel river: Buoyancy and dispersal. Harmful Algae 66:79–87.

Brooks, S. P., and A. Gelman. 1998. General Methods for Monitoring Convergence of Iterative Simulations. Journal of Computational and Graphical Statistics 7:434–455.

Brown, S. M., J. R. Blaszczak, R. K. Shriver, R. C. Jones, A. Sohrab, R. Goel, G. L. Boyer, B. Wei, K. M. Manoylov, T. R. Nelson, J. M. Zabrecky, and R. Stancheva. 2025. Growth and anatoxin-a production of *Microcoleus* (Cyanobacteria) strains from streams in California, USA. Harmful Algae 144:102834.

Carey, C. C., W. M. Woelmer, M. E. Lofton, R. J. Figueiredo, B. J. Bookout, R. S. Corrigan, V. Daneshmand, A. G. Hounshell, D. W. Howard, A. S. L. Lewis, R. P. McClure, H. L. Wander, N. K. Ward, and R. Q. Thomas. 2022. Advancing lake and reservoir water quality management with near-term, iterative ecological forecasting. Inland Waters 12:107–120.

Carter, A. M., R. O. Hall Jr., R. Feijó-Lima, M. DeGrandpre, Q. Shangguan, and H. M. Valett. 2025. Algal assemblage drives patterns in ecosystem structure but not metabolism in a productive river. Ecology 106:e70262.

Christensen, V. G., and E. Khan. 2020. Freshwater neurotoxins and concerns for human, animal, and ecosystem health: A review of anatoxin-a and saxitoxin. Science of The Total Environment 736:139515.

Conklin, K. Y., R. Stancheva, T. G. Otten, R. Fadness, G. L. Boyer, B. Read, X. Zhang, and R. G. Sheath. 2020. Molecular and morphological characterization of a novel dihydroanatoxin-a producing *Microcoleus* species (cyanobacteria) from the Russian River, California, USA. Harmful Algae 93:101767.

De Cicco, L. A., R. M. Hirsch, D. Lorenz, J. D. Watkins, and M. Johnson. 2024. dataRetrieval: R packages for discovering and retrieving water data available from Federal hydrologic web services. U.S. Geological Survey, Reston, VA.

Dietze, M. C. 2017. Ecological Forecasting. Princeton University Press.

Dietze, M. C., A. Fox, L. M. Beck-Johnson, J. L. Betancourt, M. B. Hooten, C. S. Jarnevich, T. H. Keitt, M. A. Kenney, C. M. Laney, L. G. Larsen, H. W. Loescher, C. K. Lunch, B. C. Pijanowski, J. T. Randerson, E. K. Read, A. T. Tredennick, R. Vargas, K. C. Weathers, and E. P. White. 2018. Iterative near-term ecological forecasting: Needs, opportunities, and challenges. Proceedings of the National Academy of Sciences 115:1424–1432.

Diez-Chiappe, A., M. Á. Muñoz-Martín, S. Cirés, A. Quesada, and E. Perona. 2025. Protected high-mountain rivers harbor widespread toxic *Microcoleus*-dominated mats with distinct genetic profiles. Harmful Algae 149:102942.

Faassen, E. J., L. Harkema, L. Begeman, and M. Lurling. 2012. First report of (homo)anatoxin-a and dog neurotoxicosis after ingestion of benthic cyanobacteria in The Netherlands. Toxicon 60:378–384.

Fredrickson, A., A. Richter, K. A. Perri, and S. R. Manning. 2023. First Confirmed Case of Canine Mortality Due to Dihydroanatoxin-a in Central Texas, USA. Toxins 15:485.

Genzoli, L., and R. O. Hall. 2025. Linking aquatic vegetation structure with ecosystem metabolism throughout the Klamath River, California, USA. Ecological Applications 35:e70089.

Genzoli, L., R. O. Hall, T. G. Otten, G. S. Johnson, J. R. Blaszczak, and J. Kann. 2024. Benthic cyanobacterial proliferations drive anatoxin production throughout the Klamath River watershed, California, USA. Freshwater Science 43:307–324.

Gugger, M., S. Lenoir, C. Berger, A. Ledreux, J.-C. Druart, J.-F. Humbert, C. Guette, and C. Bernard. 2005. First report in a river in France of the benthic cyanobacterium *Phormidium favosum* producing anatoxin-a associated with dog neurotoxicosis. Toxicon 45:919–928.

Hall, R. O., and E. R. Hotchkiss. 2017. Stream Metabolism. Pages 219–233 Methods in Stream Ecology. Elsevier.

Hart, D. D., B. J. F. Biggs, V. I. Nikora, and C. A. Flinders. 2013. Flow effects on periphyton patches and their ecological consequences in a New Zealand river. Freshwater Biology 58:1588–1602.

Heath, M., S. A. Wood, R. G. Young, and K. G. Ryan. 2016. The role of nitrogen and phosphorus in regulating *Phormidium* sp. (cyanobacteria) growth and anatoxin production. FEMS Microbiology Ecology 92:fiw021.

Heath, M., S. Wood, and K. Ryan. 2011. Spatial and temporal variability in *Phormidium* mats and associated anatoxin-a and homoanatoxin-a in two New Zealand rivers. Aquatic Microbial Ecology 64:69–79.

Herschy, R. 1993. The velocity-area method. Flow Measurement and Instrumentation 4:7–10.

Huisman, J., G. A. Codd, H. W. Paerl, B. W. Ibelings, J. M. H. Verspagen, and P. M. Visser. 2018. Cyanobacterial blooms. Nature Reviews Microbiology 16:471–483.

Kaminski, A., B. Bober, Z. Lechowski, and J. Bialczyk. 2013. Determination of anatoxin-a stability under certain abiotic factors. Harmful Algae 28:83–87.

Kelly, L. T., D. G. Beach, J. R. Blaszczak, K. Bouma-Gregson, S. M. Brown, H. Cheng, J. L. Davidson, J. Fastner, M. Francis, A. G. Jimenez, L. Genzoli, R. Goel, D. Gonzalez, K. M. Handley, S. Hilt, J.-F. Humbert, R. Jamieson, L. Johnston, P. Junier, J. Lawrence, P. McCarron, S. Meissner, J. Mormando, J. Puddick, C. Quiblier, N. Rajpirathap, C. Schampera, A. Selwood, K. Shearer, A. Sohrab, R. Stancheva, C. Valadez-Cano, J. M. Zabrecky, and S. A. Wood. 2026. The global proliferation of aquatic, benthic *Microcoleus*: taxonomy, distribution, toxin production, ecology, and future directions. Water Research:125441.

Kelly, L. T., K. Bouma-Gregson, J. Puddick, R. Fadness, K. G. Ryan, T. W. Davis, and S. A. Wood. 2019. Multiple cyanotoxin congeners produced by sub-dominant cyanobacterial taxa in riverine cyanobacterial and algal mats. PLOS ONE 14:e0220422.

Kust, A., P. Urajová, P. Hrouzek, D. L. Vu, K. Čapková, L. Štenclová, K. Řeháková, E. Kozlíková-Zapomělová, O. Lepšová-Skácelová, A. Lukešová, and J. Mareš. 2018. A new microcystin producing *Nostoc* strain discovered in broad toxicological screening of non-planktic Nostocaceae (cyanobacteria). Toxicon 150:66–73.

Leopold, L. B., and T. Maddock. 1953. The hydraulic geometry of stream channels and some physiographic implications. Page 64. Report, Washington, D.C.

Lofton, M. E., J. A. Brentrup, W. S. Beck, J. A. Zwart, R. Bhattacharya, L. S. Brighenti, S. H. Burnet, I. M. McCullough, B. G. Steele, C. C. Carey, K. L. Cottingham, M. C. Dietze, H. A. Ewing, K. C. Weathers, and S. L. LaDeau. 2022. Using near-term forecasts and uncertainty partitioning to inform prediction of oligotrophic lake cyanobacterial density. Ecological Applications 32:e2590.

Lofton, M. E., D. W. Howard, R. Q. Thomas, and C. C. Carey. 2023. Progress and opportunities in advancing near-term forecasting of freshwater quality. Global Change Biology 29:1691–1714.

Lowman, H. E., R. K. Shriver, R. O. Hall, J. W. Harvey, P. Savoy, C. B. Yackulic, and J. R. Blaszczak. 2024. Macroscale controls determine the recovery of river ecosystem productivity following flood disturbances. Proceedings of the National Academy of Sciences 121:e2307065121.

Loza, V., E. Perona, J. Carmona, and P. Mateo. 2013. Phenotypic and genotypic characteristics of *Phormidium*-like cyanobacteria inhabiting microbial mats are correlated with the trophic status of running waters. European Journal of Phycology 48:235–252.

McAllister, T. G., S. A. Wood, J. Atalah, and I. Hawes. 2018a. Spatiotemporal dynamics of Phormidium cover and anatoxin concentrations in eight New Zealand rivers with contrasting nutrient and flow regimes. Science of The Total Environment 612:71–80.

McAllister, T. G., S. A. Wood, M. J. Greenwood, F. Broghammer, and I. Hawes. 2018b. The effects of velocity and nitrate on *Phormidium* accrual cycles: a stream mesocosm experiment. Freshwater Science 37:496–509.

McAllister, T. G., S. A. Wood, and I. Hawes. 2016. The rise of toxic benthic Phormidium proliferations: A review of their taxonomy, distribution, toxin content and factors regulating prevalence and increased severity. Harmful Algae 55:282–294.

McAllister, T. G., S. A. Wood, E. M. MacKenzie, and I. Hawes. 2020. Reach- and mat-scale differences in Microcoleus autumnalis (cyanobacterium) accrual along velocity and nitrate gradients in three New Zealand rivers. Canadian Journal of Fisheries and Aquatic Sciences 77:401–412.

McCarron, P., C. Rafuse, S. Scott, J. Lawrence, M. R. Bruce, E. Douthwright, C. Murphy, M. Reith, and D. G. Beach. 2023. Anatoxins from benthic cyanobacteria responsible for dog mortalities in New Brunswick, Canada. Toxicon 227:107086.

McKay, L., T. Bondelid, J. Johnston, R. Moore, and A. Rea. 2012. NHDPlus Version 2: User Guide.

Méjean, A., K. Dalle, G. Paci, S. Bouchonnet, S. Mann, V. Pichon, and O. Ploux. 2016. Dihydroanatoxin-a Is Biosynthesized from Proline in *Cylindrospermum stagnale* PCC 7417: Isotopic Incorporation Experiments and Mass Spectrometry Analysis. Journal of Natural Products 79:1775–1782.

Murphy, J. C., R. M. Gorney, L. A. Lucas, and J. A. Zwart. 2025. A systematic literature review of forecasting and predictive models of harmful algal blooms in flowing waters. Preprint.

NCRWQCB. 2022. Benthic Cyanobacteria and Cyanotoxin Monitoring in Northern California Rivers, 2016-2019. Freshwater Harmful Algal Bloom Monitoring and Response Program, North Coast Regional Water Quality Control Board, Santa Rosa, CA.

NCRWQCB. 2024. Implementation of a Benthic Cyanobacteria Tiered Monitoring Program for Public Health Protection in Northern California Rivers. Freshwater Harmful Algal Bloom Monitoring and Response Program, North Coast Regional Water Quality Control Board, Santa Rosa, CA.

Ode, P. R., A. E. Fetscher, and Busse, Lilian B. 2016. Standard Operating Procedures (SOP) for the Collection of Field Data for Bioassessments of California Wadeable Streams: Benthic Macroinvertebrates, Algae, and Physical Habitat. Technical Report, California Water Board Surface Water Ambient Monitoring Program.

Odum, H. T. 1956. Primary Production in Flowing Waters. Limnology and Oceanography 1:102– 117.

PME. 2021. miniDOT Logger User’s Manual. PME, Vista, CA.

Power, M. E. 1992. Hydrologic and trophic controls of seasonal algal blooms in northern California rivers. Archiv für Hydrobiologie 125:385–410.

Power, M. E., S. Chandra, P. Gleick, and W. E. Dietrich. 2024. Anticipating responses to climate change and planning for resilience in California’s freshwater ecosystems. Proceedings of the National Academy of Sciences 121:e2310075121.

Power, M., R. Lowe, P. Furey, J. Welter, M. Limm, J. Finlay, C. Bode, S. Chang, M. Goodrich, and J. Sculley. 2009. Algal mats and insect emergence in rivers under Mediterranean climates: towards photogrammetric surveillance. Freshwater Biology 54:2101–2115.

Puddick, J., R. Van Ginkel, C. D. Page, J. S. Murray, H. E. Greenhough, J. Bowater, A. I. Selwood, S. A. Wood, M. R. Prinsep, P. Truman, R. Munday, and S. C. Finch. 2021. Acute toxicity of dihydroanatoxin-a from *Microcoleus autumnalis* in comparison to anatoxin-a. Chemosphere 263:127937.

Puschner, B., B. Hoff, and E. R. Tor. 2008. Diagnosis of Anatoxin-a Poisoning in Dogs from North America. Journal of Veterinary Diagnostic Investigation 20:89–92.

Quiblier, C., S. Wood, I. Echenique-Subiabre, M. Heath, A. Villeneuve, and J.-F. Humbert. 2013. A review of current knowledge on toxic benthic freshwater cyanobacteria--ecology, toxin production and risk management. Water Research 47:5464–5479.

Raymond, P. A., C. J. Zappa, D. Butman, T. L. Bott, J. Potter, P. Mulholland, A. E. Laursen, W. H. McDowell, and D. Newbold. 2012. Scaling the gas transfer velocity and hydraulic geometry in streams and small rivers. Limnology and Oceanography: Fluids and Environments 2:41–53.

Ricker, W. E. 1954. Stock and Recruitment. Journal of the Fisheries Research Board of Canada 11:559–623.

Robichon, C., D. Latour, F. Perrière, S. Dolédec, and J. Robin. 2025. Determinism of anatoxin-a production in *Phormidium*-dominated biofilms of a regulated river (Ain, France). Harmful Algae 148:102912.

Robichon, C., J. Robin, and S. Dolédec. 2023. Relative effect of hydraulics, physico-chemistry and other biofilm algae on benthic cyanobacteria assemblages in a regulated river. Science of The Total Environment 872:162142.

Rodell, M., P. R. Houser, U. Jambor, J. Gottschalck, K. Mitchell, C.-J. Meng, K. Arsenault, B. Cosgrove, J. Radakovich, M. Bosilovich, J. K. Entin, J. P. Walker, D. Lohmann, and D. Toll. 2004. The Global Land Data Assimilation System. Bulletin of the American Meteorological Society 85.

Rousso, B. Z., E. Bertone, R. Stewart, and D. P. Hamilton. 2020. A systematic literature review of forecasting and predictive models for cyanobacteria blooms in freshwater lakes. Water Research 182:115959.

Sartory, D. P., and J. U. Grobbelaar. 1984. Extraction of chlorophyll a from freshwater phytoplankton for spectrophotometric analysis. Hydrobiologia 114:177–187.

Savoy, P., A. P. Appling, J. B. Heffernan, E. G. Stets, J. S. Read, J. W. Harvey, and E. S. Bernhardt. 2019. Metabolic rhythms in flowing waters: An approach for classifying river productivity regimes. Limnology and Oceanography 64:1835–1851.

Schaeffer, B. A., E. Urquhart, M. Coffer, W. Salls, R. P. Stumpf, K. A. Loftin, and P. Jeremy Werdell. 2022. Satellites quantify the spatial extent of cyanobacterial blooms across the United States at multiple scales. Ecological Indicators 140:108990.

Smith, Z. J., D. E. Conroe, K. L. Schulz, and G. L. Boyer. 2020. Limnological Differences in a Two-Basin Lake Help to Explain the Occurrence of Anatoxin-a, Paralytic Shellfish Poisoning Toxins, and Microcystins. Toxins 12:559.

Sohrab, A., S. Kaiser, B. Bhattarai, R. Stancheva, and R. Goel. 2025. The ecology of cyanobacteria and their synergism with bacterioplankton in benthic mats under nutrient limitations in the Virgin River in Zion’s National Park. Science of The Total Environment 997:180194.

Stancheva, R., S. Brown, G. L. Boyer, B. Wei, R. Goel, S. Henry, N. V. Kristan, and B. Read. 2025. Effect of salinity stress and nitrogen depletion on growth, morphology and toxin production of freshwater cyanobacterium Microcoleus anatoxicus Stancheva & Conklin. Hydrobiologia 852:561–574.

Strunecký, O., J. Komárek, J. Johansen, A. Lukešová, and J. Elster. 2013. Molecular and morphological criteria for revision of the genus *Microcoleus* (Oscillatoriales, Cyanobacteria). Journal of Phycology 49:1167–1180.

Thomson-Laing, G. 2018. Phormidium in the Maitai River: a review of current knowledge and the development of a predictive model. Page 53. Cawthron Report, Prepared for Nelson City Council.

Thomson-Laing, G., N. Dyer, R. Whyte-Wilding, and S. A. Wood. 2021. In situ river experiments to explore variability in *Microcoleus autumnalis* mat expansion. Hydrobiologia 848:445–467.

Toporowska, M., B. Pawlik-Skowrońska, and R. Kalinowska. 2014. Accumulation and effects of cyanobacterial microcystins and anatoxin-a on benthic larvae of *Chironomus* spp. (Diptera: Chironomidae). European Journal of Entomology 111:83–90.

Uehlinger, U., H. Bührer, and P. Reichert. 1996. Periphyton dynamics in a floodprone prealpine river: evaluation of significant processes by modelling. Freshwater Biology 36:249–263.

U.S. EPA. 2015. Method 545: Determination of Cylindrospermopsin and Anatoxin-a in Drinking Water by Liquid Chromatography Electrospray Ionization Tandem Mass Spectrometry (LC/ESI-MS/MS). Cincinnati, OH.

U.S. Geological Survey. 2021. Watershed Boundary Dataset.

U.S. Geological Survey. 2025. USGS Water Data for the Nation: U.S. Geological Survey National Water Information System database.

Vadeboncoeur, Y., and M. E. Power. 2017. Attached Algae: The Cryptic Base of Inverted Trophic Pyramids in Freshwaters. Annual Review of Ecology, Evolution, and Systematics 48:255–279.

Wood, S. A., J. Atalah, A. Wagenhoff, L. Brown, K. Doehring, R. G. Young, and I. Hawes. 2017a. Effect of river flow, temperature, and water chemistry on proliferations of the benthic anatoxin-producing cyanobacterium *Phormidium*. Freshwater Science 36:63–76.

Wood, S. A., L. Biessy, and J. Puddick. 2018. Anatoxins are consistently released into the water of streams with Microcoleus autumnalis-dominated (cyanobacteria) proliferations. Harmful Algae 80:88–95.

Wood, S. A., M. W. Heath, J. Kuhajek, and K. G. Ryan. 2010. Fine-scale spatial variability in anatoxin-a and homoanatoxin-a concentrations in benthic cyanobacterial mats: implication for monitoring and management. Journal of Applied Microbiology 109:2011– 2018.

Wood, S. A., L. T. Kelly, K. Bouma-Gregson, J. Humbert, H. D. Laughinghouse, J. Lazorchak, T. G. McAllister, A. McQueen, K. Pokrzywinski, J. Puddick, C. Quiblier, L. A. Reitz, K. G. Ryan, Y. Vadeboncoeur, A. Zastepa, and T. W. Davis. 2020. Toxic benthic freshwater cyanobacterial proliferations: Challenges and solutions for enhancing knowledge and improving monitoring and mitigation. Freshwater Biology 65:1824–1842.

Wood, S. A., J. Puddick, R. Fleming, and A. H. Heussner. 2017b. Detection of anatoxin-producing *Phormidium* in a New Zealand farm pond and an associated dog death. New Zealand Journal of Botany 55:36–46.

Wood, S. A., F. M. J. Smith, M. W. Heath, T. Palfroy, S. Gaw, R. G. Young, and K. G. Ryan. 2012. Within-Mat Variability in Anatoxin-a and Homoanatoxin-a Production among Benthic Phormidium (Cyanobacteria) Strains. Toxins 4:900–912.

Wood, S., and J. Puddick. 2017. The Abundance of Toxic Genotypes Is a Key Contributor to Anatoxin Variability in Phormidium-Dominated Benthic Mats. Marine Drugs 15:307.

Wurtsbaugh, W. A., H. W. Paerl, and W. K. Dodds. 2019. Nutrients, eutrophication and harmful algal blooms along the freshwater to marine continuum. WIREs Water 6:e1373.

Xia, Y., K. Mitchell, M. Ek, J. Sheffield, B. Cosgrove, E. Wood, L. Luo, C. Alonge, H. Wei, J. Meng, B. Livneh, D. Lettenmaier, V. Koren, Q. Duan, K. Mo, Y. Fan, and D. Mocko. 2012. Continental-scale water and energy flux analysis and validation for the North American Land Data Assimilation System project phase 2 (NLDAS-2): 1. Intercomparison and application of model products. Journal of Geophysical Research: Atmospheres 117.

Yates, K. L., P. J. Bouchet, M. J. Caley, K. Mengersen, C. F. Randin, S. Parnell, A. H. Fielding, A. J. Bamford, S. Ban, A. M. Barbosa, C. F. Dormann, J. Elith, C. B. Embling, G. N. Ervin, R. Fisher, S. Gould, R. F. Graf, E. J. Gregr, P. N. Halpin, R. K. Heikkinen, S. Heinänen, A. R. Jones, P. K. Krishnakumar, V. Lauria, H. Lozano-Montes, L. Mannocci, C. Mellin, M. B. Mesgaran, E. Moreno-Amat, S. Mormede, E. Novaczek, S. Oppel, G. Ortuño Crespo, A. T. Peterson, G. Rapacciuolo, J. J. Roberts, R. E. Ross, K. L. Scales, D. Schoeman, P. Snelgrove, G. Sundblad, W. Thuiller, L. G. Torres, H. Verbruggen, L. Wang, S. Wenger, M. J. Whittingham, Y. Zharikov, D. Zurell, and A. M. M. Sequeira. 2018. Outstanding Challenges in the Transferability of Ecological Models. Trends in Ecology & Evolution 33:790–802.

Zabrecky, J. M., T. A. Elliott, M. Hickey, H. Lei, R. S. Christova, G. Boyer, L. Genzoli, G. Johnson, and J. R. Blaszczak. 2025. Anatoxin concentrations, algal assemblages, and water quality data for the South Fork Eel, Salmon, and Russian Rivers in northern California, 2022-2023. Environmental Data Initiative. ver 3.

