## Appendix S1 for "Predicting taxon-specific benthic cyanobacterial mat cover and anatoxin concentrations in northern California rivers"

**Supporting Information**
**Appendix S1**

Predicting taxon-specific benthic cyanobacterial mat cover and anatoxin concentrations in northern California rivers

Jordan M. Zabrecky[1*], Taryn A. Elliott[1], Meaghan Hickey[1], Keith Bouma-Gregson[2], Gregory L. Boyer[3], Rich Fadness[4], Laurel Genzoli[1], Ramesh Goel[5], Grant Johnson[6], Robert Shriver[1], Rosalina Stancheva[7], Michael Thomas[4], Zac Triumph[3], Joanna R. Blaszczak[1]

[1]Department of Natural Resources and Environmental Science, University of Nevada, Reno, NV, USA
[2]U.S. Geological Survey California Water Science Center, Sacramento, CA, USA
[3]Department of Chemistry, State University of New York College of Environmental Science and Forestry, Syracuse, NY, USA
[4]North Coast Regional Water Quality Control Board, Santa Rosa, CA, USA
[5]Department of Civil and Environmental Engineering, University of Utah, Salt Lake City, UT, USA
[6]Department of Natural Resources, Karuk Tribe, Orleans, CA, USA
[7]Department of Environmental Science and Policy, George Mason University, Fairfax, VA, USA

Any use of trade, firm, or product names is for descriptive purposes only and does not imply endorsement by the U.S. Government.

**Section S1.** Supporting method for estimation of dissolved oxygen residence distances upstream of sensors.

We estimated the dissolved oxygen residence distance upstream of each sensor placement. This residence distance refers to the average distance in which dissolved oxygen travels before leaving the river by equilibrating with the atmosphere. Within this distance, a river is assumed to be relatively homogenous for single-station metabolism estimation given there are not any major discontinuities in river character. We calculated this distance as the distance in which 95% of the dissolved oxygen has equilibrated with the atmosphere using the equation $3v/K$ where $v$ is daily average stream velocity (m $d^{-1}$) and $K$ is the gas exchange rate ($d^{-1}$) (Hall and Hotchkiss 2017). To obtain velocity, we created a velocity-discharge relationship with U.S. Geological Survey (USGS) gage data downloaded with the "dataRetrieval" package (De Cicco et al. 2024) and used the velocity at median discharge of our study period for each site. For $K$, we used the median $O_2$ gas exchange rate ($K_{600}$) estimate obtained from metabolism modeling (see Appendix S1: Section S6). All of our survey reaches were located upstream of the associated sensor within this calculated distance and there were no major discontinuities in river character from the survey reaches to sensor placement (Appendix S1: Table S3).

**Section S2.** Supporting methods for microscopy to confirm the presence of *Microcoleus* or *Anabaena*/*Cylindrospermum* in collected cyanobacteria samples.

Microscopy was performed on all preserved composite samples to confirm presence of *Microcoleus* and *Anabaena* or *Cylindrospermum* using an Olympus BX40 compound microscope (Olympus Corporation, Shinjuku, Japan). All 2022 analyses were completed on preserved samples; however, for some samples in 2023, we analyzed live samples that were within ten days of collection. We prepared three slides per sample. The sample was transferred to a petri dish with three distinct markings on the underside where an aliquot of the sample was withdrawn and used to prepare three unbiased slides. We analyzed each slide using the transect approach wherein the entirety of the slide is viewed from left to right. We recorded percent abundance of non-algal material, green algae (filamentous and non-filamentous), diatoms (separating out *Epithemia* from all other diatoms), coccoid cyanobacteria, and each genus of filamentous cyanobacteria (e.g., *Microcoleus*, *Anabaena*, *Nostoc*, etc.). We limited analyses to ten minutes per slide. For both *Microcoleus* and *Anabaena*/*Cylindrospermum* samples, we were able to confirm their respective presence in all samples (Appendix S1: Figure S2). We calculated relative abundance as the average for each group across all three analyzed slides. We then recalculated relative abundances excluding the nonalgal material to determine the relative abundance of algal groups (Appendix S1: Figure S3).

**Section S3**. Supporting method for determination of chlorophyll-*a* in lyophilized composite benthic cyanobacterial samples.

We used a Trilogy Laboratory Fluorometer (Turner Designs, San Jose, CA) to measure chlorophyll-*a* of each lyophilized *Microcoleus* and *Anabaena*/*Cylindrospermum* composite sample. 2.0 ± 0.5 mg of lyophilized material was measured and suspended in 10 mL of 90% ethanol and heated to 78°C for five minutes to extract chlorophyll-*a* (Sartory and Grobbelaar 1984). Once cooled, samples were diluted (in order to be within instrument detection limits) in a 2 mL cuvette with 90% ethanol before analyzing and then acidified with 150 µL of 0.1 N HCl to obtain both chlorophyll-*a* and pheophytin concentrations. Detection limits for chlorophyll-*a* were 25 µg/L.

**Section S4:** Supporting methods and results for microcystins and cylindrospermopsins analyses.

In addition to anatoxin analyses, extracted samples were also analyzed for cylindrospermopsins and microcystins. Samples were analyzed by the Boyer Lab at State University of New York College of Environmental Science and Forestry (SUNY ESF) using a

Waters ACQUITY TQD mass spectrometer coupled with a Waters ACQUITY UPLC solvent delivery system using a 2.0 x 150 mm Ace C-18 column (Waters, Milford, MA) for cylindrospermopsins and a Waters 2695 solvent delivery system coupled to a 2996 photodiode array detector and ZQ4000 mass single quad mass spectrometer for microcystins, both following previously published methods (U.S. EPA 2015, Boyer 2020). Cylindrospermopsins analyses included three derivatives: cylindrospermopsin, epi-cylindrospermopsin, and deoxy-cylindrospermopsin. Instrument detection limits were calculated based on standards and method detection limits were calculated for each sample individually based on the instrument detection limit, sample weight, and extractant volume. Method detection limits ranged from 0.0029 to 0.1837 μg cylindrospermopsin per g lyophilized sample and 0.06 to 0.21 μg microcystins per g lyophilized sample.

We did not detect cylindrospermopsins in any of our samples; however, we detected quantifiable microcystins in three *Microcoleus* composite samples and four *Anabaena/Cylindrospermum* composite samples (Appendix S1: Table S5). One *Microcoleus* composite sample was collected from the South Fork Eel River in 2022 (0.73 μg microcystins per g lyophilized sample) and the other two were collected from the Salmon River in 2022 (1.08 and 1.37 μg microcystins per g lyophilized sample). All *Anabaena/Cylindrospermum* composite samples with detectable microcystins were collected from the South Fork Eel River in 2023 (range: 0.35-0.66 μg microcystins per g lyophilized sample).

**Section S5.** Supporting methods for analyses of surface water samples for orthophosphate and dissolved inorganic nitrogen (DIN).

Surface water samples were analyzed for orthophosphate ($PO_4^{3-}$), ammonium ($NH_4$), and nitrate ($NO_3$) using a SEAL AQ400 discrete analyzer (SEAL Analytical, Mequon, WI) at the University of Nevada, Reno following U.S. EPA methods 365.1 revision 2.0, 350.1 revision 2.0, and 353.2 revision 2.0, respectively (U.S. EPA 1993a, 1993b, 1993c). As our water samples were at a pH >7 at time of sampling, we calculated the proportions of ammonia-based nitrogen ($NH_3$) and ammonium-based nitrogen ($NH^{4+}$) using an equation to incorporate the influence of temperature and pH (Emerson et al. 1975). Detection limits were 0.402 μg P/L for orthophosphate, 0.002 mg N/L for ammonium, and 0.003 mg N/L for nitrate. We calculated dissolved inorganic nitrogen (DIN) by summing ammonium and nitrate. For samples with concentrations below detection limits, we used half the detection limit as the sample concentration in predictive modeling.

**Section S6.** Supporting methods for dissolved oxygen sensor maintenance and calibration and estimation of gross primary productivity (GPP).

After each summer of sampling, we completed a miniDOT dissolved oxygen sensor (Precision Measuring Engineering, Vista, CA) calibration using the bucket calibration method described by Precision Measurement Engineering (PME 2021), which involved placing sensors in a bucket of cold water with a bubbler to obtain 100% oxygen saturation. We calculated the theoretical dissolved oxygen (DO) concentration at 100% oxygen saturation using the water temperature and local pressure with an empirically-derived equation (Garcia and Gordon 1992) and calculated the offset between this theoretical DO concentration and the measured DO concentration for each sensor. We then applied these offsets to our DO data from the field prior to metabolism modeling. A calibration was not completed for the sensor deployed at the Russian River as it was stolen before final retrieval; however, we ultimately used DO data from the USGS to model metabolism for that river.

To prepare data for metabolism modeling, we removed outliers and brief (<6 hour) abnormalities or in our DO data occurring only during periods when DO linearly increased or decreased and linearly interpolated them using the "zoo" package in R (Zeileis and Grothendieck 2005). Longer periods (>6 hours) of abnormalities (e.g., biofouling identified via increasing amplitudes that halted after sensor cleaning) were removed and not replaced. As we identified biofouling on our sensors placed in the Russian and Salmon Rivers, we also modeled metabolism for those sites using publicly available data from the USGS (U.S. Geological Survey 2025) and proprietary data with permission from the Karuk Tribe and processed it as above. The dissolved oxygen sensor maintained by the Karuk Tribe was located where we placed our dissolved oxygen sensor. However, the USGS dissolved oxygen sensor at the Russian River in Cloverdale, CA (gage no: 11463000) was located ~10.6 km upstream of our DO sensor placed at the most downstream reach (RUS-1S), though this was still within our estimated dissolved oxygen residence distance for this site (Table S3).

To overcome the coarseness of Global Land Data Assimilation System (GLDAS) data (Rodell et al. 2004), we used GLDAS barometric pressure data to create a linear model between time and pressure. We then applied this model to more localized pressure measurements we took using an Extech SD 700 Datalogger (Extech Instruments, Nashua, NH) during each reach visit. We opted to not use the "StreamLight" package (Savoy et al. 2021) to make adjustments on North American Land Data Assimilation (NLDAS) light data as our rivers at summer baseflows meandered within a larger bankfull width from high winter flows, thus sometimes did not have any light shading from nearby trees to be considered.

To obtain a depth-discharge relationship, we kayaked within the dissolved oxygen footprint of our sensors at each site and obtained a minimum of eighty depth measurements. We obtained depths for the South Fork Eel River at both sites (SFE-Lower and SFE-Upper) on three occasions, each with three different discharge values, and created a linear model between log-transformed average kayak depth to log-transformed discharge (Leopold and Maddock 1953) to determine depth at various discharge values. For the Russian and Salmon Rivers, we were only able to obtain kayak depths at a single discharge value. Instead, we downloaded USGS channel geomorphology data at the gage using the "dataRetrieval" package (De Cicco et al. 2024), created a log-depth log-discharge relationship using this channel depth, and offset the depths by comparing the depth we obtained via kayak at the same discharge (Appendix S1: Figure S6).

We ran separate metabolism models for each site (grouping years together) in "streamMetabolizer" (Appling et al. 2018) with the inverse Bayesian model specification set to incorporate both observation and process error and to use the trapezoid rule to solve dissolved oxygen changes. We used partial pooling to constrain variability in modeled values of $K_{600}$ and adjusted the number and range of binning to be specific to the discharge of each site. We calculated $K_{600}$ priors for each site using an equation incorporating velocity, depth, and slope (Raymond et al. 2012; Appendix S1: Table S6). We created a velocity-discharge relationship with USGS gage data, used depths from our depth-discharge relationship, and obtained slopes from previous estimates at each river (Ayers Associates 1999, Foster and Kelsey 2012, Stillwater Sciences 2016). We used the velocity and depth at the median discharge of our study period for each site in the calculation. Metabolism models were run for differing burn-in steps and saved steps (ranging from 2,000 burn-in and 1,000 saved steps to 6,000 burn-in and 4,000 saved steps) to ensure convergence while maintaining efficiency. All Gelman-Rubin statistics (r-hats) for gross primary productivity (GPP), ecosystem respiration (ER), and $K_{600}$ for final estimates were below 1.05 (Appendix S1: Figure S7). Missing GPP values were interpolated linearly using the "zoo" package (Zeileis and Grothendieck 2005) before predictive modeling.

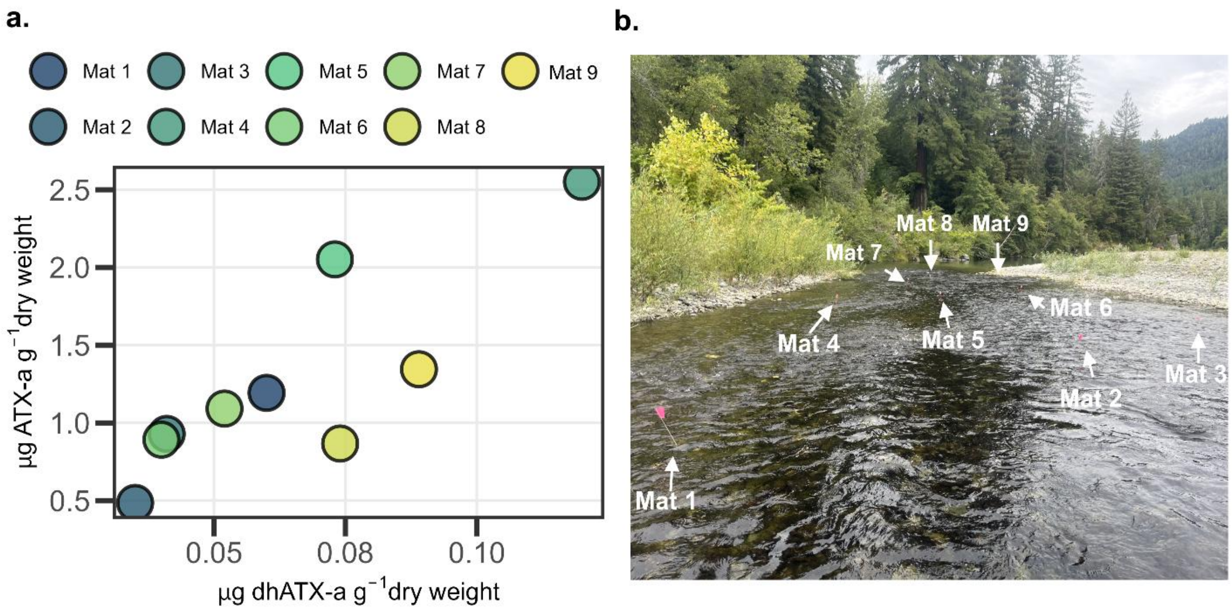


**Figure S1:** (a) Variation of dihydro-anatoxin-a (dhATX-a) and anatoxin-a (ATX-a) in nine individual mats each taken from a different cobble within the same riffle on the same day. (b) Photo of the riffle with the flags marking where each mat was taken. The riffle was located at reach SFE-Lower-3 and was approximately 15 m wide and 45 m long. Photo Credits: Jordan Zabrecky.

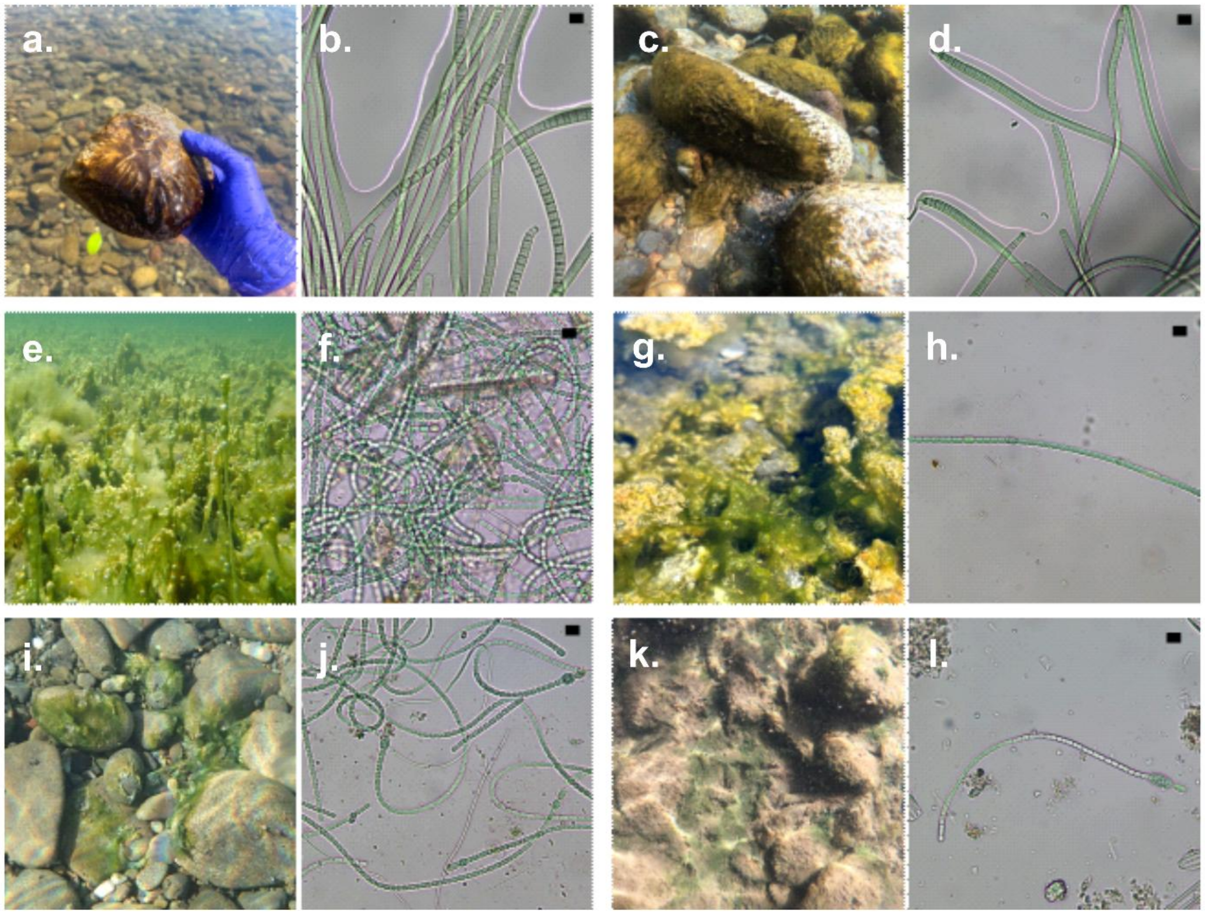


**Figure S2:** (a) Macrosopic and (b) microscopic *Microcoleus* from the South Fork Eel River. (c) Macroscopic and (d) microscopic *Microcoleus* from the Salmon River. (e) Macroscopic and (f) microscopic *Anabaena* from the South Fork Eel River. (g) Macroscopic and (h) microscopic *Anabaena* from the Russian River. (i) Macroscopic and (j) microscopic *Cylindrospermum* from the South Fork Eel River. (k) Macroscopic and (l) microscopic *Cylindrospermum* from the Russian River. All scale bars on microscopy photos are 10 µm. Photo credits: Jordan Zabrecky (a, g, k), Taryn Elliott (b, d, f, h, j, l), Joanna Blaszczak (c), and Andrea Garcia-Jiménez (e, i).

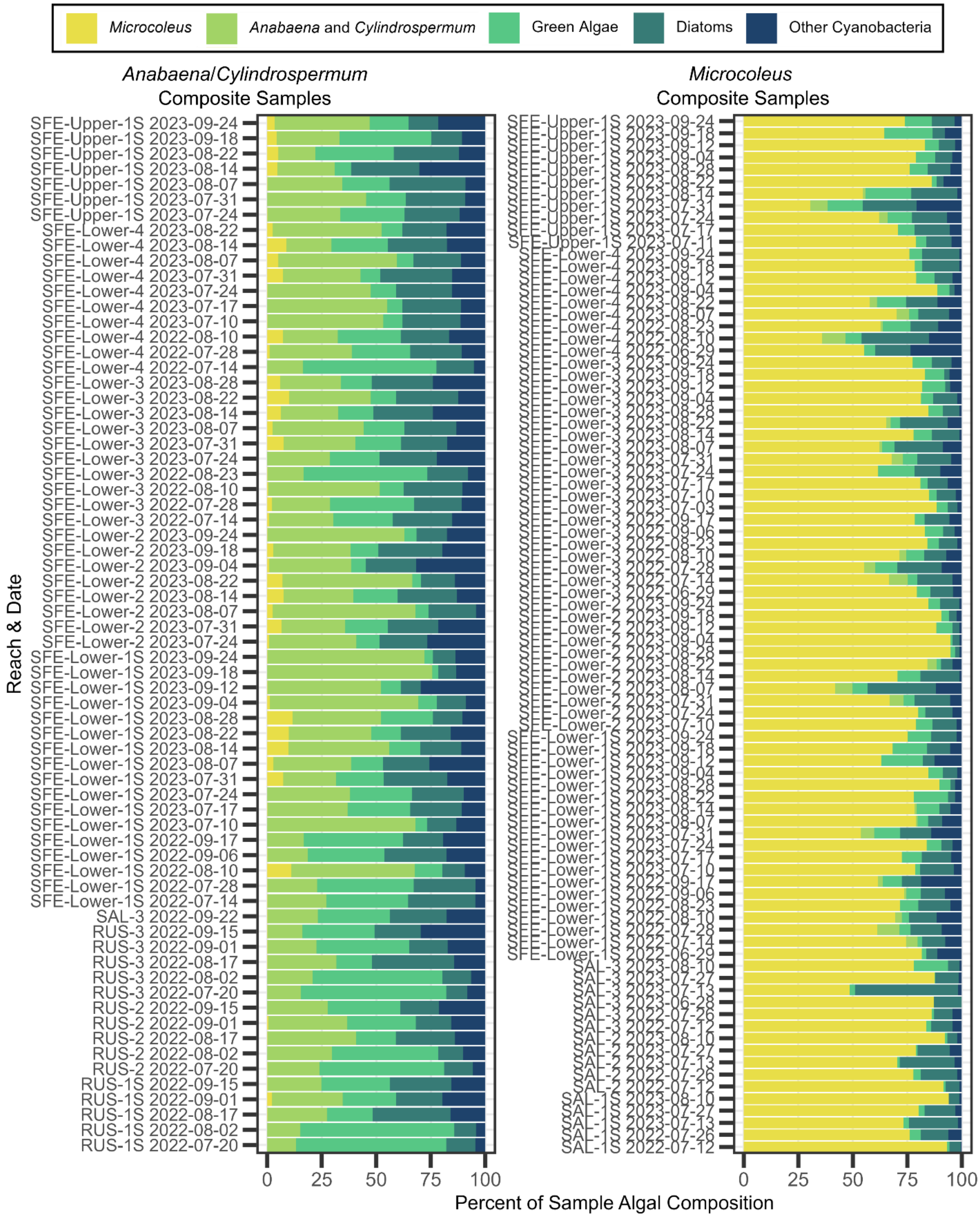


**Figure S3.** Algal composition of *Anabaena/Cylindrospermum* and *Microcoleus* composite samples as determined via microscopy. Sample names include the river abbreviation: "SFE" for South Fork Eel River, "SAL" for Salmon River, and "RUS" for Russian River.

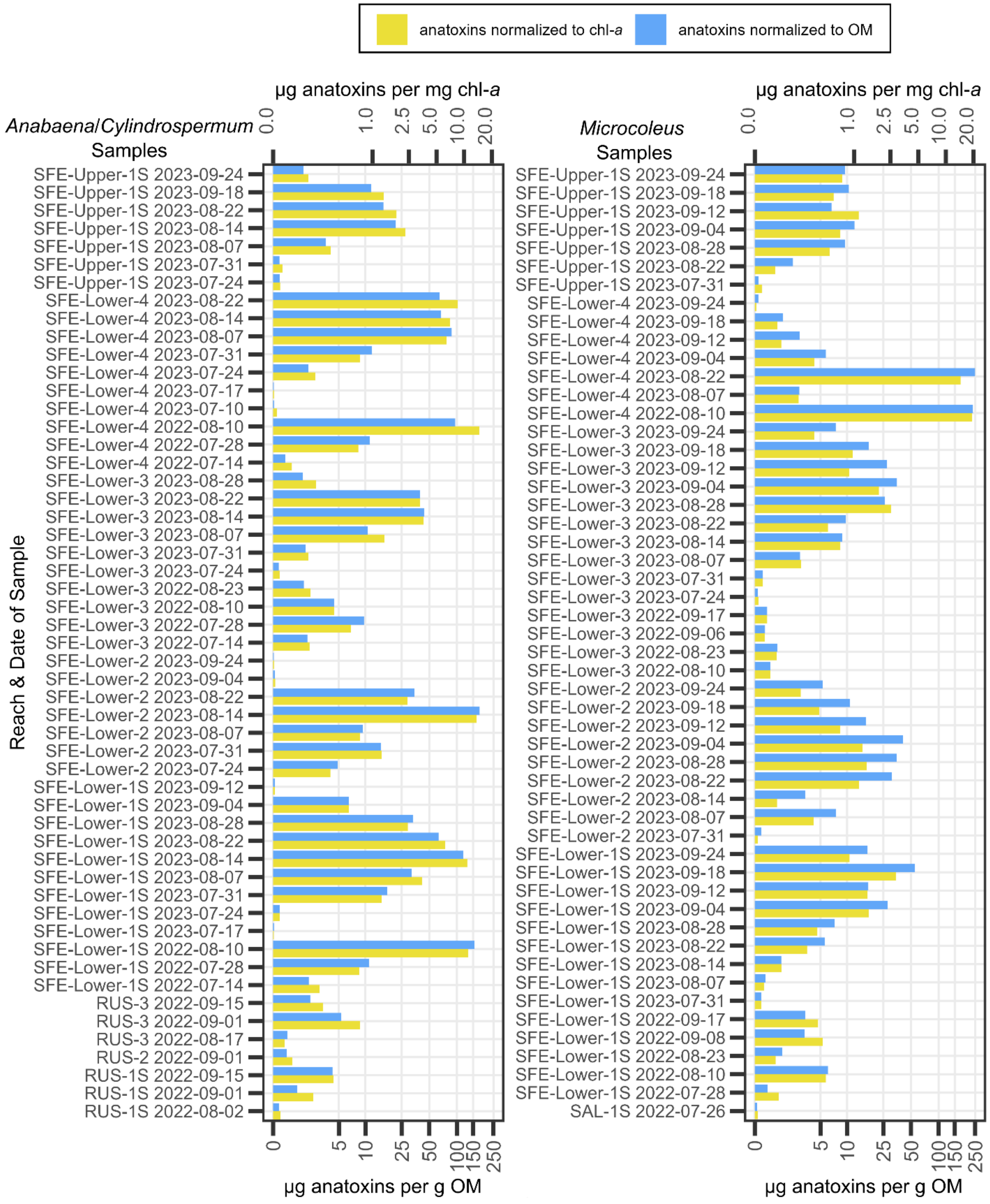


**Figure S4.** Total anatoxins concentrations (sum of all measured congeners) normalized to milligrams chlorophyll-*a* (chl-*a*) and grams organic matter (OM) for each sample.

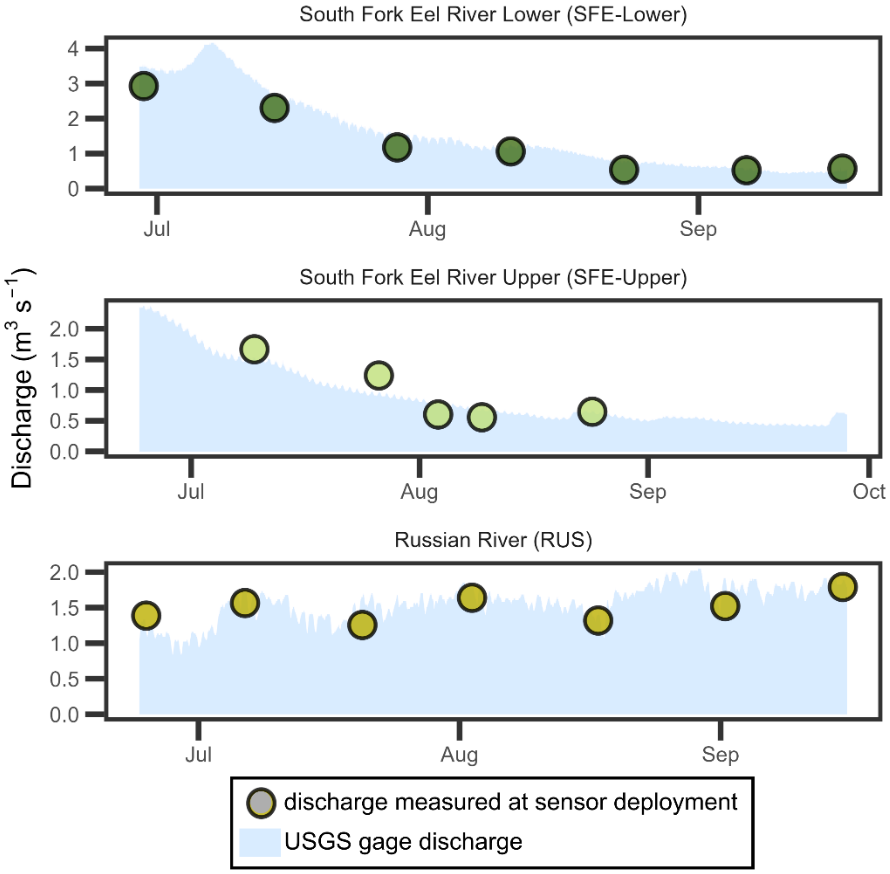


**Figure S5.** Discharge measured at sensor deployment locations versus discharge from nearby U.S. Geological Survey (USGS) gages.

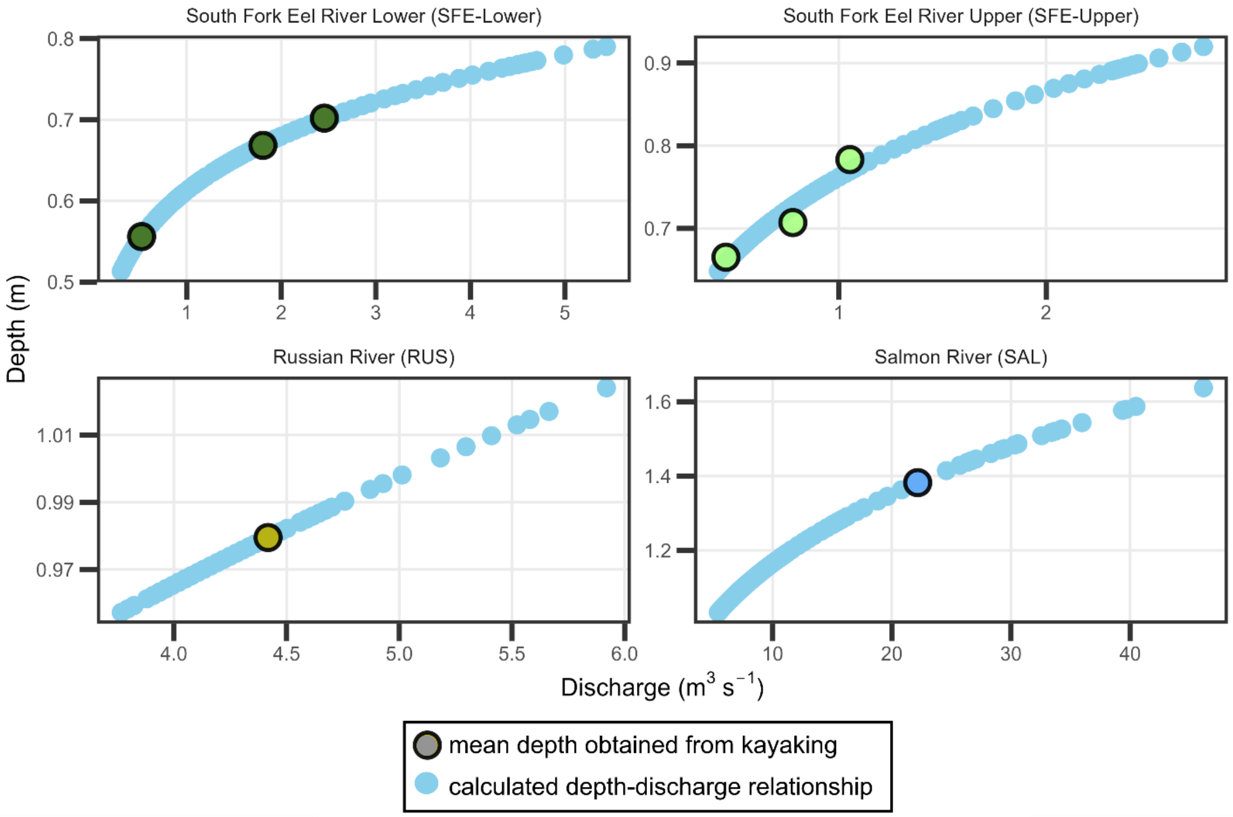


**Figure S6.** Depth-discharge relationships calculated for each sensor site and average measured depth obtained from kayaking at a given discharge.

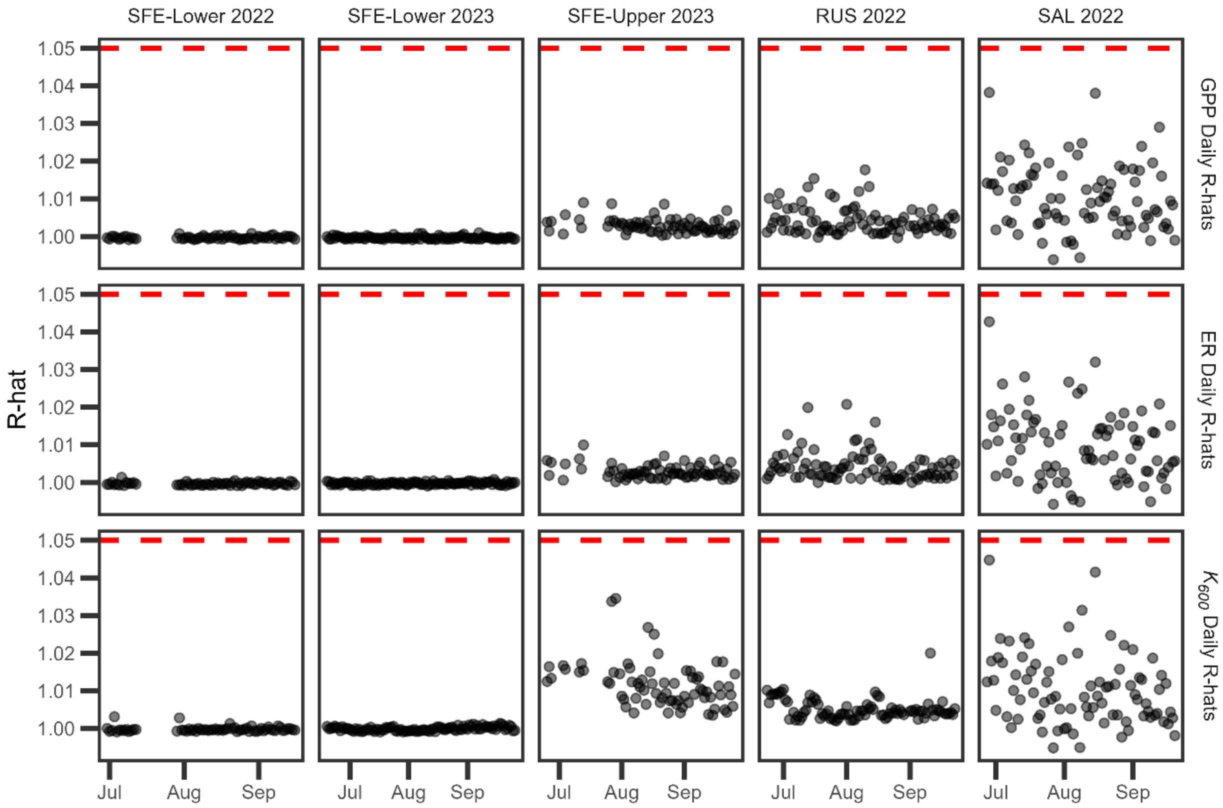


**Figure S7.** Gelman-Rubin diagnostics (r-hats) for daily gross primary productivity (GPP), ecosystem respiration (ER), and the $O_2$ gas exchange rate ($K_{600}$) estimates at all sites below the 1.05 threshold (shown with a dashed red line) to indicate model convergence.

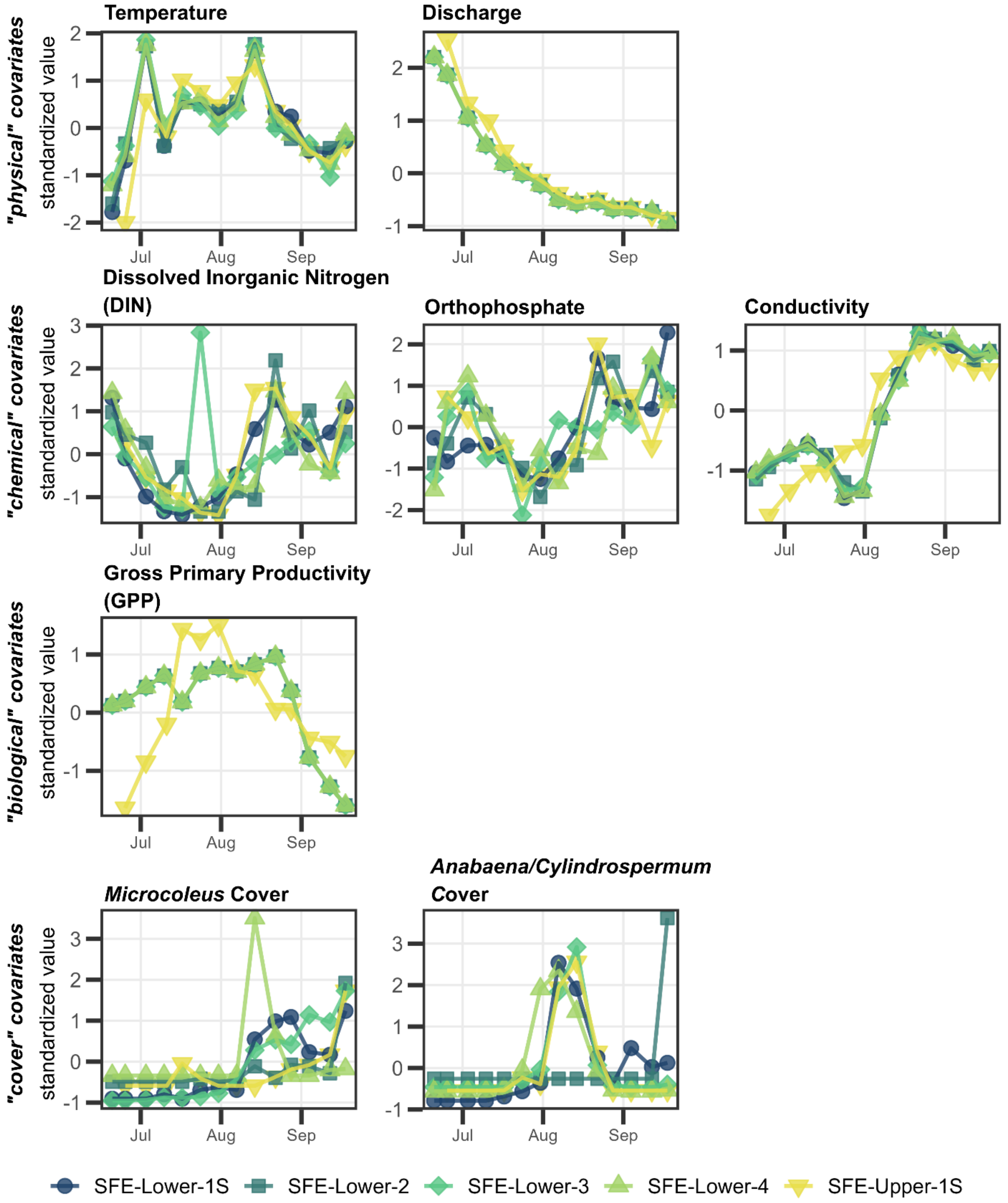


**Figure S8**. Covariates used in predictive modeling (Q3) of the South Fork Eel River (SFE). Each covariate was standardized while grouped by reach.

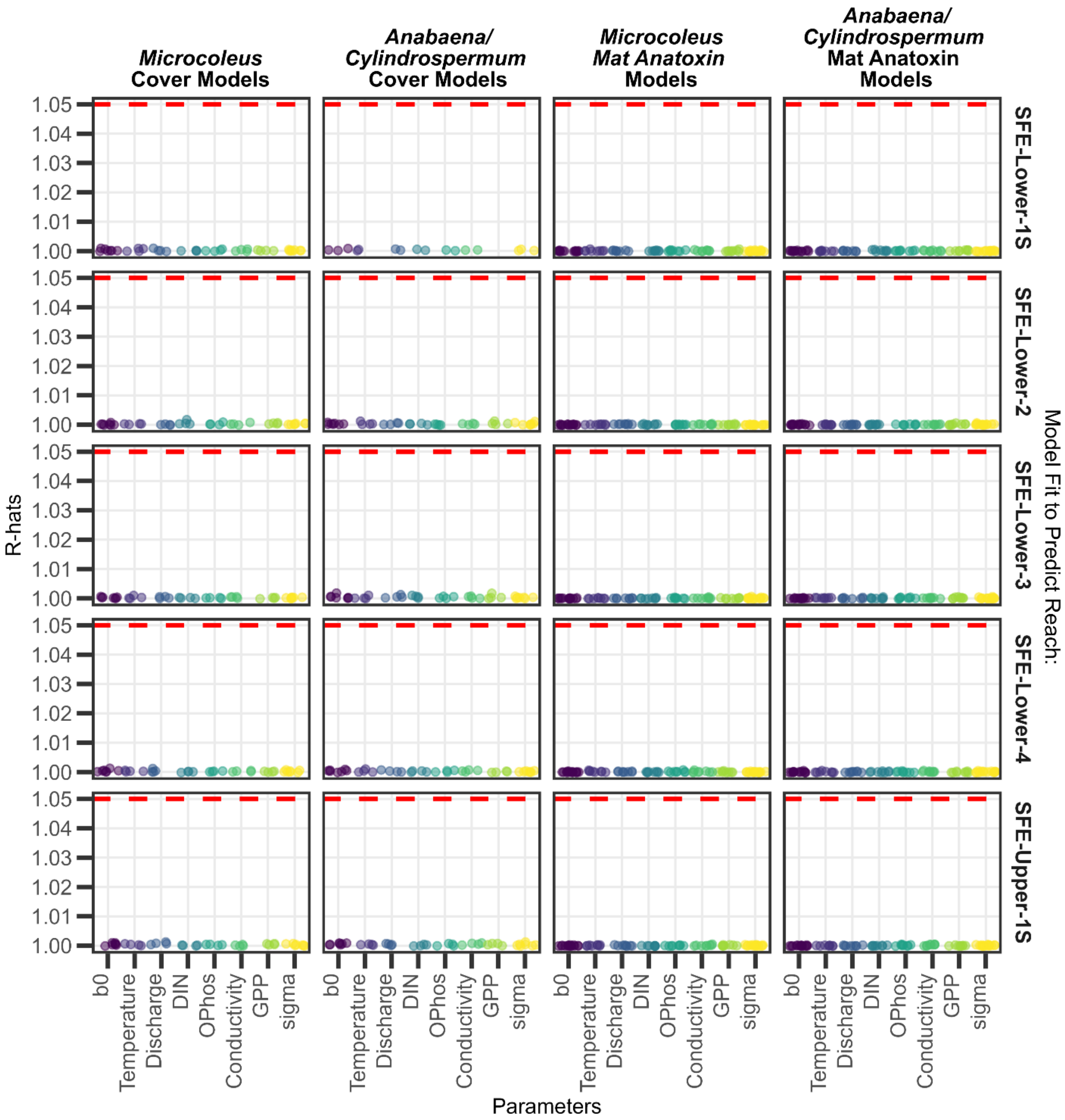


**Figure S9.** Gelman-Rubin diagnostics (r-hats) for parameters in predictive models of the South Fork Eel River (SFE), all below the 1.05 threshold (shown with a dashed red line) to indicate model convergence. Abbreviated parameters on the x-axis are defined as follows: dissolved inorganic nitrogen ("DIN"), orthophosphate ("OPhos"), and gross primary productivity ("GPP").

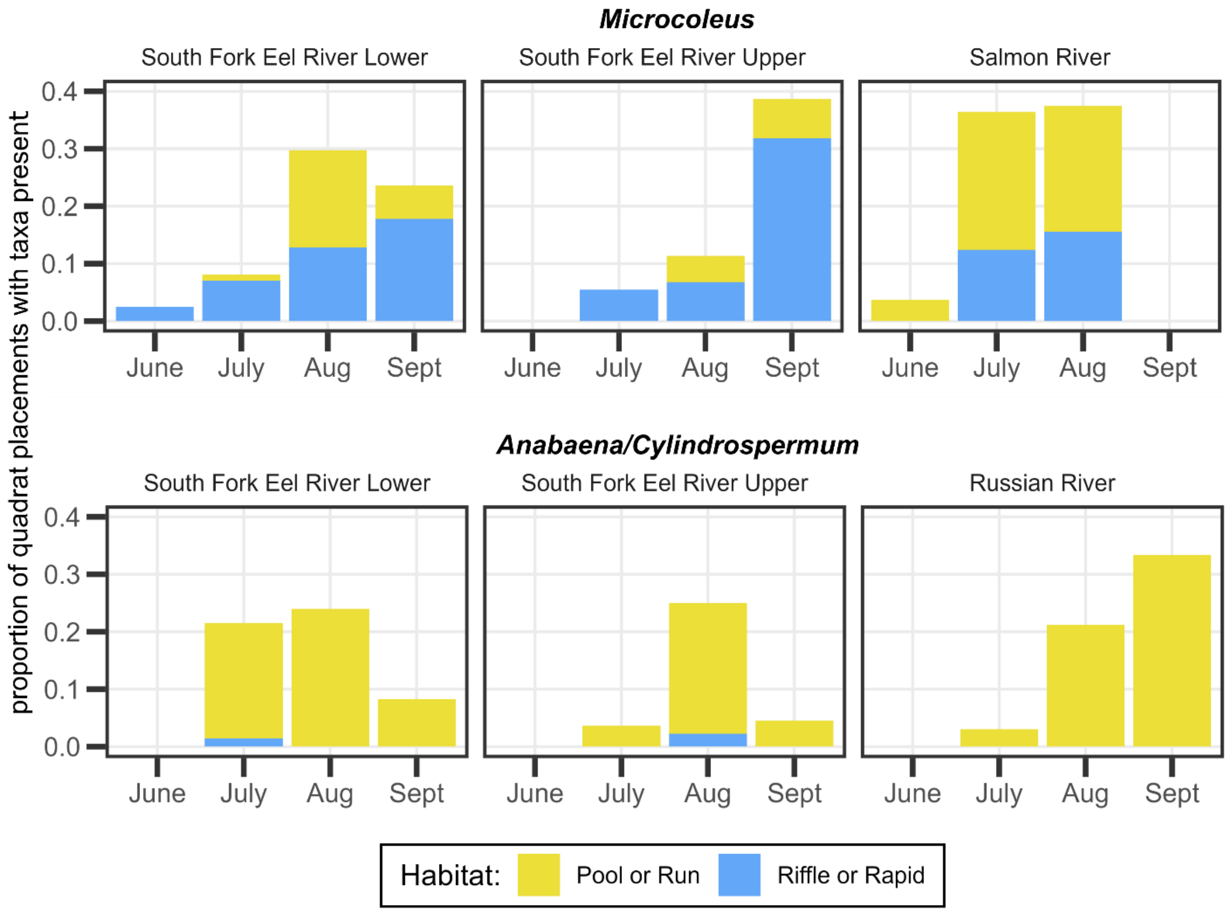


**Figure S10.** Proportion of quadrat placements from percent cover surveys that contained either *Microcoleus* or *Anabaena*/*Cylindrospermum*. Color indicates habitat (i.e., run or pool, or, riffle or rapid) of quadrat placement.

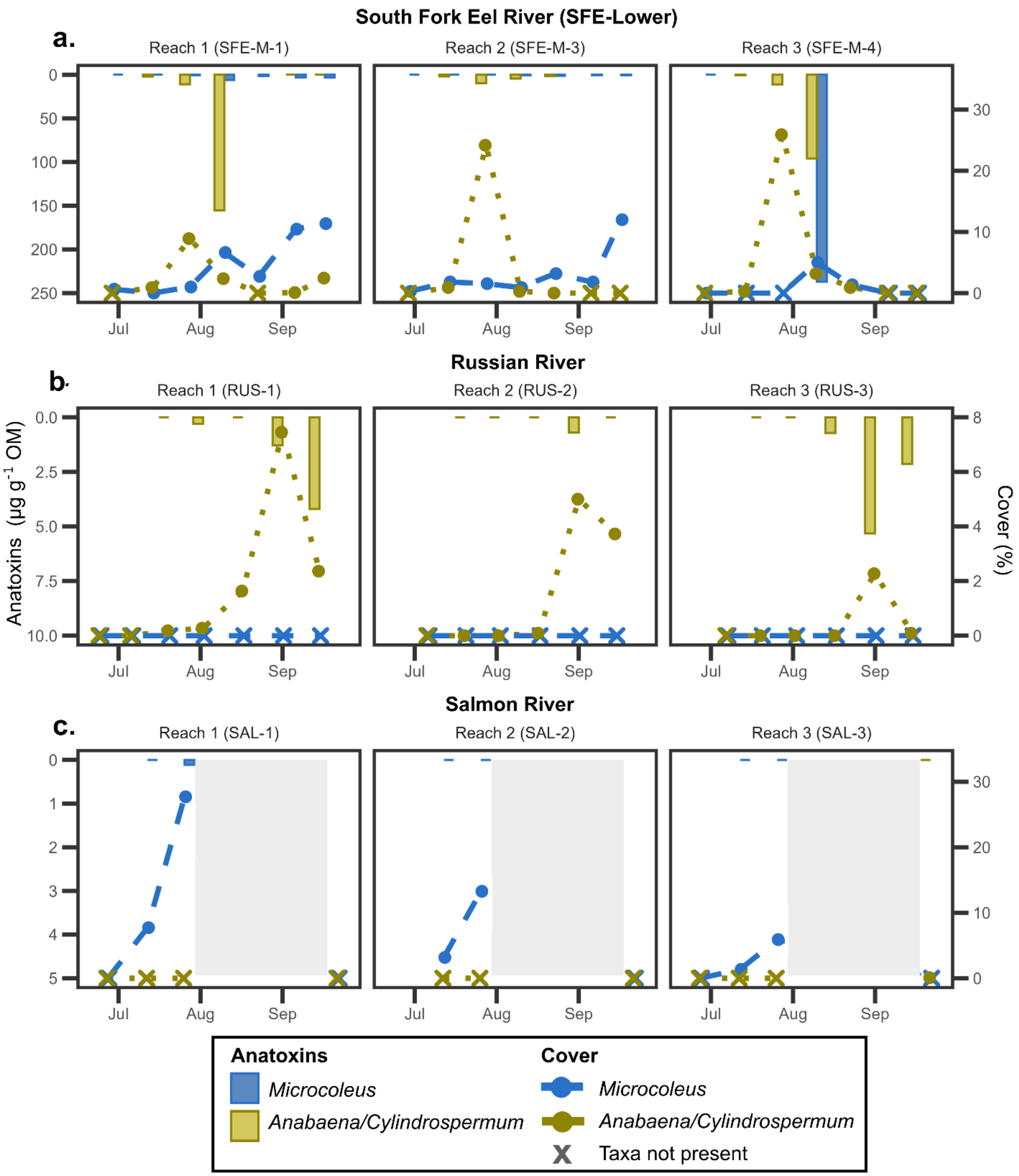


**Figure S11.** Taxon-specific benthic cyanobacteria cover and anatoxin concentrations for each reach sampled on the (a) South Fork Eel, (b) Russian, and (c) Salmon Rivers. Anatoxins are shown as the mean across all three reaches (bars increasing downward where absence of a bar indicates no sample was taken or not enough sample was available for analyses) associated with the left y-axis. Cover is shown as the mean (points and lines increasing upward) across all three reaches with  +/- one standard deviation (error bars) associated with the right y-axis.

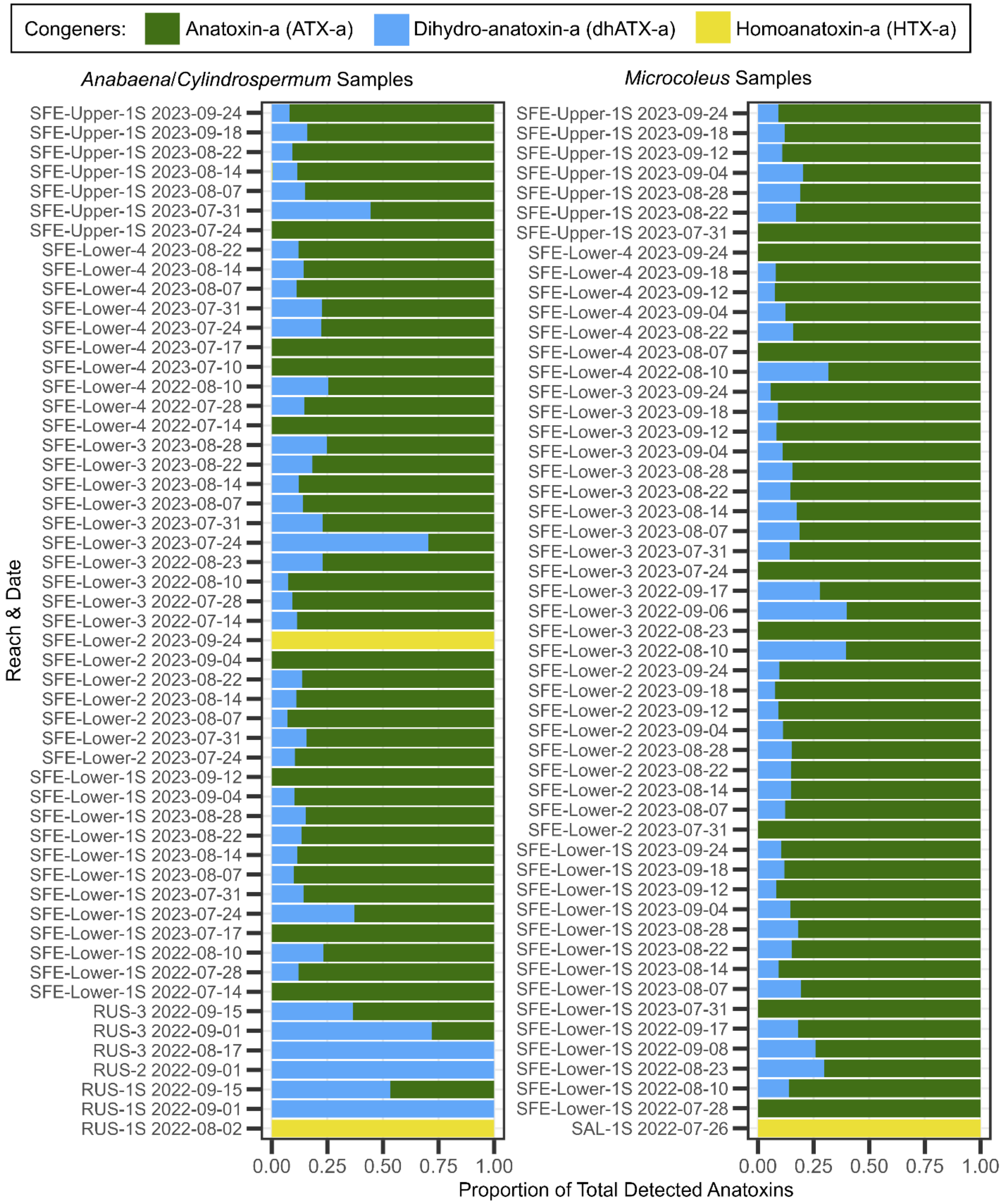


**Figure S12.** Proportion of analyzed anatoxin congeners detected for each benthic cyanobacteria sample. Sample names include the river abbreviation: "SFE" for South Fork Eel River, "SAL" for Salmon River, and "RUS" for Russian River.

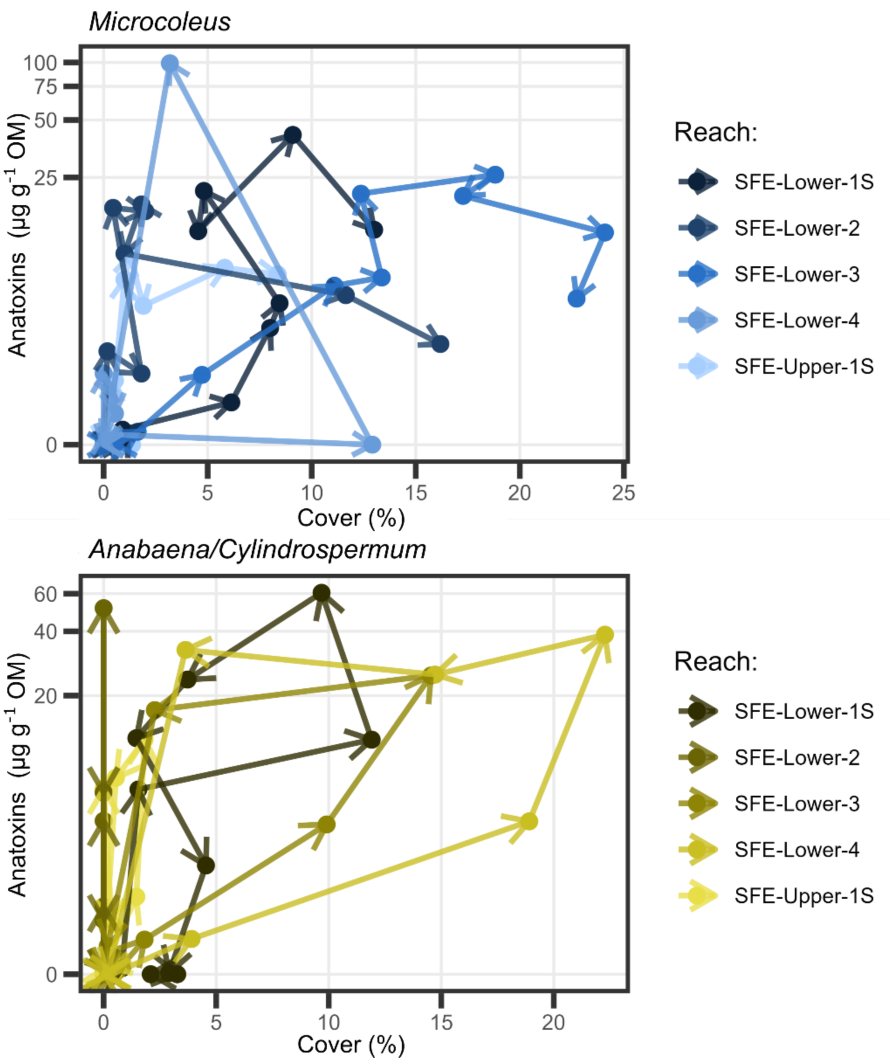


**Figure S13.** Taxon-specific benthic cyanobacteria anatoxin concentrations versus cover for *Microcoleus* and *Anabaena*/*Cylindrospermum* for each reach sampled in the South Fork Eel River (SFE) in 2023. Arrows indicate direction of time across the summer.

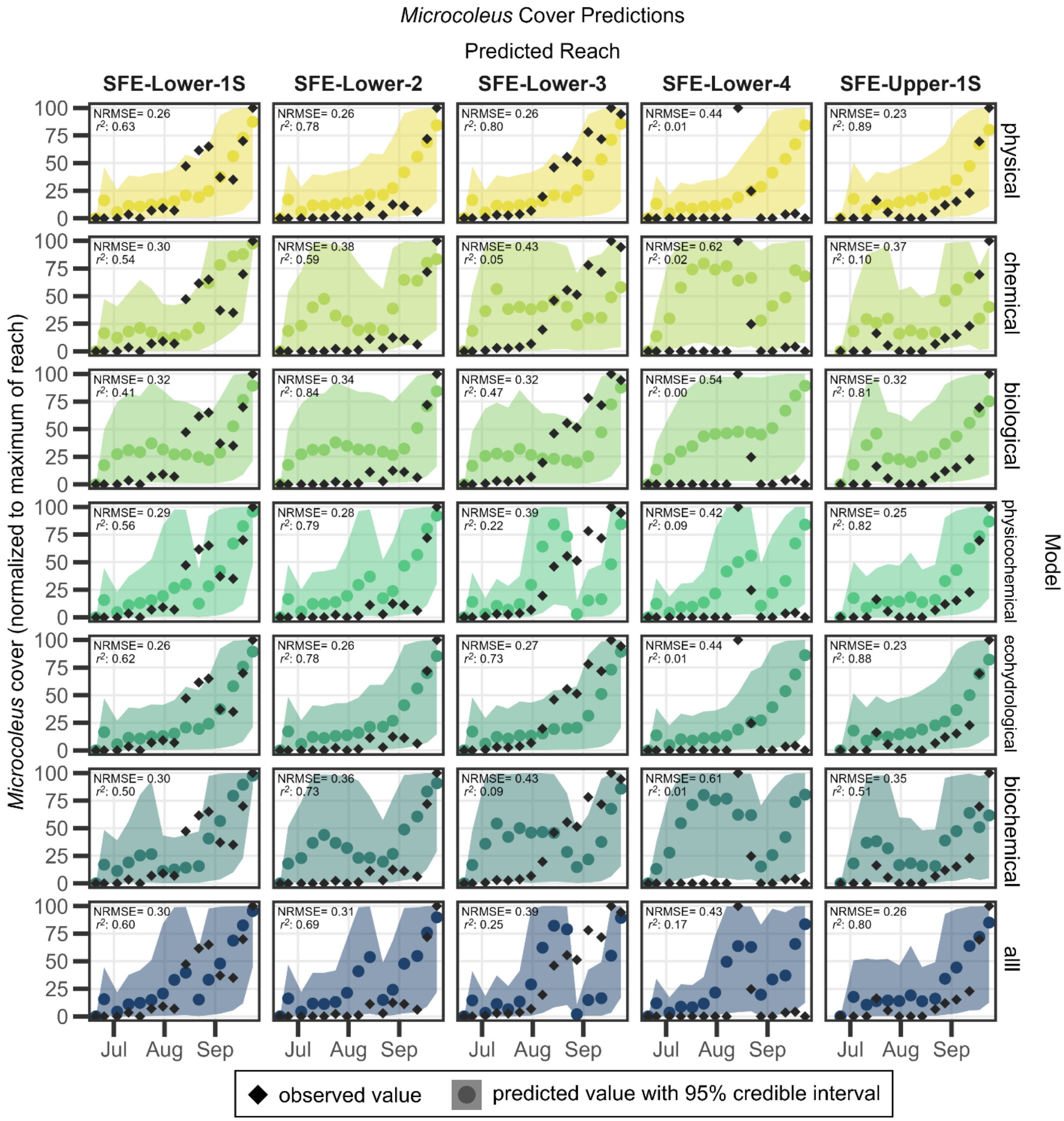


**Figure S14.** Predicted versus observed values for weekly predictions of *Microcoleus* cover in the South Fork Eel River (SFE; normalized to maximum observed at a given reach).

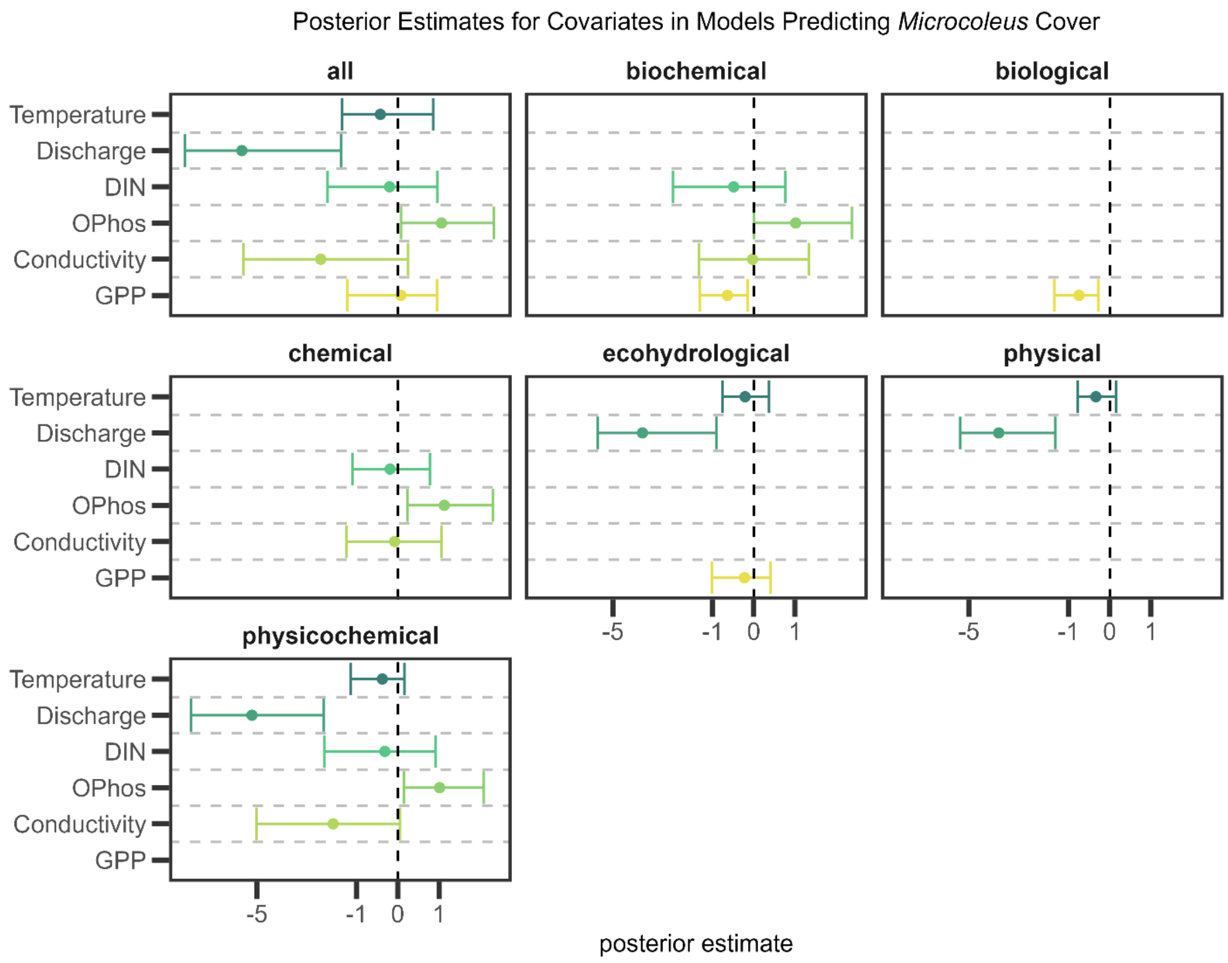


**Figure S15.** Mean (point) and 95% credible interval (range) of posterior estimates for covariates in models predicting *Microcoleus* cover.

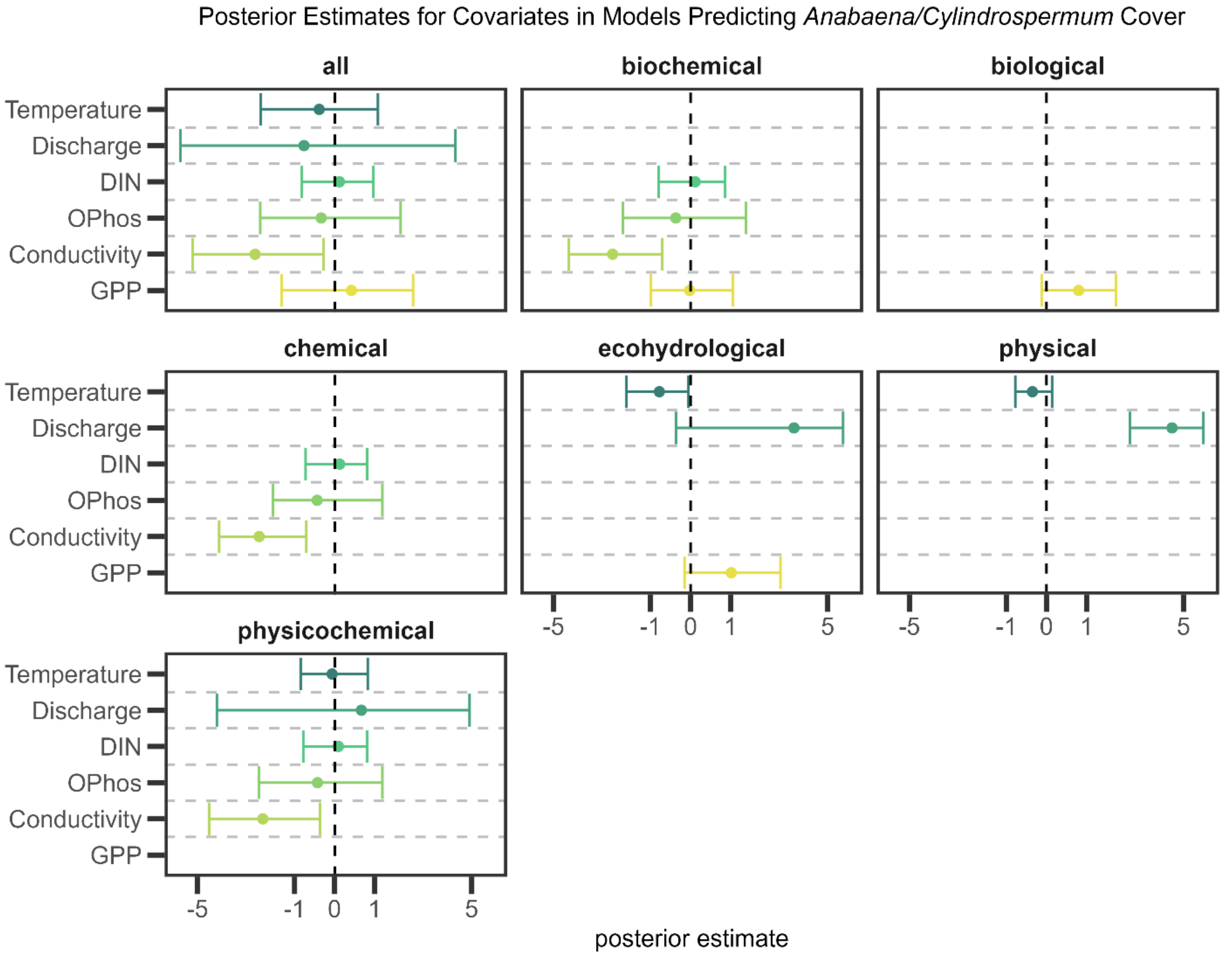


**Figure S16.** Mean (point) and 95% credible interval (range) of posterior estimates for covariates in models predicting *Anabaena/Cylindrospermum* cover. Abbreviated parameters on the y-axis are defined as follows: dissolved inorganic nitrogen ("DIN"), orthophosphate ("OPhos"), and gross primary productivity ("GPP").

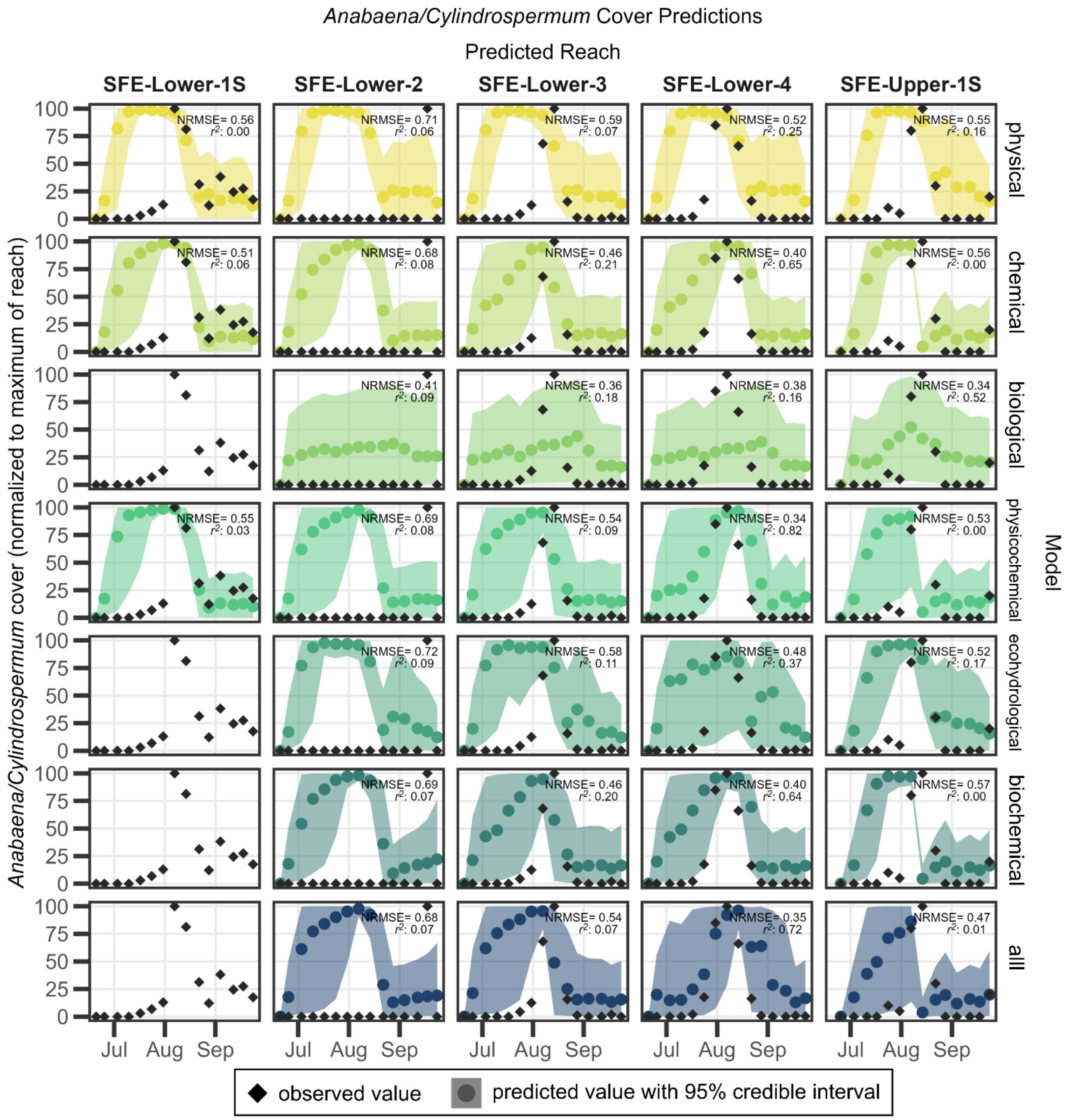


**Figure S17.** Predicted versus observed values for weekly predictions of *Anabaena/Cylindrospermum* cover (normalized to maximum observed at a given reach) in the South Fork Eel River (SFE). The four panels that do not have a colored point and ribbon did not have models that converged so are omitted.

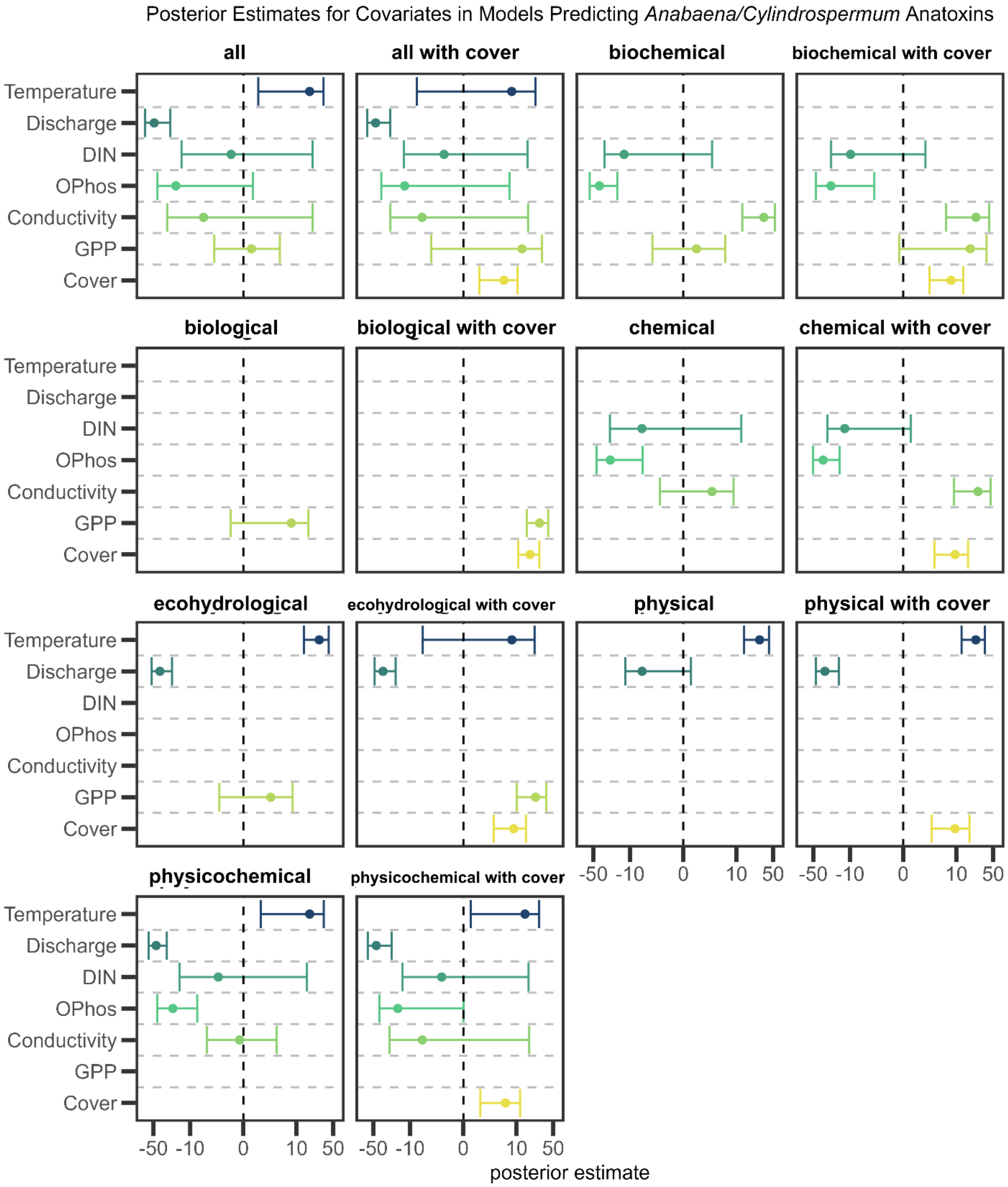


**Figure S18.** Mean (point) and 95% credible interval (range) of posterior estimates for covariates in models predicting *Anabaena/Cylindrospermum* anatoxin concentrations. Abbreviated parameters on the y-axis are defined as follows: dissolved inorganic nitrogen ("DIN"), orthophosphate ("OPhos"), and gross primary productivity ("GPP").

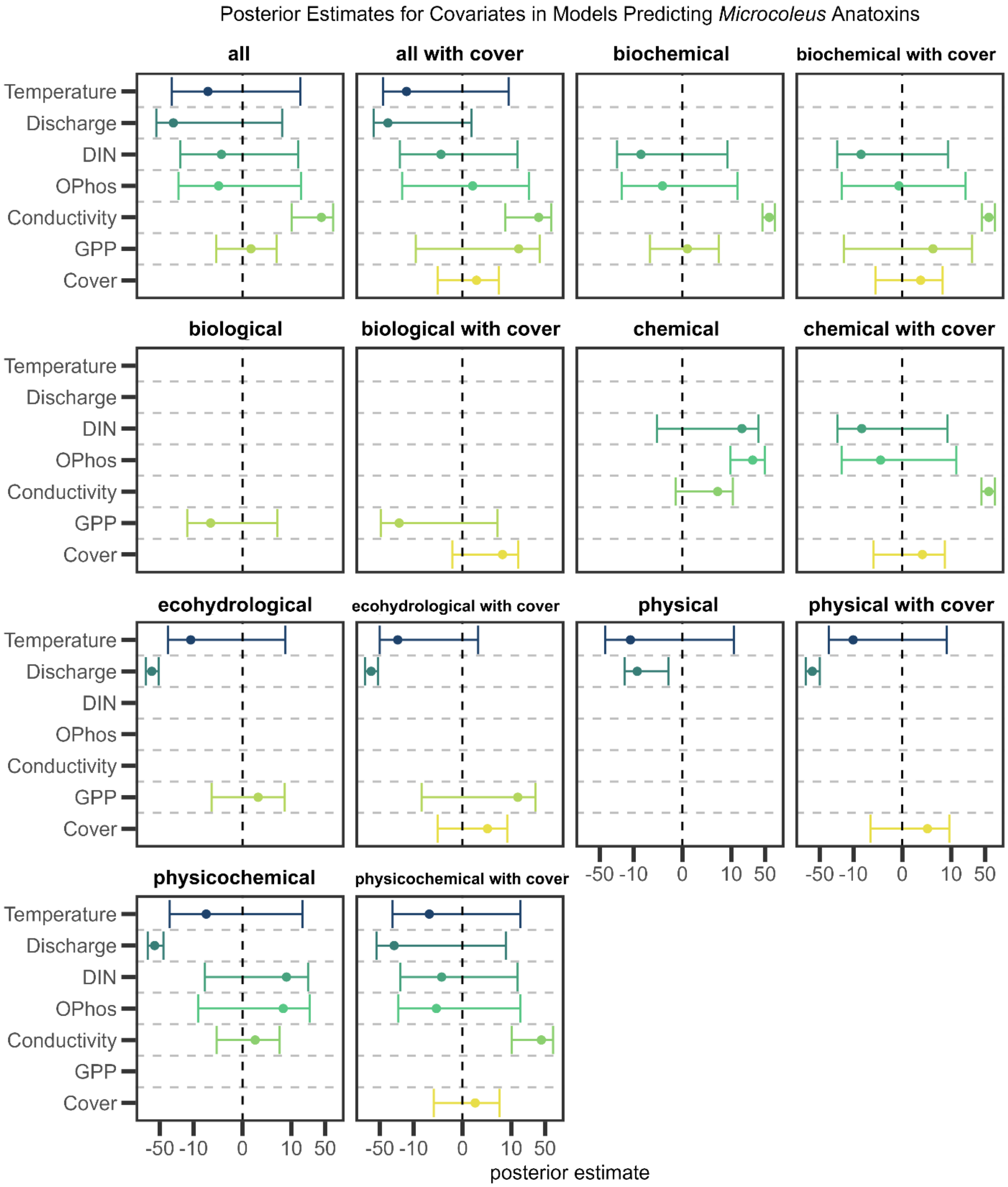


**Figure S19.** Mean (point) and 95% credible interval (range) of posterior estimates for covariates in models predicting *Microcoleus* anatoxin concentrations. Abbreviated parameters on the y-axis are defined as follows: dissolved inorganic nitrogen ("DIN"), orthophosphate ("OPhos"), and gross primary productivity ("GPP").

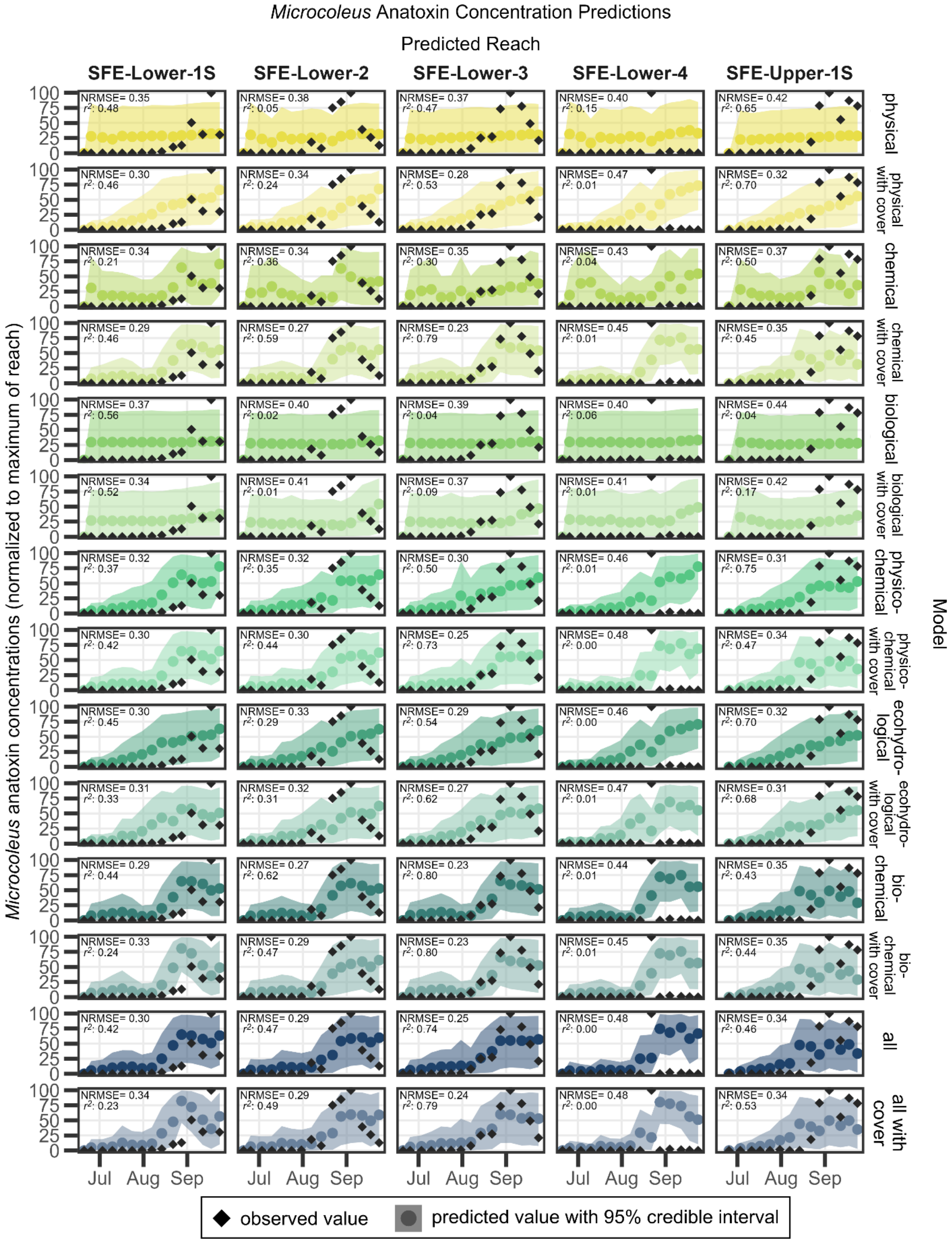


**Figure S20.** Predicted versus observed values for weekly predictions of *Microcoleus* anatoxins (normalized to maximum observed at a given reach) in the South Fork Eel River (SFE).

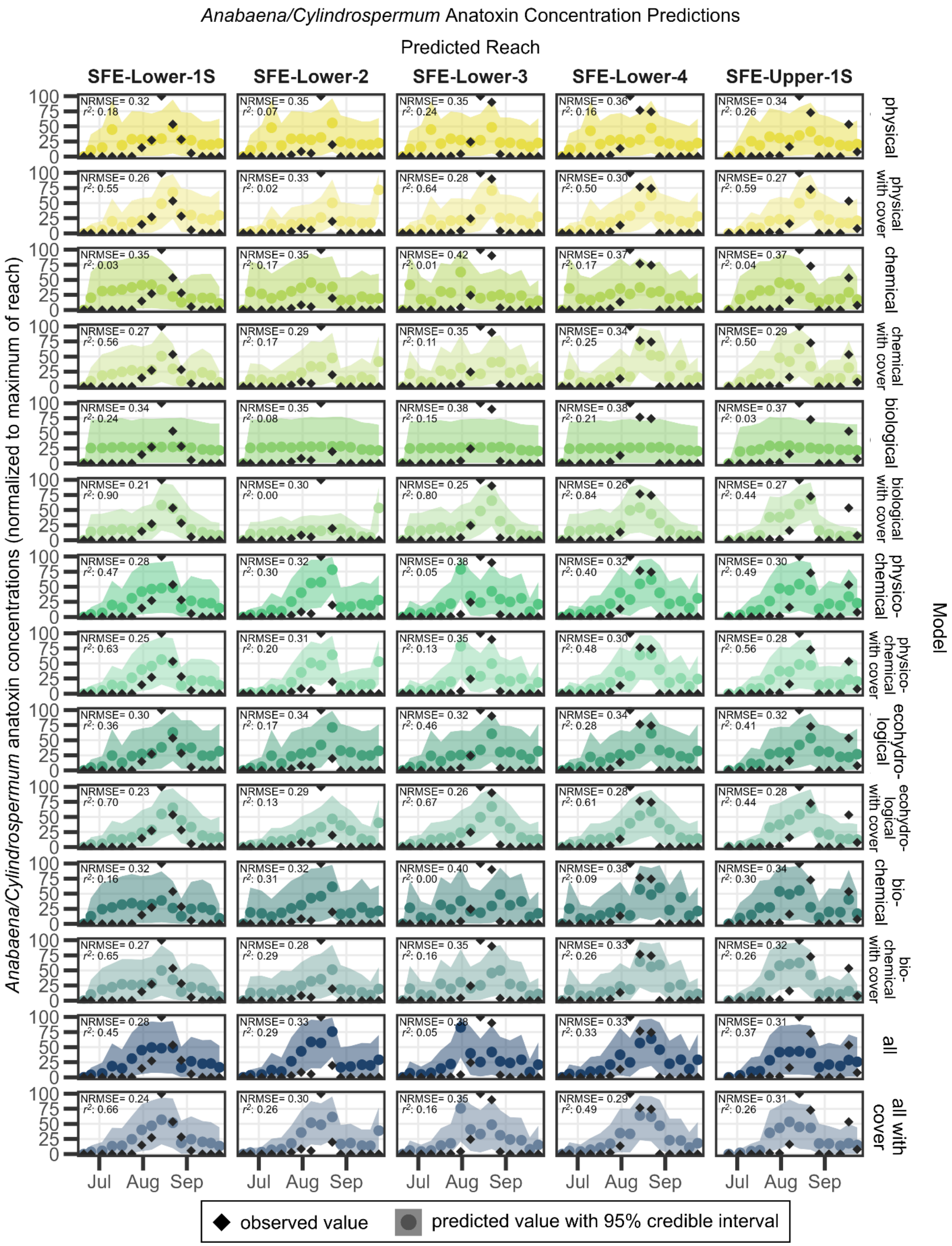


**Figure S21.** Predicted versus observed values for weekly predictions of *Anabaena/Cylindrospermum* anatoxins (normalized to maximum observed at a given reach)

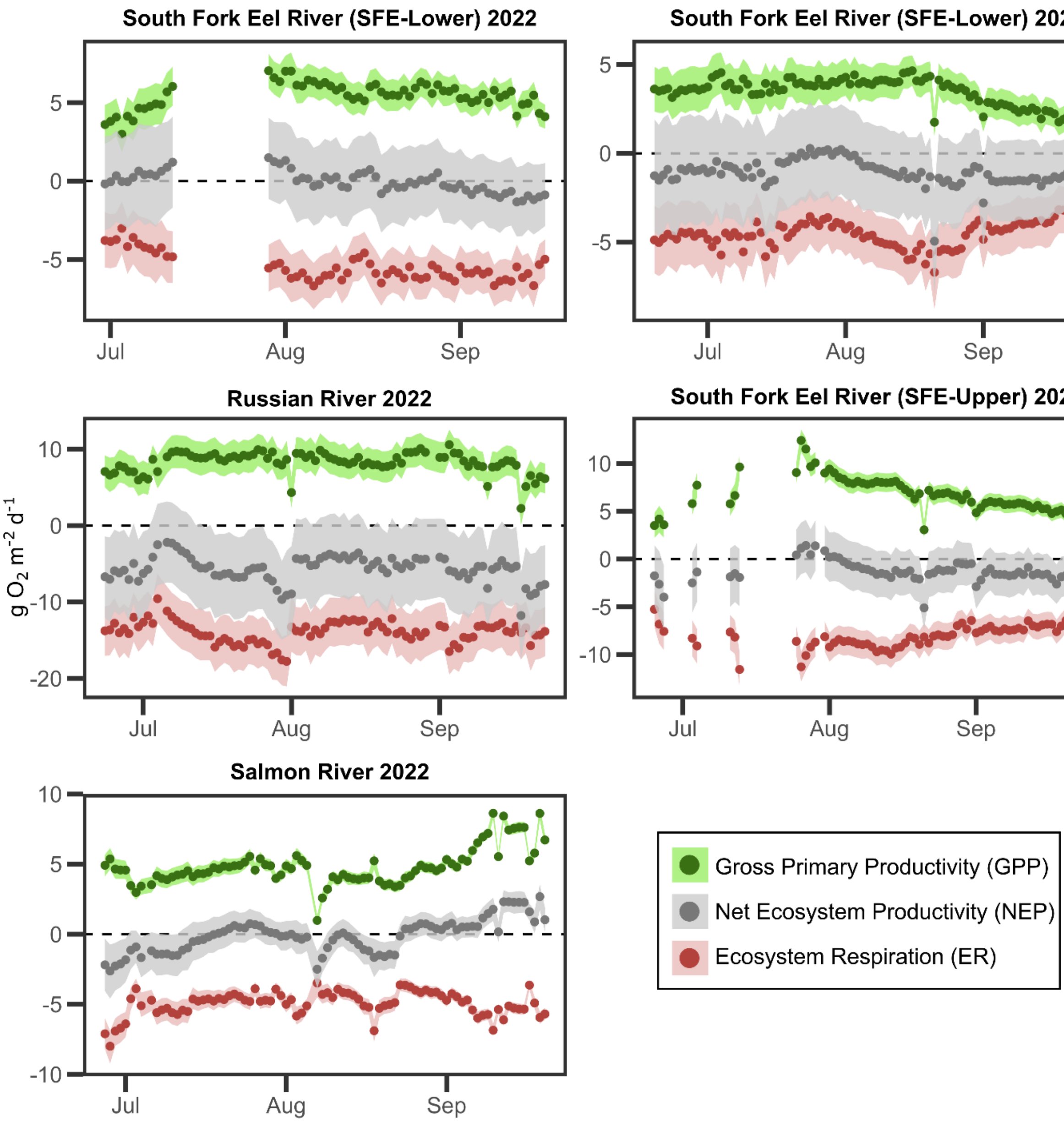


**Figure S22.** Estimates of gross primary productivity (GPP), ecosystem respiration (ER), and net primary productivity (NEP) for all sites sampled each year. NEP was calculated as the difference between estimated GPP and ER. Estimates are shown as the mean (point) with 95% credible interval (ribbon).

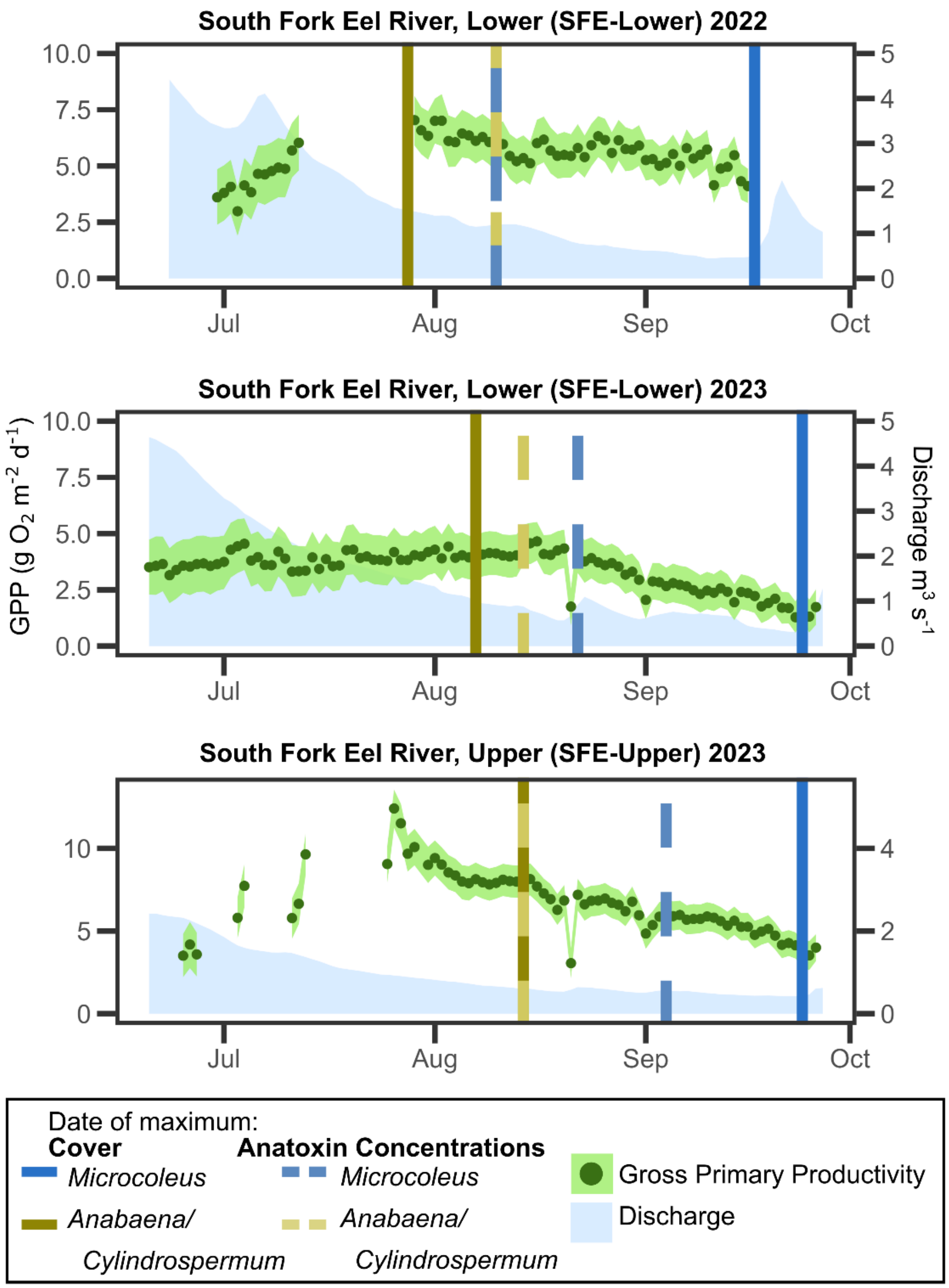


**Figure S23.** Gross primary productivity (GPP) estimates and discharge for the South Fork Eel River Lower in 2022 and 2023 and Upper in 2022. GPP is shown as the mean (point) with 95% credible interval (ribbon) associated with the left y-axis and discharge is shown as the blue area associated with the right y-axis. Vertical lines show dates with maximum taxon-specific cover and anatoxin concentrations (averaged across all four study reaches for SFE-Lower and for the single study reach in SFE-Upper).

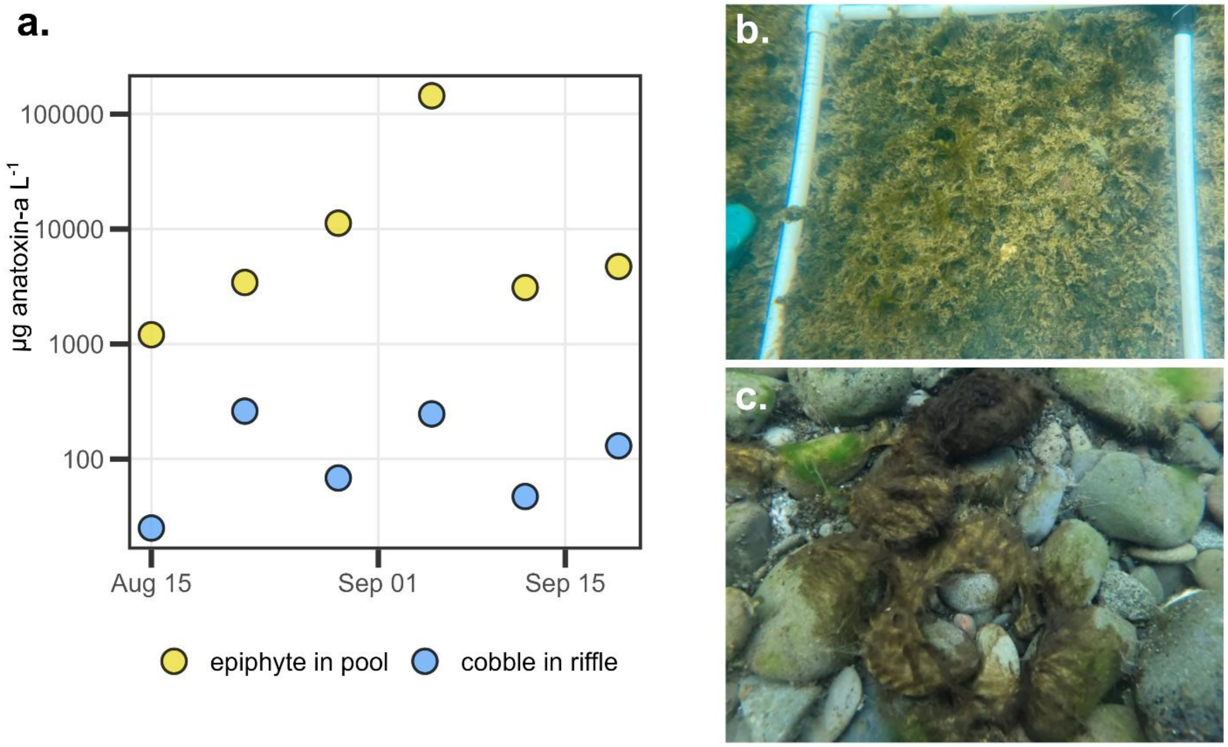


**Figure S24.** (a) Anatoxin-a concentrations for single *Microcoleus* mats in the South Fork Eel River taken either growing (b) as an epiphyte in pools or slower-moving regions or (c) on a cobble in a riffle. Data are from a North Coast Regional Water Quality Control Board study that occurred concurrently with this study during the summer of 2023 at reach SFE-Lower-1S and used with permission (NCRWQCB 2024). Photo credits: Taryn Elliott (b) and Jordan Zabrecky (c).

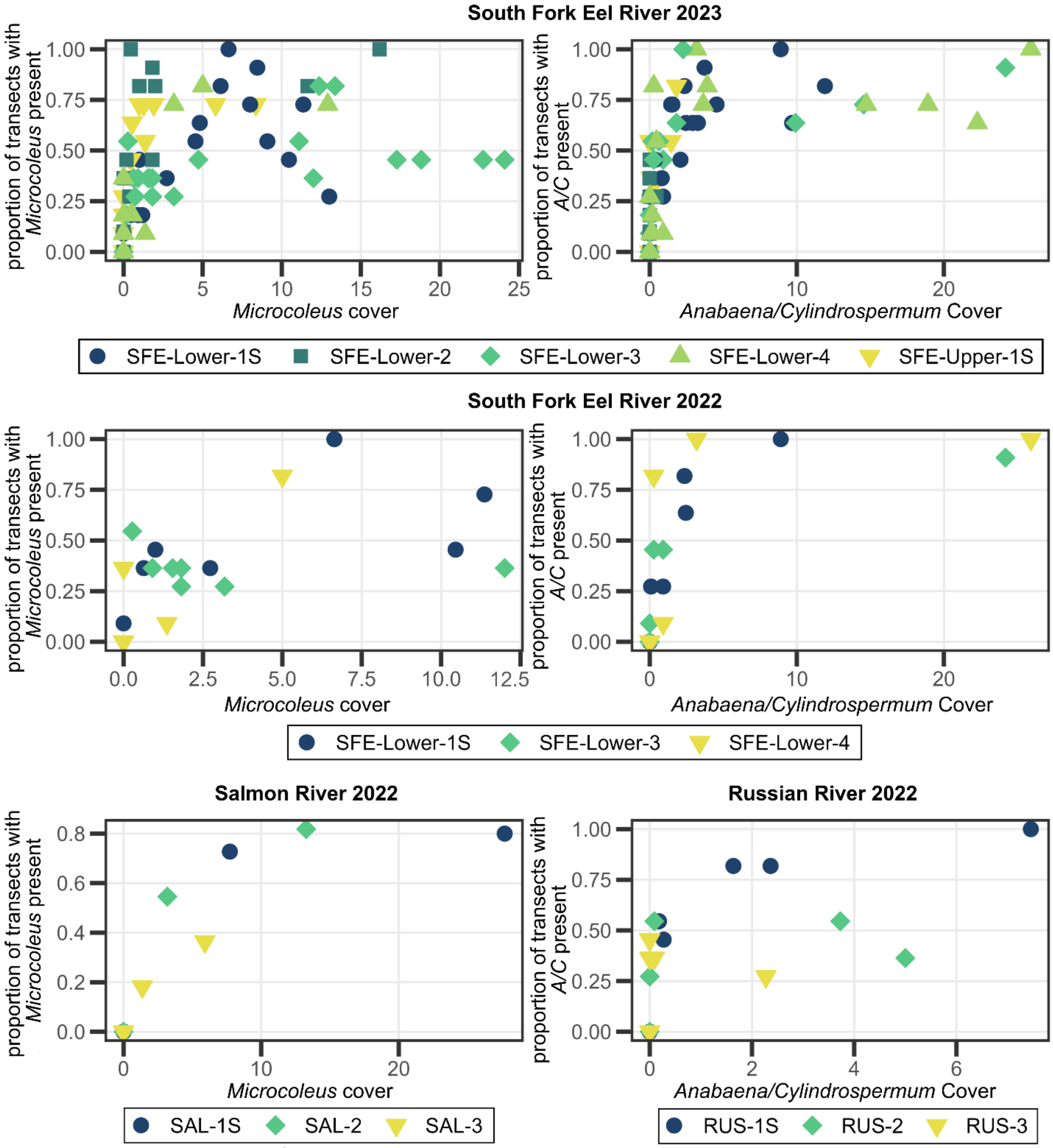


**Figure S25.** Proportion of transects sampled with presence vs. percent cover as obtained by study design for *Microcoleus* or *Anabaena*/*Cylindrospermum* on each river sampled each year. Note that many transects can contain presence of taxa, but percent cover may still be low either due to very small amounts of taxa at each transect or presence was mostly outside of 25, 50, 75% river width intervals used with quadrat percent cover survey.

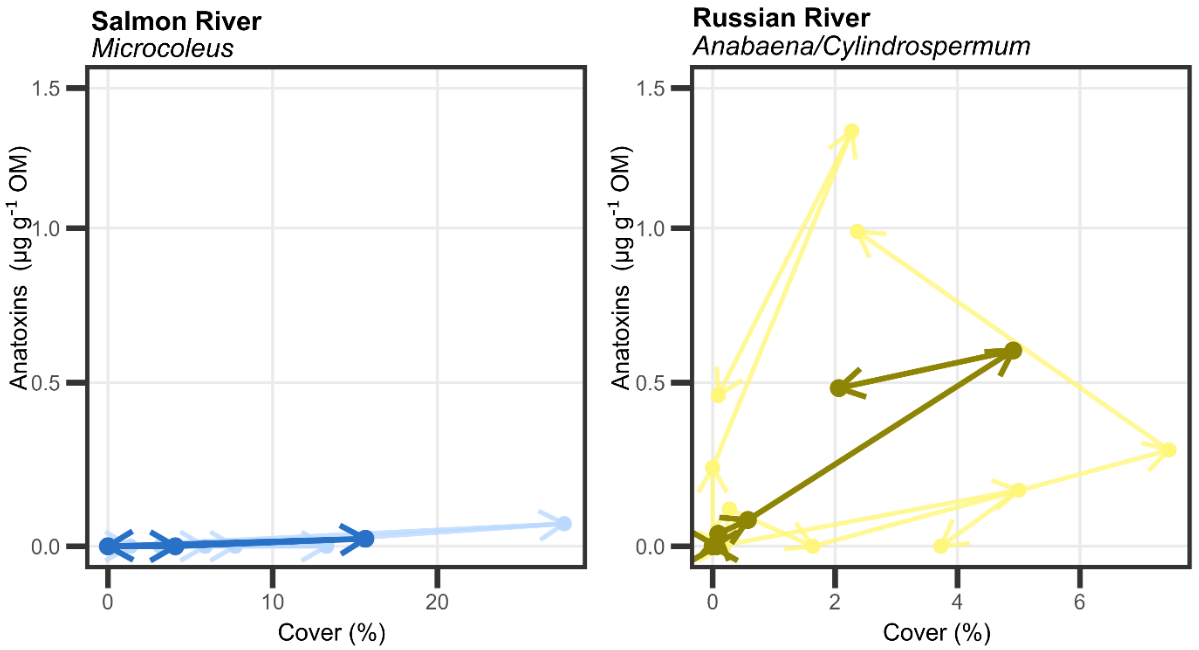


**Figure S26.** Taxon-specific benthic cyanobacteria anatoxin concentrations versus cover for *Microcoleus* in the Salmon River and *Anabaena*/*Cylindrospermum* in the Russian River from 2022 biweekly sampling. Mean behavior across all reaches is shown with the darker bolded line and individual reach behavior are shown with lighter lines. Arrows indicate direction of time across the summer.

**Table S1.** Watershed land cover attributes and range of observed water quality parameters and discharge for each sensor site during the study period. Land cover attributes were obtained from Model My Watershed (Stroud Water Research Center 2017) using U.S. Geological Survey (USGS) Hydrologic Units (HUC-10) and the 2019 National Land Cover Database (Dewiz 2021). Discharge was obtained from the nearby USGS gage (U.S. Geological Survey 2025).Water quality parameters were obtained from discrete sampling events as part of this study (Zabrecky et al. 2025). Values for the Russian River and Salmon River are from 2022, values for the South Fork Eel River (Upper) are from 2023, and values for the South Fork Eel River (Lower) are from both 2022 (top values) and 2023 (bottom values).

| | | **South Fork Eel River (Lower)** ***(SFE-Lower)*** | **South Fork Eel River (Upper)** ***(SFE-Upper)*** | **Russian River** ***(RUS)*** | **Salmon River** ***(SAL)*** |
|---|---|---|---|---|---|
| Corresponding USGS stations | USGS Watershed Unit (HUC-10 No.) | Lower South Fork Eel River (1801010604) | Middle South Fork Eel River (1801010603) | Upper Russian River (1801011004) | Salmon River (1801021004) |
| | USGS Gage (Gage No.) | SF Eel R NR Miranda CA (11476500) | SF Eel R a Leggett CA (11475800) | Russian R NR Cloverdale CA (11463000) | Salmon R a Somes Bar CA (11522500) |
| Land Cover Attributes | % Developed | 5.54% | 4.61% | 7.21% | 2.84% |
| | % Agriculture | 0.02% | 0.00% | 3.43% | 0.06% |
| | % Forested | 77.92% | 90.63% | 34.94% | 56.60% |
| | % Shrub or Grassland | 15.36% | 3.84% | 53.95% | 40.26% |
| Range of Discharge and Water Quality Parameters | Discharge ($m^3 s^{-1}$) | 0.4-4.2<br>0.3-4.8 | 0.4-2.4 | 0.8-2.1 | 4.4-19.1 |
| | Temperature (°C) | 19.7-28.5<br>19.0-29.2 | 17.2-22.7 | 18.8-24.8 | 16.6-22.6 |
| | pH | 7.97-8.68<br>7.90-8.04 | 7.84-8.04 | 7.98-8.56 | 7.72-8.35 |
| | Specific Conductivity ($\mu S\ cm^{-1}$) | 216.4-251.2<br>206.0-262.0 | 185.9-240.0 | 235.3-267.8 | 108.5-125.5 |
| | Ammonium ($mg\ N\ L^{-1}$) | 0.003-0.063<br><0.002-0.019 | <0.002-0.010 | 0.003-0.031 | 0.007-0.015 |
| | Nitrate ($mg\ N\ L^{-1}$) | 0.005-0.033<br>0.009-0.056 | 0.008-0.038 | 0.007-0.020 | 0.008-0.031 |
| | Ortho-phosphate ($\mu g\ P\ L^{-1}$) | 3.50-7.01<br>3.24-6.95 | 3.66-5.54 | 5.03-11.20 | 4.99-8.04 |

**Table S2.** Location and sampling duration and frequency of reaches sampled in 2022. Reaches with a suffix of "S" are the most-downstream reaches that have the dissolved oxygen placed either within-reach or downstream (within 250 m). Latitude and longitude are the coordinates for the most downstream transect of the reach.

| River | Reach | Latitude | Longitude | Sampling Duration & Frequency |
|---|---|---|---|---|
| South Fork Eel River (Lower) (SFE-Lower) | SFE-Lower-1S | 40.199481 | -123.775892 | 6/29/2022 - 9/17/2022 (7 events; biweekly) |
| | SFE-Lower-3 | 40.188030 | -123.771416 | 6/29/2022 - 9/17/2022 (7 events; biweekly) |
| | SFE-Lower-4 | 40.182204 | -123.776178 | 6/29/2022 - 9/17/2022 (7 events; biweekly) |
| Russian River (RUS) | RUS-1S | 38.806280 | -123.006417 | 6/24/2022 - 9/15/2022 (7 events; biweekly) |
| | RUS-2 | 38.832283 | -123.011850 | 7/6/2022 - 9/15/2022 (6 events; biweekly) |
| | RUS-3 | 38.843114 | -123.023883 | 7/7/2022 - 9/15/2022 (6 events; biweekly) |
| Salmon River (SAL) | SAL-1S | 41.378694 | -123.475306 | 6/27/2022 - 9/22/2022 (4 events[1]; biweekly) |
| | SAL-2 | 41.378127 | -123.463236 | 7/12/2022 - 9/22/2022 (3 events[1]; biweekly) |
| | SAL-3 | 41.373662 | -123.454126 | 6/27/2022 - 9/22/2022 (4 events[1]; biweekly) |

[1]*Sampling on the Salmon River was interrupted by wildfires from 8/9/2022 to 9/6/2022*

**Table S3.** Median velocity and $O_2$ gas exchange rate ($K_{600}$) used with the equation $3v/K$ to estimate the 95% dissolved oxygen residence distance and the river distance of the most upstream 150-m reach from the dissolved oxygen sensor.

| **Sensor Site** | **Median Velocity** (m/s) | **Median Estimated Gas Exchange, $K_{600}$** ($d^{-1}$) | **Estimated Dissolved Oxygen Footprint** (km) | **Distance from Sensor to Furthest Upstream 150 m Survey Reach** (km) |
|---|---|---|---|---|
| South Fork Eel River (Lower) (SFE-Lower) | 0.20 | 3.4 | 15.2 | 2.5 |
| South Fork Eel River (Upper) (SFE-Upper) | 0.22 | 4.7 | 11.9 | 0.4 |
| Salmon River (SAL) | 0.38 | 10.2 | 9.6 | 2.7 |
| Russian River (RUS) | 0.41 | 5.2 | 20.6 | 5.3 |

**Table S4.** Location and sampling duration and frequency of reaches sampled in 2023. Reaches with a suffix of "S" are the most-downstream reaches that have the dissolved oxygen placed either within-reach or downstream (within 250 m). Latitude and longitude are the coordinates for the most downstream transect of the reach.

| Sensor Site | Reach | Latitude | Longitude | Sampling Duration & Frequency |
|---|---|---|---|---|
| South Fork Eel River (Lower) (SFE-Lower) | SFE-Lower-1S | 40.199481 | -123.775892 | 6/20/2023 - 9/24/2023 (15 events; weekly) |
| | SFE-Lower-2 | 40.192318 | -123.769671 | 6/20/2023 - 9/24/2023 (15 events; weekly) |
| | SFE-Lower-3 | 40.188030 | -123.771416 | 6/20/2023 - 9/24/2023 (15 events; weekly) |
| | SFE-Lower-4 | 40.182204 | -123.776178 | 6/20/2023 - 9/24/2023 (15 events; weekly) |
| South Fork Eel River (Upper) (SFE-Upper) | SFE-Upper-1S | 39.876046 | -123.726920 | 6/25/2023 - 9/24/2023 (14 events; weekly) |

**Table S5.** Microcystin concentrations for samples with microcystins detected. Reach names include the river abbreviation: "SFE" for South Fork Eel River, "SAL" for Salmon River, and "RUS" for Russian River.

| Composite Sample Type | Reach Collected | Date Sampled | Microcystins (µg microcystins $g^{-1}$ lyophilized sample) |
|---|---|---|---|
| *Microcoleus* | SAL-2 | 2022-07-12 | 1.37 |
| *Microcoleus* | SFE-Lower-1S | 2022-07-14 | 0.73 |
| *Microcoleus* | SAL-1S | 2022-07-26 | 1.11, 1.08* |
| *Anabaena/Cylindrospermum* | SFE-Lower-3 | 2023-07-24 | 0.16 |
| *Anabaena/Cylindrospermum* | SFE-Lower-1S | 2023-07-31 | 0.66, 0.64* |
| *Anabaena/Cylindrospermum* | SFE-Lower-3 | 2023-07-31 | 0.35 |
| *Anabaena/Cylindrospermum* | SFE-Lower-2 | 2023-09-18 | 0.2, 0.21* |

**indicates composite was split into triplicate samples for toxin analyses (so if two samples are shown, the third did not contain any detectable microcystins)*

**Table S6.** Slope, median depth, median velocity used to calculate $O_2$ gas exchange rate ($K_{600}$) priors using equation 1 from Raymond et al. (2012).

| **Sensor Site** | **Slope** | **Median Depth** (m) | **Median Velocity** (m $s^{-1}$) | $K_{600}$ **Prior Estimate** |
|---|---|---|---|---|
| South Fork Eel River (Lower) (SFE-Lower) | 0.00125 | 0.63 | 0.20 | 2.41 |
| South Fork Eel River (Upper) (SFE-Upper) | 0.00250 | 0.70 | 0.22 | 5.19 |
| Salmon River (SAL) | 0.00460 | 1.11 | 0.38 | 18.6 |
| Russian River (RUS) | 0.00102 | 0.86 | 0.41 | 4.56 |

**Table S7.** Summary statistics for *Microcoleus* and *Anabaena*/*Cylindrospermum* presence/absence, cover, and anatoxin concentrations, and gross primary productivity (GPP) for each river sampled biweekly in 2022. Mean and maximum cover and anatoxin concentrations are whole-river values that were averaged across all three reaches sampled. “NA” indicates “not applicable”.

| River | **South Fork Eel River (SFE-Lower)** | | **Russian River (RUS)** | | **Salmon River[1] (SAL)** | |
|---|---|---|---|---|---|---|
| Taxa | *Microcoleus* | *Anabaena/ Cylindrospermum* | *Microcoleus* | *Anabaena/ Cylindrospermum* | *Microcoleus* | *Anabaena/ Cylindrospermum* |
| Transects Sampled with (Nearby) Presence (%) | 34.6% *(80 of 231)* | 37.2% *(86 of 231)* | 0% *(0 of 209)* | 39.7% *(83 of 209)* | 30.8% *(37 of 120)* | 3.3% *(4 of 120)* |
| Duration of Presence | 6/29-9/17 | 7/14-9/17 | *NA* | 7/20-9/15 | 7/12-7/26 | 9/22 |
| Mean Cover (%) | 2.9% | 3.4% | 0% | 1.2% | 5.4% | 0.008% |
| Max Cover (%) | 12.0% | 25.9% | 0% | 7.5% | 27.7% | 0.09% |
| Date of Max Cover | 9/17 | 7/28 | *NA* | 9/1 | 7/26 | 9/22 |
| Samples with Detected Anatoxins (%) | 62.5% *(10 of 16)* | 83.3% *(10 of 12)* | *NA* | 46.7% *(7 of 15)* | 16.7% *(1 of 6)* | 0% *(0 of 1)* |
| Mean Anatoxin Concentration (μg anatoxins $g^{-1}$ OM) | 12.1 | 14.0 | *NA* | 0.70 | 0.009 | 0.0 |
| Max Anatoxin Concentration (μg anatoxins $g^{-1}$ OM) | 81.3 | 85.2 | *NA* | 2.4 | 0.04 | *NA* |
| Date of Max Anatoxin Concentration | 8/10 | 8/10 | *NA* | 9/1 | 7/26 | *NA* |
| Mean GPP (g $O_2$ $m^{-2}$ $d^{-1}$) | 5.4 | | 8.3 | | 4.8 | |
| Max GPP (g $O_2$ $m^{-2}$ $d^{-1}$) | 7.0 | | 10.5 | | 8.6 | |
| Date of Max GPP | 7/29 | | 9/3 | | 9/10 | |
| Min GPP (g $O_2$ $m^{-2}$ $d^{-1}$) | 3.0 | | 2.2 | | 0.99 | |
| Date of Min GPP | 7/3 | | 9/18 | | 8/7 | |

[1]*Sampling on the Salmon River was interrupted by wildfires from 8/9/2022 to 9/6/2022 reducing the total sampling events from seven to four*

**Table S8.** Anatoxin concentrations for *Microcoleus* and *Anabaena*/*Cylindrospermum* samples for all three reaches on each river sampled biweekly in 2022. Numeric values are total anatoxin concentrations in µg anatoxins per g organic matter; “not present” indicates that no samples were taken as the taxa was not visibly present; “limited sample” indicates that not enough of the taxa was present to collect and analyze for anatoxin; “no river visit” indicates that we did not sample that river on that date; “no reach visit” indicates that we did not sample that reach on that date; and “ND” indicates that a sample was taken but no anatoxins were detected. Values are comma separated in order of most downstream to most upstream reach (SFE-Lower-1S, SFE-Lower-3, SFE-Lower-4; RUS-1S, RUS-2, RUS-3; SAL-1S, SAL-2, SAL-3).

| River | **South Fork Eel River**<br>***(SFE-Lower)*** | | **Russian River**<br>***(RUS)*** | **Salmon River**<br>***(SAL)*** | |
|---|---|---|---|---|---|
| Taxa | *Microcoleus* | *Anabaena/*<br>*Cylindrospermum* | *Anabaena/*<br>*Cylindrospermum* | *Microcoleus* | *Anabaena/*<br>*Cylindrospermum* |
| Visit 1<br>*(6/24-6/29)* | ND,<br>ND,<br>ND | not present,<br>not present,<br>not present | not present,<br>no reach visit,<br>no reach visit | not present,<br>no reach visit,<br>not present | not present,<br>no reach visit,<br>not present |
| Visit 2<br>*(7/6-7/14)* | ND,<br>ND,<br>not present | 2.04,<br>1.93,<br>0.62 | not present,<br>not present,<br>not present | ND,<br>ND,<br>ND | not present,<br>not present,<br>not present |
| Visit 3<br>*(7/20-7/28)* | 0.64,<br>ND,<br>not present | 11.0,<br>9.63,<br>11.1 | ND,<br>ND,<br>ND | 0.11,<br>ND,<br>ND | not present,<br>not present,<br>not present |
| Visit 4<br>*(8/2-8/10)* | 6.08,<br>1.80,<br>237.0 | 155.2,<br>4.41,<br>95.9 | 0.30,<br>ND,<br>ND | no river visit[1] | no river visit[1] |
| Visit 5<br>*(8/17-8/23)* | 1.49,<br>1.20,<br>limited sample | not present<br>1.70,<br>limited sample | ND,<br>ND,<br>0.72 | no river visit[1] | no river visit[1] |
| Visit 6<br>*(9/1-9/6)* | 3.20,<br>0.52,<br>not present | ND<br>not present<br>not present | 1.29,<br>0.70,<br>5.32 | no river visit[1] | no river visit[1] |
| Visit 7<br>*(9/15-9/22)* | 3.28,<br>0.63,<br>not present | ND<br>not present<br>not present | 4.20,<br>ND,<br>2.13 | not present,<br>not present,<br>not present | not present,<br>not present,<br>ND |

[1]*Sampling on the Salmon River was interrupted by wildfires from 8/9/2022 to 9/6/2022 reducing the total sampling events from seven to four*

**Table S9.** Summary statistics for *Microcoleus* and *Anabaena*/*Cylindrospermum* presence/absence, cover, and anatoxin concentrations for each reach of the South Fork Eel River (SFE) sampled weekly in 2023.

| Reach | **SFE-Lower-1S** | **SFE-Lower-2** | **SFE-Lower-3** | **SFE-Lower-4** | **SFE-Upper-1S** | **SFE-Lower-1S** | **SFE-Lower-2** | **SFE-Lower-3** | **SFE-Lower-4** | **SFE-Upper-1S** |
|---|---|---|---|---|---|---|---|---|---|---|
| Taxa | *Microcoleus* | | | | | *Anabaena/Cylindrospermum* | | | | |
| Transects Sampled with (Nearby) Presence (%) | 35.8% *(59 of 165)* | 48.8% *(78 of 160)* | 40.0% *(66 of 165)* | 18.8% *(31 of 165)* | 42.9% *(66 of 154)* | 49.1% *(81 of 165)* | 20.0% *(32 of 160)* | 33.9% *(56 of 160)* | 32.1% *(53 of 165)* | 24.0% *(37 of 154)* |
| Duration of Presence | 7/3-9/24 | 6/20; 7/10; 7/24-9/24 | 7/3-9/24 | 6/25; 7/24-9/24 | 6/25-7/31; 8/14-9/24 | 7/10-9/24 | 7/10-9/4; 9/18-9/24 | 7/17-9/4; 9/18-9/24 | 7/10-8/28; 9/18-9/24 | 7/24-8/22; 9/18-9/24 |
| Mean Cover (%) | 3.8% | 2.4% | 8.6% | 1.1% | 1.5% | 2.8% | 0.03% | 2.0% | 4.3% | 0.3% |
| Max Cover (%) | 13.0% | 16.2% | 24.1% | 12.9% | 8.4% | 11.9% | 0.5% | 14.5% | 22.3% | 1.8% |
| Date of Max Cover | 9/24 | 9/24 | 9/18 | 8/14 | 9/24 | 8/7 | 9/18 | 8/14 | 8/7 | 8/14 |
| Samples with Detected Anatoxins (%) | 75.0% *(9 of 12)* | 81.8% *(9 of 11)* | 76.9% *(10 of 13)* | 100% *(6 of 6)* | 63.6% *(7 of 11)* | 75.0% *(9 of 12)* | 87.5% *(7 of 8)* | 85.7% *(6 of 7)* | 100% *(7 of 7)* | 100% *(7 of 7)* |
| Mean Anatoxin Concentration (µg anatoxins $g^{-1}$ OM) | 8.8 | 10.0 | 9.0 | 17.6 | 3.6 | 18.1 | 16.0 | 6.5 | 15.6 | 3.9 |
| Max Anatoxin Concentration (µg anatoxins $g^{-1}$ OM) | 55.4 | 41.2 | 35.3 | 250.4 | 12.0 | 118.1 | 176.6 | 43.8 | 87.8 | 21.7 |
| Date of Max Anatoxin Concentration | 9/18 | 9/4 | 9/4 | 8/22 | 9/4 | 8/14 | 8/14 | 8/14 | 8/7 | 8/14 |

**Table S10.** Anatoxin concentrations for *Microcoleus* samples for all five reaches on the South Fork Eel River (SFE) sampled weekly in 2023. Numeric values are total anatoxin concentrations in µg anatoxins per g organic matter; "not present" indicates that no samples were taken as the taxa was not visibly present; "limited sample" indicates that not enough of the taxa was present to collect and analyze for anatoxin; "no reach visit" indicates that we did not sample that reach on that date; and "ND" indicates that a sample was taken but no anatoxins were detected.

| Reach | **SFE-Lower-1S** | **SFE-Lower-2** | **SFE-Lower-3** | **SFE-Lower-4** | **SFE-Upper-1S** |
|---|---|---|---|---|---|
| Taxa | *Microcoleus* | | | | |
| Visit 1 *(6/20)* | not present | limited sample material | not present | not present | no reach visit |
| Visit 2 *(6/25)* | not present | not present | not present | limited sample material | limited sample material |
| Visit 3 *(7/3)* | limited sample material | not present | ND | not present | limited sample material |
| Visit 4 *(7/10-7/11)* | ND | ND | ND | not present | ND |
| Visit 5 *(7/17)* | ND | not present | ND | not present | ND |
| Visit 6 *(7/24)* | ND | ND | 0.14 | limited sample material | ND |
| Visit 7 *(7/31)* | 0.32 | 0.32 | 0.41 | limited sample material | 0.19 |
| Visit 8 *(8/7)* | 0.53 | 7.5 | 2.8 | 2.7 | not present |
| Visit 9 *(8/14)* | 1.4 | 3.3 | 8.9 | limited sample material | ND |
| Visit 10 *(8/22)* | 5.6 | 30.9 | 9.7 | 250.4 | 2.2 |
| Visit 11 *(8/28)* | 7.2 | 34.9 | 25.9 | limited sample material | 9.5 |
| Visit 12 *(9/4)* | 28.0 | 41.2 | 35.3 | 5.8 | 12.0 |
| Visit 13 *(9/12)* | 17.2 | 16.1 | 27.4 | 2.7 | 6.7 |
| Visit 14 *(9/18)* | 55.4 | 10.8 | 17.3 | 1.5 | 10.5 |
| Visit 15 *(9/24)* | 16.8 | 5.3 | 7.4 | 0.17 | 9.4 |

**Table S11.** Anatoxin concentrations for *Anabaena/Cylindrospermum* samples for all five reaches on the South Fork Eel River (SFE) sampled weekly in 2023. Numeric values are total anatoxin concentrations in µg anatoxins per g organic matter; "not present" indicates that no samples were taken as the taxa was not visibly present; "limited sample" indicates that not enough of the taxa was present to collect and analyze for anatoxin; "no reach visit" indicates that we did not sample that reach on that date; and "ND" indicates that a sample was taken but no anatoxins were detected.

| Reach | **SFE-Lower-1S** | **SFE-Lower-2** | **SFE-Lower-3** | **SFE-Lower-4** | **SFE-Upper-1S** |
|---|---|---|---|---|---|
| Taxa | *Anabaena/Cylindrospermum* | | | | |
| Visit 1 *(6/20)* | not present | not present | not present | not present | no reach visit |
| Visit 2 *(6/25)* | not present | not present | not present | not present | not present |
| Visit 3 *(7/3)* | not present | not present | not present | not present | not present |
| Visit 4 *(7/10-7/11)* | ND | limited sample material | not present | 0.06 | not present |
| Visit 5 *(7/17)* | 0.04 | limited sample material | limited sample material | 0.04 | not present |
| Visit 6 *(7/24)* | 0.33 | 4.9 | 0.28 | 2.0 | 0.33 |
| Visit 7 *(7/31)* | 17.4 | 14.8 | 1.8 | 11.8 | 0.31 |
| Visit 8 *(8/7)* | 32.1 | 9.4 | 10.7 | 87.8 | 3.5 |
| Visit 9 *(8/14)* | 118.1 | 176.6 | 43.8 | 67.3 | 21.7 |
| Visit 10 *(8/22)* | 63.2 | 34.7 | 39.4 | 65.2 | 15.8 |
| Visit 11 *(8/28)* | 33.5 | limited sample material | 1.6 | limited sample material | not present |
| Visit 12 *(9/4)* | 6.5 | 0.09 | limited sample material | not present | not present |
| Visit 13 *(9/12)* | 0.09 | not present | not present | not present | not present |
| Visit 14 *(9/18)* | ND | ND | ND | limited sample material | 11.6 |
| Visit 15 *(9/24)* | ND | 0.04 | limited sample material | limited sample material | 1.7 |

**References**


Appling, A. P., R. O. Hall, C. B. Yackulic, and M. Arroita. 2018. Overcoming Equifinality: Leveraging Long Time Series for Stream Metabolism Estimation. Journal of Geophysical Research: Biogeosciences 123:624–645.

Ayers Associates. 1999. Geomorphic and Sediment Evaluation of the Klamath River, California, Below Iron Gate Dam. Fort Collins, CO.

Boyer, G. L. 2020. LCMS-SOP Determination of Microcystins in Water Samples by High Performance Liquid Chromatography (HPLC) with Single Quadrupole Mass Spectrometry (MS). protocols.io.

De Cicco, L. A., R. M. Hirsch, D. Lorenz, J. D. Watkins, and M. Johnson. 2024. dataRetrieval: R packages for discovering and retrieving water data available from Federal hydrologic web services. U.S. Geological Survey, Reston, VA.

Dewiz, J. 2021. . U.S. Geological Survey data release.

Emerson, K., R. C. Russo, R. E. Lund, and R. V. Thurston. 1975. Aqueous Ammonia Equilibrium Calculations: Effect of pH and Temperature. Journal of the Fisheries Research Board of Canada 32:2379–2383.

Foster, M. A., and H. M. Kelsey. 2012. Knickpoint and knickzone formation and propagation, South Fork Eel River, northern California. Geosphere 8:403–416.

Garcia, H. E., and L. I. Gordon. 1992. Oxygen solubility in seawater: Better fitting equations. Limnology and Oceanography 37:1307–1312.

Hall, R. O., and E. R. Hotchkiss. 2017. Stream Metabolism. Pages 219–233 Methods in Stream Ecology. Elsevier.

Leopold, L. B., and T. Maddock. 1953. The hydraulic geometry of stream channels and some physiographic implications. Page 64. Report, Washington, D.C.

NCRWQCB. 2024. Implementation of a Benthic Cyanobacteria Tiered Monitoring Program for Public Health Protection in Northern California Rivers. Freshwater Harmful Algal Bloom Monitoring and Response Program, North Coast Regional Water Quality Control Board, Santa Rosa, CA.

PME. 2021. miniDOT Logger User's Manual. PME, Vista, CA.

Raymond, P. A., C. J. Zappa, D. Butman, T. L. Bott, J. Potter, P. Mulholland, A. E. Laursen, W. H. McDowell, and D. Newbold. 2012. Scaling the gas transfer velocity and hydraulic geometry in streams and small rivers. Limnology and Oceanography: Fluids and Environments 2:41–53.

Rodell, M., P. R. Houser, U. Jambor, J. Gottschalck, K. Mitchell, C.-J. Meng, K. Arsenault, B. Cosgrove, J. Radakovich, M. Bosilovich, J. K. Entin, J. P. Walker, D. Lohmann, and D. Toll. 2004. The Global Land Data Assimilation System. Bulletin of the American Meteorological Society 85.

Sartory, D. P., and J. U. Grobbelaar. 1984. Extraction of chlorophyll a from freshwater phytoplankton for spectrophotometric analysis. Hydrobiologia 114:177–187.

Savoy, P., E. Bernhardt, L. Kirk, M. J. Cohen, and J. B. Heffernan. 2021. A seasonally dynamic model of light at the stream surface. Freshwater Science 40:286–301.

Stillwater Sciences. 2016. Sonoma County Aggregate Resources Management Plan: 2009-2014 Russian River Monitoring Results. Berkeley, CA.

Stroud Water Research Center. 2017. Model My Watershed.

U.S. EPA. 1993a. Method 365.1: Determination of Phosphorus by Semi-Automated Colorimetry. Cincinnati, OH.

U.S. EPA. 1993b. Method 350.1: Nitrogen, Ammonia (Colorimetric, Automated Phenate). Cincinnati, OH.

U.S. EPA. 1993c. Method 353.2: Determination of Nitrate-Nitrite Nitrogen by Automated Colorimetry. Cincinnati, OH.

U.S. EPA. 2015. Method 545: Determination of Cylindrospermopsin and Anatoxin-a in Drinking Water by Liquid Chromatography Electrospray Ionization Tandem Mass Spectrometry (LC/ESI-MS/MS). Cincinnati, OH.

U.S. Geological Survey. 2025. USGS Water Data for the Nation: U.S. Geological Survey National Water Information System database.

Zabrecky, J. M., T. A. Elliott, M. Hickey, H. Lei, R. S. Christova, G. Boyer, L. Genzoli, G. Johnson, and J. R. Blaszczak. 2025. Anatoxin concentrations, algal assemblages, and water quality data for the South Fork Eel, Salmon, and Russian Rivers in northern California, 2022-2023. Environmental Data Initiative. ver 3.

Zeileis, A., and G. Grothendieck. 2005. zoo: S3 Infrastructure for Regular and Irregular Time Series. Journal of Statistical Software 14:1–27.
